# BAcTrace a new tool for retrograde tracing of neuronal circuits

**DOI:** 10.1101/2020.01.24.918656

**Authors:** Sebastian Cachero, Marina Gkantia, Alexander S. Bates, Shahar Frechter, Laura Blackie, Amy McCarthy, Ben Sutcliffe, Alessio Strano, Yoshinori Aso, Gregory S.X.E. Jefferis

**Affiliations:** Neurobiology Division, MRC Laboratory of Molecular Biology, Cambridge, UK; Janelia Research Campus, Howard Hughes Medical Institute, Ashburn, United States; MRC London Institute of Medical Sciences, Imperial College London, UK; Cancer Research UK Manchester Institute, The University of Manchester, UK; Department of Cancer and Developmental Biology & Zayed Centre for Research into Rare Disease in Children, University College London, UK

## Abstract

New tools and techniques have enabled many key advances in our understanding of the brain. To elucidate circuit function, it is necessary to identify, record from and manipulate networks of connected neurons. Here we present BAcTrace (<u>B</u>otulinum <u>Ac</u>tivated <u>Tr</u>acer), the first fully genetically encoded, retrograde, transsynaptic labelling system. BAcTrace is based on *C. botulinum* neurotoxin A, Botox, which we have engineered to act as a Trojan horse that jumps retrogradely between neurons to activate an otherwise silent transcription factor. We validate BAcTrace at three connections in the *Drosophila* olfactory system and show that it enables electrophysiological recordings of connected neurons. Finally, in a challenging circuit with highly divergent connections, we used Electron Microscopy connectomics to show that BAcTrace correctly identifies 12 out of 16 connections.

## 1. Introduction

The development of genetic tools to elucidate connectivity and manipulate neurons and circuits has been key to advancing our understanding of how the brain works. Increasingly these tools are being used to study diseases of the nervous system and develop effective treatments [31, 57].

In the context of circuit research, the ability to identify and manipulate pre- or post-synaptic cells to neurons of interest is of crucial importance. For example, if genetic drivers are available for sensory neurons in the skin then one might want to label downstream, post-synaptic cells in the nerve cord. Conversely, when studying motor circuits, genetic drivers for motor neurons might be available and revealing upstream, pre-synaptic cells will be appropriate. Tools to label downstream neurons (e.g. for “walking” from sensory input towards motor-outputs as in the first example) are called anterograde, while retrograde tools reveal the input neurons to a given population.

*Drosophila melanogaster* is a key model organisms to study the genetic and circuit basis of animal behaviour (e.g. see [62, 15]). The fly has a rich behavioural repertoire encoded in a relatively small nervous system. This numerical simplicity is paired with extensive collections of genetic reagents, both to investigate gene function (e.g. mutant and RNAi collections) and to label and manipulate most neuronal classes (using the orthogonal expression systems Gal4, LexA, QF and their split versions [4, 45, 30, 58]). While these reagents offer excellent genetic access to neurons, until recently the fly lacked tools to map synaptic connections between neurons. This has recently changed with the development of electron microscopy methods to map connections in larval [38] and adult [63] brains. Furthermore, two genetically encoded systems for anterograde tracing: *trans*-Tango [54] and TRACT [22] have recently been developed. Despite these important additions to the experimental toolbox, a retrograde labelling system is still missing. Rabies virus and its modifications constitute the most notable examples of retrograde transsynaptic tools [31]. While in mice rabies has been used to great success, its applicability to flies is far from simple, both because of the experimental difficulties of delivering viruses via brain injections and because the virus neurotropism may not extend to flies.

We now introduce BAcTrace (<u>B</u>otulinum <u>Ac</u>tivated <u>Tr</u>acer), a fully genetically encoded retrograde tracing tool for *Drosophila*. We first established the system in tissue culture to demonstrate its viability. Then we implemented and refined BAcTrace in flies, showing that it can reveal the connectivity between olfactory Projection Neurons (PNs) and 3 different classes of presynaptic neurons: Olfactory Receptor Neurons, Kenyon Cells of the Mushroom Bodies and Lateral Horn Neurons. BAcTrace provides a powerful new way to test connectivity and manipulate components in circuits with high sensitivity and specificity.

## 2. Results

### 2.1. System design

Transsynaptic tools must induce labelling in connected neurons with high signal to noise ratio avoiding false positives. Labelling must depend on some form of communication between connected cells, either by transfer of material between them or through contact-based signal induction. Rabies virus falls within the first category: infected neurons make new viral particles which spread via synaptic contacts and once in the cytosol of connected neurons get amplified, strongly labelling them. *trans*-Tango and TRACT belong to the contact-based category; pre-synaptic neurons make a ligand that binds a receptor in post-synaptic partners triggering a conformational change that eventually activates a transcription factor.

To reveal neuronal connectivity in the fly brain, we designed BAcTrace, a novel transsynaptic labelling system which is fully genetically encoded and, in contrast to contact based systems, shares two key feature of rabies: 1. labelling is triggered by protein transfer between connected neurons and 2. this transfer is followed by a signal amplification step. We reasoned that this design could allow increases in sensitivity without compromising signal-to-noise ratio. At the core of our system sits a neuronal trojan horse, *Clostridium botulinum* neurotoxin A1 (BoNT/A). BoNT/A is well suited to synthetic biology because it is a modular protein (Fig. 1A and S1) with a well-studied mechanism of action (Fig. S2). BoNT/A is made as a single polypeptide that gets cleaved by proteases generating a Light Chain (LC) and a Heavy Chain (HC) (Fig. 1A). During intoxication in vertebrates, the receptor binding domain (RBD), located in the C-terminal half of the HC, enables enrichment on neuronal membranes by interacting with a neuronal specific lipid (polysialoganglioside GT1b). Upon neurotransmitter vesicle fusion the RBD gains access to the vesicle lumen and binds with high affinity to a second partner, synaptic protein SV2 [14]. When the vesicle is recycled and acidified, the translocation domain (TD) undergoes a conformational change injecting the LC across the vesicle membrane. The LC is a protease highly specific for the human protein SNAP25 (hSNAP25) and once in the cytosol it cleaves its target, preventing further neurotransmitter release [35, 44].

**Figure 1:**
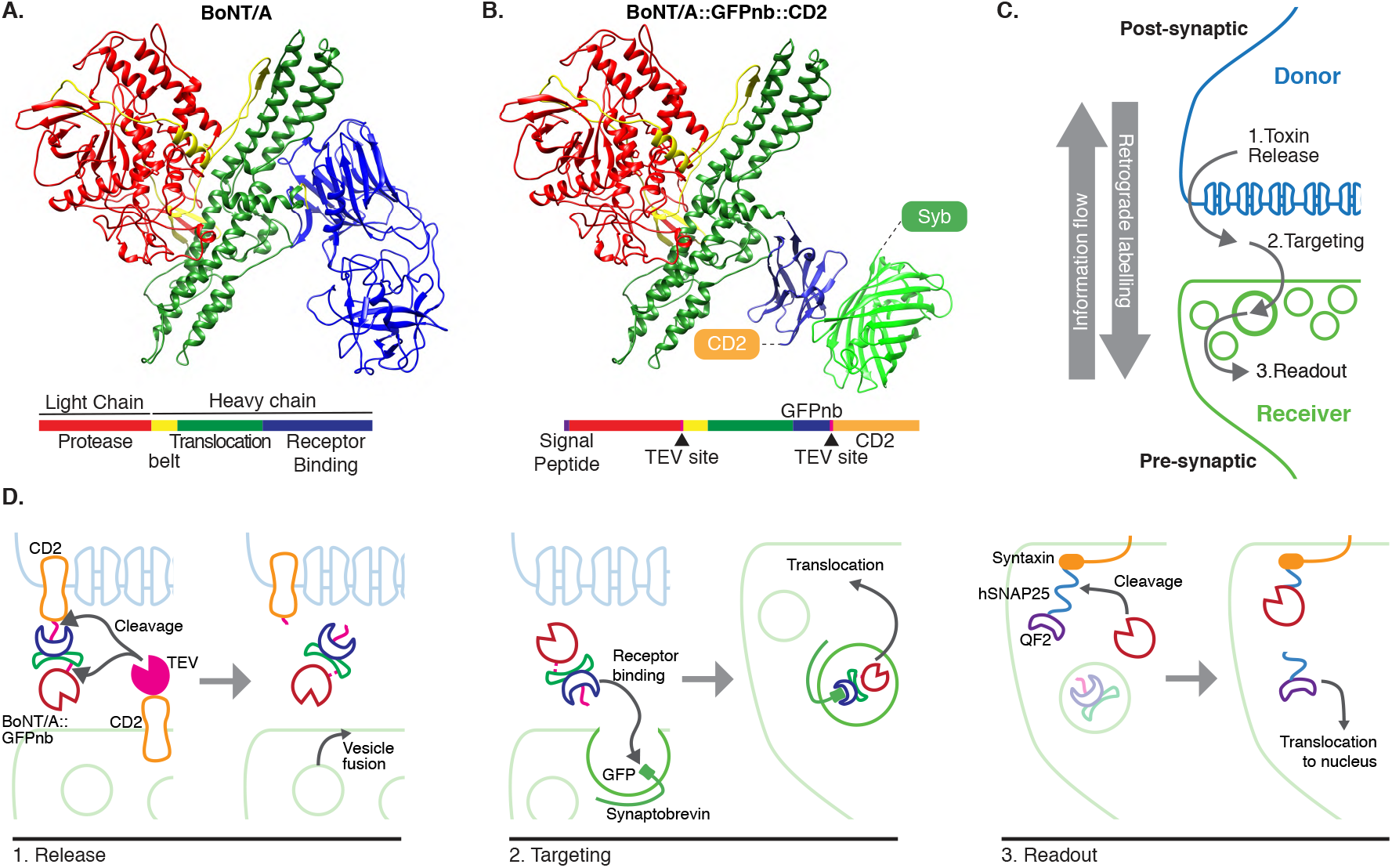
System design. (**A**) Molecular structure of BoNT/A (3BTA, [29]). Toxin domains are coloured red (protease), yellow (belt), green (Translocation, TD) and blue (Receptor Binding, RBD). (**B**) Retargeted BoNT/A: the RBD from (A) has been replaced by an anti-GFP nanobody (blue). Also shown in green is the bound GFP (3OGO, [28]). CD2 and Synaptobrevin targeting the toxin and the GFP to the plasma and neurotransmitter vesicle membranes are also indicated. (**C**) Schematic showing the three steps leading to the labelling. (**D**) Schematic of the proteins and molecular mechanisms mediating each step in (C). All colours are as in (B) except pink sections in BoNT/A::GFPnb::CD2 which indicate TEV cleavage sites engineered into the toxin for its membrane release by TEV.

In BAcTrace BoNT/A is expressed in post-synaptic “Donor” neurons and transferred, similarly to rabies, to connected pre-synaptic “Receiver” cells to trigger expression of an effector gene. By virtue of *Botulinum* neurotoxin intrinsic mechanism of action our system works retrogradely, opposite to the flow of information. BAcTrace can be broken down into three steps (Fig. 1C,D): 1. Toxin release from Donor neurons into the synaptic cleft, 2. Targeting to neurotransmitter vesicles and escape into the cytosol and 3. Readout in the form of activating effector gene expression.

During step 1 (Fig. 1D) BoNT/A is made in Donor neurons attached to the extracellular portion of a transmembrane protein (CD2) and the Tobacco Etch Protease (TEV), a protease previously used in flies [40], is made in Receiver neurons attached to the extracellular portion of a second transmembrane protein. At synapses both proteins interact and TEV cleaves two recognition sites, releasing the toxin from the membrane, and allowing separation of the LC and HC after translocation (Fig. 1B,D). Because flies lack BoNT/A vertebrate receptor SV2, we switched the RBD for a single-chain anti-GFP nanobody (GFPnb) [28] to create BoNT/A::GFPnb::CD2 (Fig. 1B and S3). This modified toxin is targeted to neurotransmitter vesicles of Receiver neurons that express a Synaptobrevin::GFP fusion, oriented with GFP inside the vesicle. This re-targeting also makes the toxin safe, since it can no longer bind to vertebrate neurons. At the end of step 2, the toxin LC protease enters the pre-synaptic terminal.

As a readout we made a toxin sensor by linking a QF2 transcription factor [46] to a *Drosophila* Syntaxin molecule via amino acids 141-206 of hSNAP25 [60] (QF2::hSNAP25::Syx). Syntaxin targets the sensor to the synaptic membrane, so that QF2 cannot activate transcription. However cytosolic LC will release QF2, triggering expression of a QUAS-reporter transgene. BoNT/A LC is highly specific and it has been shown to not cleave *Drosophila* SNAP25 (dSNAP25) [60], so toxin expression should not be harmful to flies.

### 2.2. BAcTrace is active in Drosophila cells

We started by testing, step by step, the feasibility of our approach in *Drosophila* tissue culture S2 cells. To establish that the retargeted BoNT/A::GFPnb-GFP receptor pair could mediate toxin transfer we expressed and purified BoNT/A::GFPnb from bacteria. We added the purified toxin to S2 cells rendered sensitive by expression of a hTfR::GFP receptor which cycles between the plasma membrane and a low pH compartment ([20] and supplemental text 7.1.1). The cells were also transfected with a FLAG tagged hSNAP25, a simple toxin sensor (see Fig. 2A). Cleavage of Flag::hSNAP25 was measured by western blot as a small shift of 0.9kDa (9 amino acids). Fig. 2B shows that all tested toxin concentrations induced efficient cleavage (see also Fig. S4A and supplemental text 7.1.2). Furthermore, hTfR::GFP was strictly required for BoNT/A::GFPnb to induce hSNAP25 cleavage (‘No receptor’ lanes in Fig. 2B).

**Figure 2:**
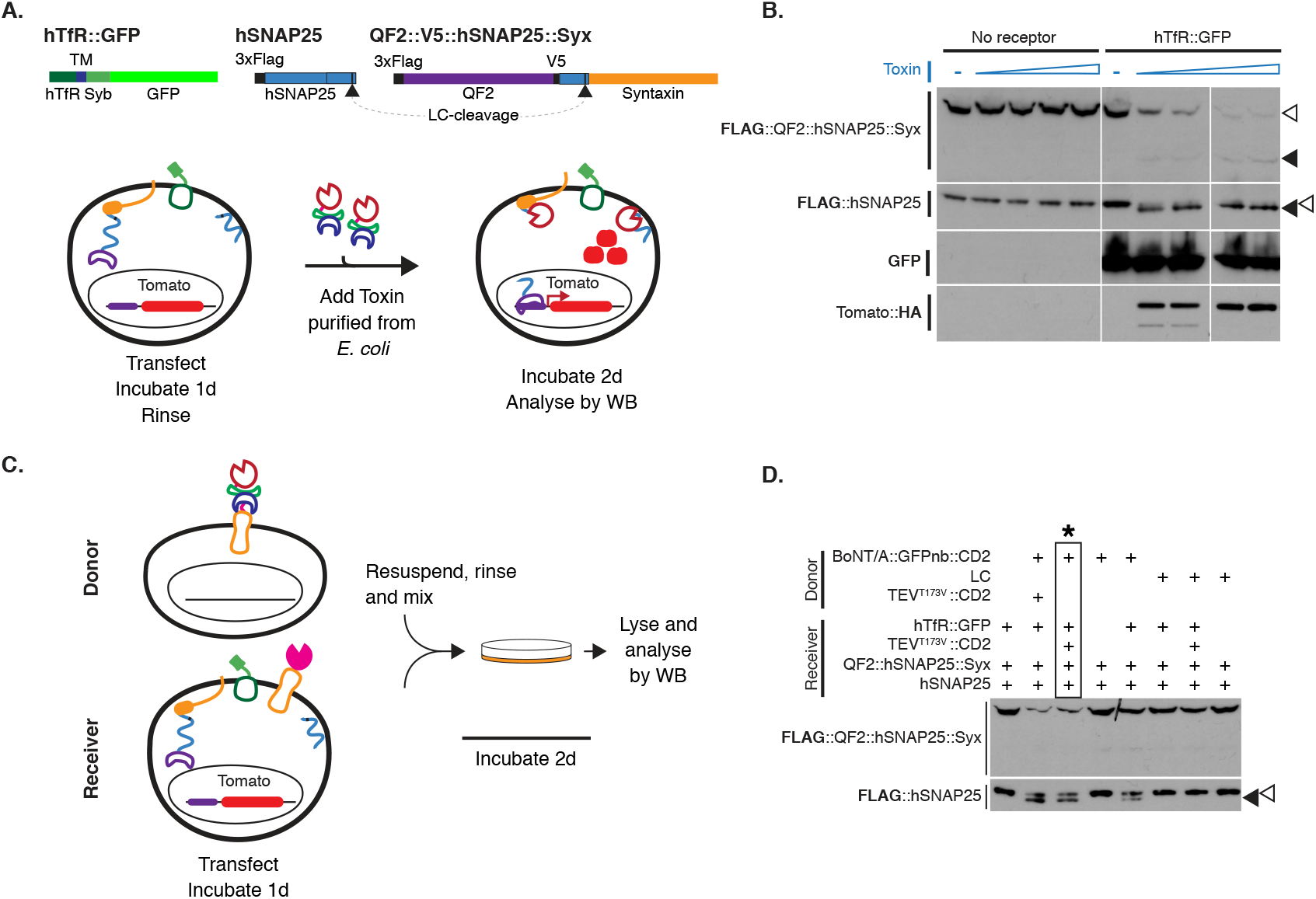
Testing BAcTrace in tissue culture. (**A**) Schematic of transgenes and tissue culture experimental outline used to test *E. coli* produced toxin. (**B**) Western blot analysis of S2 cell extracts after transfection and incubation with toxin made in bacteria. Empty arrowheads indicate un-cleaved and solid arrowheads cleaved toxin sensors. Toxin concentrations (nM) in each lane are: 0, 1.7, 3.4, 6.9, 14, 0, 1.7, 3.4, 6.9 and 14. Smaller band in Tomato::HA panels is a degradation C-terminal fragment. (**C**) Schematic of cell mixing experiments to test toxin produced in insect cells. (**D**) Western blot analysis of S2 cell extracts from cell mixing experiments. Conditions marked with * correspond to those from panel (C). Epitopes detected by antibodies are indicated in bold in (B) and (D).

We also tested the sensor to be used in flies. QF2::hSNAP25::Syx can indeed be cleaved by BoNT/A::GFPnb (Fig. 2B) and the released QF2 induces Tomato expression, confirming that function of QF2 as a transcription factor is not inhibited by the 62 amino acids of hSNAP25 left on its C-terminus. Unexpectedly, the QF2::hSNAP25::Syx cleavage product accumulates less than that of hSNAP25. This may be due to higher stability of the cleaved hSNAP25, which remains on the membrane, compared to the cleaved QF2 which becomes cytosolic.

Both TEV and BoNT/A are cytosolic proteins, usually made in a host plant or in the *C. botulinum* bacterium, respectively. For this reason it was important to verify the activity of these proteins when produced as membrane fusions exposed on the outside of *Drosophila* cells. We found that our extracellular TEV construct was inactive until we mutated the amino-acid Threonine in position 173 to Valine (TEV^T173V^) removing a predicted glycosylation site (Fig. S4C-G and supplemental text 7.1.3). Next, we tested the function of our plasma membrane-targeted BoNT/A using a cell mixing experiment. We transfected two cell populations: 1. Donor cells with either membrane targeted BoNT/A::GFPnb::CD2 and 2. Receiver cells with receptor and sensors (Fig. 2C). After incubating one day to allow plasmid expression, Donor and Receiver cells were rinsed several times to remove traces of transfection reagents and mixed together. Two days later we could detect hSNAP25 cleavage, confirming that our BoNT/A::GFPnb::CD2 fusion protein was able to pass from the Donor cell to the cytoplasmic compartment of the Receiver cell to cleave the hSNAP25 based sensor proteins (Fig. 2D). Interestingly we found that TEV^T173V^::CD2 was not essential for this transfer, perhaps due to TEV independent cleavage and release of BoNT/A::GFPnb from the Donor cell membrane. Critically, just as for the experiments with bacterial BoNT/A::GFPnb (Fig. 2B), we observed a strict requirement for TfR::GFP in Receiver cells.

### 2.3. BAcTrace works as a transsynaptic system in flies

Our cell culture results established essential principles for BAcTrace: that *Drosophila* cells can make functional BoNT/A::GFPnb::CD2 which jumps from Donor to Receiver cells, that it can enter the Receiver cell cytoplasm, cleave a hSNAP25 target site, releasing QF2 to drive reporter gene expression. We therefore made transgenic flies to test the system *in vivo*. Donor neuron components were encoded in UAS vectors, driven by Gal4, with Receiver neuron components driven by the LexA system. For transsynaptic experiments, flies containing all BAcTrace components were crossed to flies with Gal4 and LexA promoter fusions (Fig. 3A); for some experiments the LexA driver was also placed in the BAcTrace fly.

**Figure 3:**
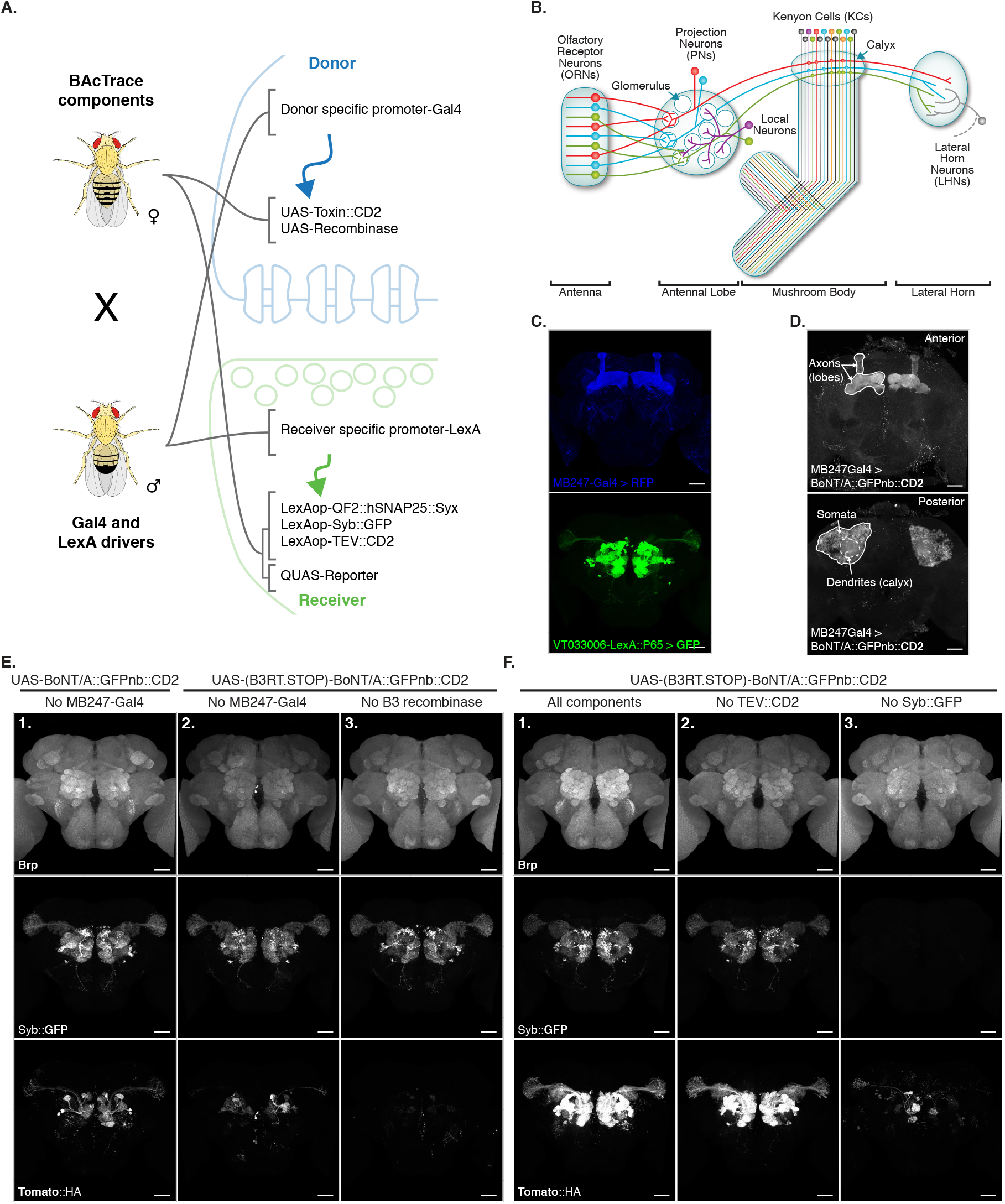
BAcTrace *in vivo*. (**A**) Schematic of the genetic strategy for expressing BAcTrace components. The female fly has all UAS, LexAop and QUAS components while the male has a Gal4 driver defining the Donor cells and a LexA driver defining the Receiver cells. (**B**) Schematic of the fly olfactory system; adapted from [6]. (**C**) Reporters showing the expression patterns of MB247-Gal4 (blue) and VT033006-LexA::P65 (green). (**D**) MB247-Gal4 driving expression of BoNT/A::GFPnb::CD2 in the mushroom bodies. (**E**) BAcTrace negative controls lacking toxin expression. (**E1**) No Gal4 driver (**E2**) Same as E1 (no Gal4) but a transcriptional stop has been placed in front of the toxin. The UAS-B3 transgene encoding the DNA recombinase that can remove the stop cassette is present. (**E3**) Stop cassette present, same as E2, MB247-Gal4 present and UAS-B3 absent. (**F**) BAcTrace experiments. (**F1**) All components present as described in Fig. 1D. (**F2**) Same as F1 except TEV transgene is absent. (**F3**) Same as F1 except the Syb::GFP receptor is absent. (C), (E) and (F) are full and (D) partial maximum intensity projections of confocal stacks. Epitopes detected by antibodies are indicated in bold in (C-F). All animals are 3-4 days old. Genotypes for each panel in Table S15. Scale bars 30µm.

We chose the fly olfactory system for initial testing because of the wealth of genetic reagents and its well-studied connectivity (Fig. 3B). Briefly, *Drosophila* has 50 types of peripheral Olfactory Receptor Neurons (ORNs). ORNs of each type (e.g. red neurons in the antenna of Fig. 3B) express one of 50 different olfactory receptor genes, conferring responses to a specific set of odours, and a general co-receptor, Odorant receptor Co-Receptor (Orco). ORNs expressing the same receptor relay information to one of 50 glomeruli in the brain’s Antennal Lobe (AL). In each glomerulus the axons of 20-100 ORNs make strong connections (e.g. rv1215 synapses per PN in glomerulus DM6) onto the dendrites of 1-8 Projection Neurons (PNs); these in turn make a modest number of reciprocal synapses (e.g.rv40 synapses per PN in glomerulus DM6) onto ORNs. PN axons project onto Kenyon Cells (KCs) in the calyx of the Mushroom Bodies (MB, required for associative learning) and to the Lateral Horn (LH, required for innate behaviours) [33]. PNs make on average 7 large boutons in the calyx, each containing 40 pre-synaptic sites which are contacted by 220 small postsynaptic elements from 11 KCs [5, 56], making these large synapses good candidates for initial testing. For our tracing experiments it is important to note that PNs connect to KCs randomly [56, 36, 7]; therefore expressing toxin even in a small number of KCs should result in all PNs labelled.

We expressed Donor components in KCs using the MB247-Gal4 driver and Receiver components in PNs using the broad LexA driver VT033006-LexA::P65 (Fig. 3C, see also Fig. S5 for the identity of PNs present in the line). We confirmed that BoNT/A::GFPnb::CD2 can be expressed and trafficked throughout KCs, detecting expression in soma, dendrites and axons (Fig. 3D).

We first established the background signal of the system *in vivo* by bringing together all components except the driver used to express toxin in the Donor cells. We found low-frequency, stochastic labelling of PNs with the QUAS-Tomato::HA reporter (Fig. 3E1). Our system can amplify very low levels of the LC protease within Receiver cells, so this signal is likely due to weak, Gal4-independent toxin expression in PNs (BAcTrace working in *cis* instead of *trans*). To reduce this background, we added a transcriptional stop cassette in front of BoNT/A::GFPnb::CD2 which can be removed by the B3 DNA recombinase [37]. This modification implements a step filter: only cells that express UAS-B3 above a threshold can activate the BoNT/A::GFPnb::CD2 transgene. Fig. 3E2 shows that the stop cassette reduces background in the absence of Gal4. Furthermore, in the absence of the B3 recombinase transgene (Fig. 3E3) there is no stochastic PN labelling, even in the presence of MB247-Gal4, indicating that the remaining stochastic background in Fig. 3E2 is due to Gal4 independent expression of B3. It is worth mentioning that this Gal4 independent expression might happen during development. Besides the stochastic background we found a much lower, non-stochastic background which is not ameliorated by the stop cassette (Fig. 3E3). We mapped the source of this signal to the small V5 tag present in the sensor (Fig. S6 and supplemental text 7.2.2). Except when indicated, in all further experiments we used a modified sensor without V5.

After minimising the background, we performed the first *in vivo* transsynaptic experiment. We found that toxin expression in the MB induced strong labelling of most PNs in the VT033006 line (compare Fig. 3E2 and F1). Just as for S2 cells, we found strong labelling even in the absence of TEV (compare Fig. 3F1 and F2). Although further experiments confirmed a small improvement in efficiency with mutant TEV^T173V^ (Fig. S7 and supplemental text 7.2.3), given that it is not essential for labelling we decided to omit it from the remaining experiments. Critically we found an absolute requirement for the Syb::GFP receptor for *in vivo* transfer of toxin (Fig. 3F3), paralleling the requirement of the hTfR::GFP receptor in tissue culture. Although we observed low level stochastic labelling in the absence of receptor in Receiver PN cells (Fig. 3F3), this was at the same level as when we omitted the driver from Donor cells (Fig. 3E2); this strongly suggests that background labelling again originates from low-level toxin expression in Receiver cells rather than receptor independent transfer from Donor cells. The strict requirement for a synaptic component (i.e. Syb::GFP) for active labelling is very significant since it should select for synaptic rather than non-synaptic cell contacts.

Our initial test used the MB247-Gal4 to express toxin in all MB KCs. However the MB contains 3 main types of KCs (i.e. α/β, α’/β’ and γ) that can be divided into 7 discrete subtypes by split Gal4 drivers [2]. Although the mushroom body as a whole receives random input from PNs [7], it is not known whether there are input biases onto different KC types. We found that expressing toxin in different KC types labelled all presynaptic PNs, but did do so with different efficiencies: α’/β’ > α/β *≈* γ (Fig. S8 and supplemental text 7.2.4). This suggests biases in PN input connectivity to different KC subtypes, an observation that should soon be possible to corroborate with Electron Microscopy (EM) connectomics information [50].

### 2.4. BAcTrace expression in Olfactory Receptor Neurons labels connected PNs

Having shown that BAcTrace can label the strong, all-to-all PN->KC synapses, we next examined its labelling specificity using an experimental configuration where only a subset of Receiver cells are connected to toxin-expressing Donor cells. We selected the reciprocal synapses from PNs to ORNs (see Fig. 3B) for two reasons: 1. there are very specific Gal4 driver lines available to drive BAcTrace expression in ORN subtypes [10, 17] and 2. while ORN axons make very strong connections to PN dendrites, EM connectomics data identified a moderate strength reciprocal connection between these two cell types [47]. For example, PNs from the DM6 glomerulus make rv40 such reciprocal synapses onto their ORNs (Fig. 4A)[55].

**Figure 4:**
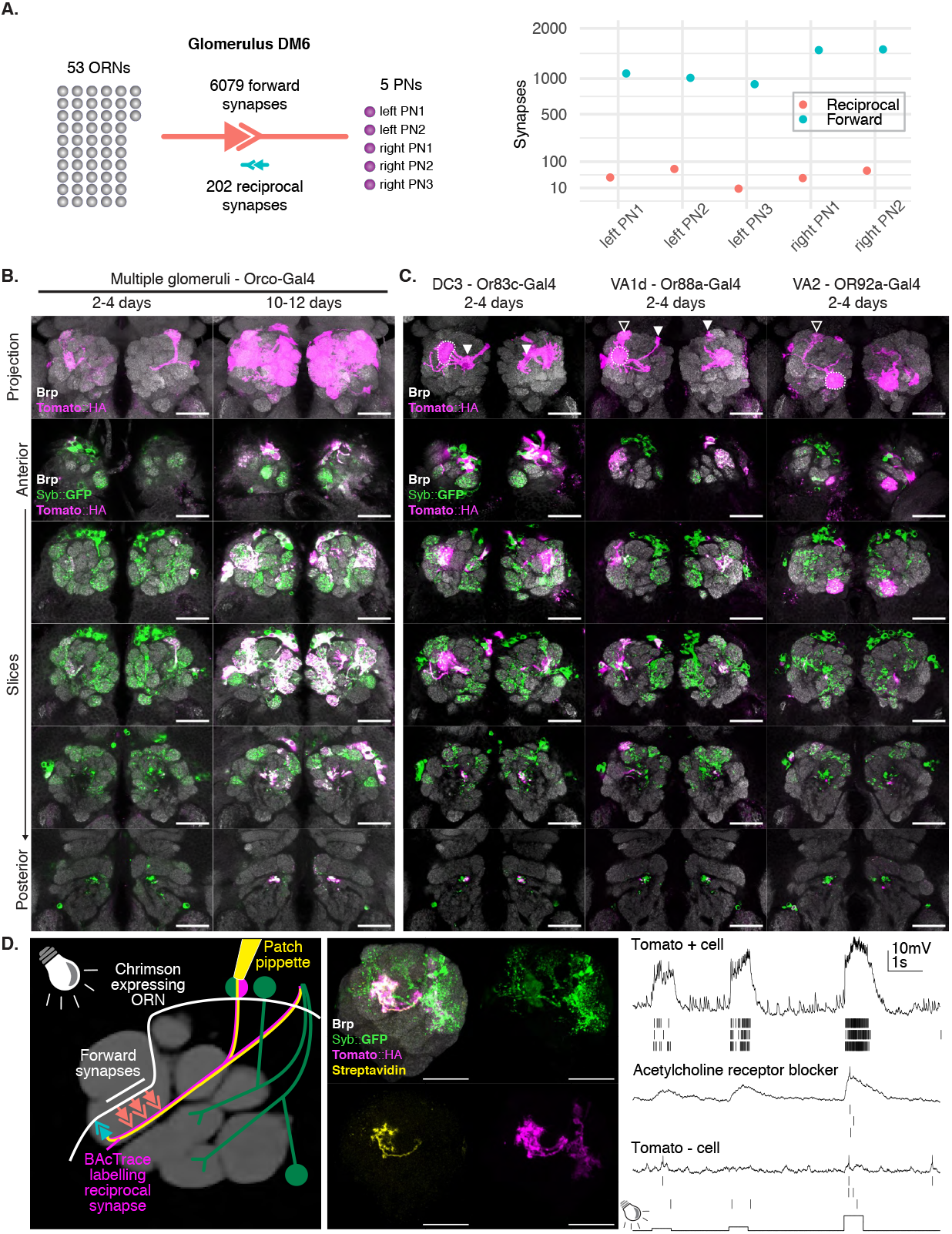
BAcTrace expression in ORNs labels connected PNs. (**A**) Summary of ORN->PN forward and reciprocal connectivity [55]. Left: aggregated numbers of forward and reciprocal synapses between PNs and ORNs in the DM6 glomerulus. Right: plot showing forward and reciprocal synapses per PN. y scale: sqrt (**B**) BAcTrace expression in Orco ORNs induce time dependent labelling in PNs. (**C**) BAcTrace expression in specific ORNs induces labelling in connected PNs (dotted lines). Occasionally labelling is also present in PNs from neighbouring glomeruli and less frequently non-neighbouring ones. Solid white arrowheads indicate neuronal soma and empty white arrowheads indicate glomerulus not innervated by the ORNs. (**D**) Whole cell patch clamp recording of a BAcTrace labelled PN. Left: Schematic showing experimental set up: BAcTrace labelled DC3 PNs (magenta) and Chrimson expressing Donor ORNs (white). Recorded cell is filled and labelled with biocytin-streptavidin (yellow). Middle: Maximum intensity projection of a confocal stack showing the recorded DC3 PN. Right: Representative voltage trace and spikes extracted from three light presentations. First spike row corresponds to shown voltage trace. Top: Recording from the cell shown in middle panel. Middle: same as top but with the addition of the nicotinic acetylcholine receptor blocker mecamylamine (200µM). Bottom: Recording from a GFP+ but Tomato-neuron showing minimal response to light. Animal in (D) is 14 days old. Epitopes detected by antibodies are indicated in bold in B, C and D. Genotypes for each panel in Table S15. Scale bars 30µm.

Given that we expected the reciprocal synapses from the ORN->PN connection to be relatively weak, we began by expressing toxin in the majority of ORNs using Orco-Gal4 (Fig. 4B). In 2-4 days old animals we detected little labelling above background. However, at nine to ten days after eclosion we found consistent labelling in most glomeruli covered by the VT033006-LexA::P65 line (see Fig. 4B). Quantifying BAcTrace labelling over time may therefore be used to characterise synapses of different strengths.

Next we expressed toxin in single ORN types and assessed their ability to label their cognate PNs (Fig. 4C and Fig. S11). In most cases strong labelling was detected in animals as young as 2 days old, e.g. Or83c, Or88a, Or92a, Or65a and Or98a while in other cases specific labelling took longer, e.g 10-13 days for Or47a. While these differences may be due to cell-type specific differences in the number of ORN->PN reciprocal synapses, they could also be the result of expression level variability of BAcTrace components in the different cell types (i.e. detection system components in PNs or toxin in ORNs).

In some cases BAcTrace also labelled PNs targeting glomeruli not innervated by Donor cells (+ in Fig. 4C and Fig. S11). This could be artifactual background labelling, as characterised before (Fig. 3E2), or might be triggered by non-synaptic contacts from the ORN axons as they traverse the AL to reach their target glomerulus. However the labelling could also be due to unexpected, synaptic contacts; for instance some ORNs and PNs have small processes in neighbouring glomeruli. Furthermore, VT033006 includes some multi-glomerular PNs which when labelled in one glomerulus would show signal in several others. Despite these few instances of unexpected labelling, BAcTrace quickly and robustly labelled PNs when toxin was expressed in their connected ORNs.

Viral transsynaptic tools such as rabies have the important caveat of being toxic to neurons. While the only know vertebrate target of BoNT/A, hSNAP25, is not cleaved in flies, we cannot rule out the existence of other targets or cytotoxic effects. To assess these possibilities, we used BAcTrace to label DC3 PNs while expressing the light gated cation channel CsChrimson in Donor, connected Or83c ORNs [25](Fig. 4D). We stimulated the ORNs using light while performing electrophysiological recordings from the Tomato labelled PNs in 14 days old flies (we obtained similar results on 5 days old animals, not shown). If BoNT/A::GFPnb::CD2 was toxic to neurons or detrimental to synaptic transmission then we would expect PNs would fail to respond. We found the opposite to be true, graded light stimulation induced increasing spiking responses in PNs indicating that ORNs are able to release neurotransmitter and stimulate connected PNs. These responses could be partially blocked by the nicotinic acetylcholine receptor blocker mecamylamine, indicating the responses are due to synaptic transmission from ORNs. Furthermore, light responses were absent from Tomato negative control neurons. Lastly, from a technical point of view, this experiment shows that BAcTrace label is strong enough for guiding cell patching using a regular fluorescent reporter.

### 2.5. Mapping connections in the Lateral Horn

For the last set of experiments we moved to the Lateral Horn (LH) where connectivity is less well understood and projections are more divergent. In the LH, each olfactory PN type has many post-synaptic Lateral Horn Neuron (LHN) partners with whom it shares fewer synapses than in our previous experiments (Fig. 3B). We expressed Receiver components in PNs using the VT033006-LexA::P65 and toxin in a panel of 8 Donor LHN types (Fig. S12A) [13]. All 8 Donors induced labelling of Receiver PNs (Fig. S12B). Consistent with our expectations of weaker connectivity from individual Receiver cells, labelling was only observed in older animals (16-18d but not 2-3d, not shown).

Unlike our previous experiments, there were significant differences in Receiver PN labelling across animals having the same Donor LHNs and even among left/right sides of the same brain (e.g. Fig. S13). To better understand these results we carried out a careful quantification, focussing on 2 LHN types for which EM connectivity data is available: PD2a1/b1 [12] and AV1a1 [23] (Fig. 5A) as well as two types for which more limited information is available: PV5c1 and AV6a1 (Fig. S14A). Fig. 5B shows a schematic of the results presented in Fig. 5C. PD2a1/b1 and AV1a1 target the dorsal and ventral LH respectively, with very little spatial overlap between them (Fig. 5A). Each Donor LHN cell type labelled Receiver PNs targeting different glomeruli; for example, VM3 and DM3 are labelled only in PD2a1/b1 while VM4 and VA1d are only labelled in AV1a1. Labelling differences can also be seen in the LH neuropile where the Receiver signal (Halo) has minimal overlap between the lines (compare LH panels in Fig. 5C). Furthermore, in the LH there is close apposition between toxin and Halo signals supporting the idea that labelling is induced by transsynaptic, retrograde transfer.

**Figure 5:**
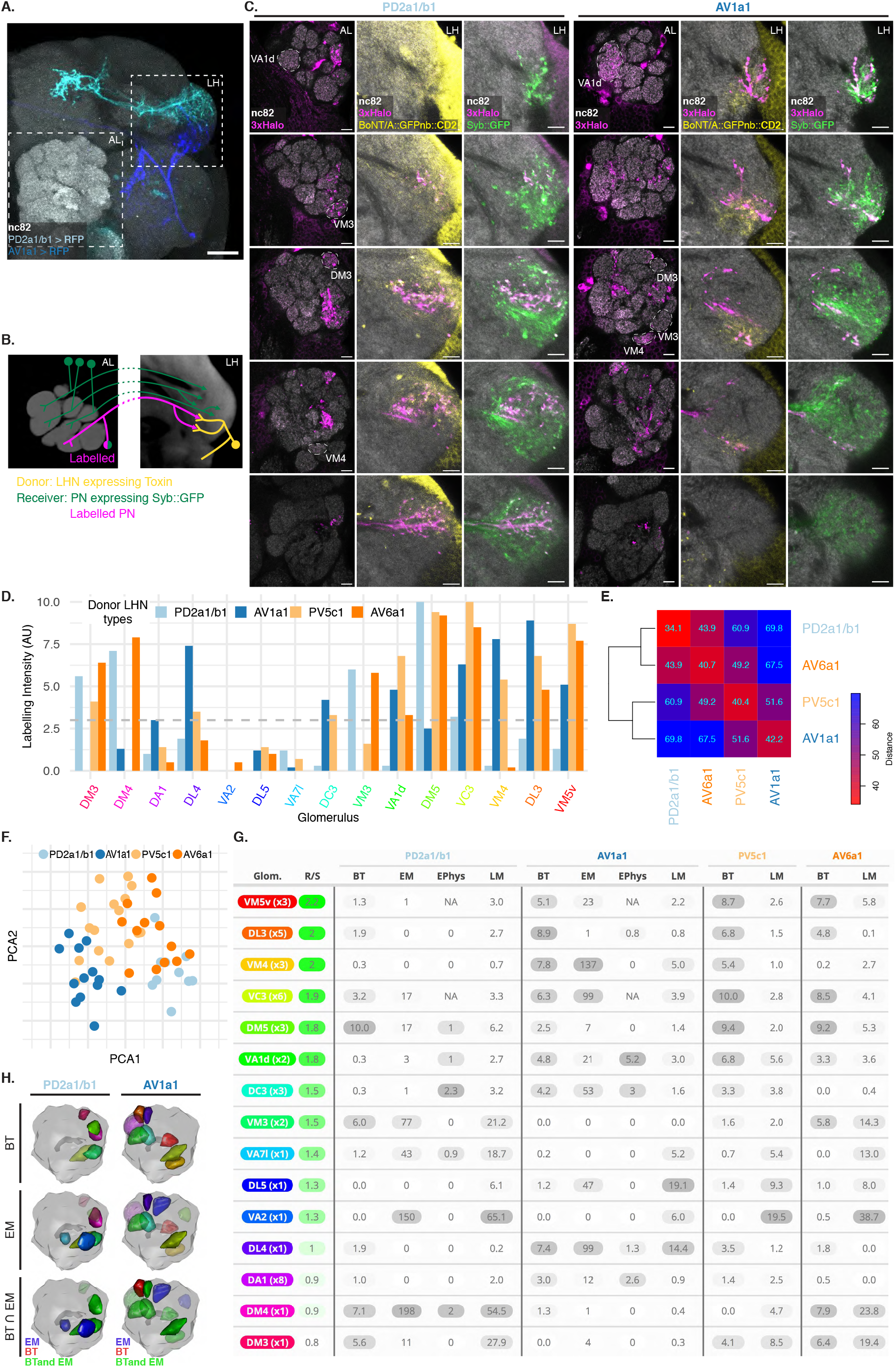
BAcTrace can reveal connections between PNs and LHNs. (**A**) Split Gal4 lines LH989 and LH1983 drive GFP expression in LHN cell types PD2a1/b1 and AV1a1, respectively. (**B**) Schematic of the Antennal Lobe (AL, left) and Lateral Horn (LH, right) depicting the results shown in (C). (**C**) Single slices from a representative AL and corresponding LH showing PNs labelled by expression of toxin in PD2a1/b1 and AV1a1. The BAcTrace reporter Halo is shown in the AL; Halo, Syb::GFP and Toxin are shown in the LH. CD2 stainings (labelling the toxin) have high background outside the neuropile. Examples of glomeruli differentially labelled between the 2 lines (VA1d, VM3, DM3 and VM4) are indicated. (**D**) Average labelling scoring per glomerulus for PD2a1/b1 (n=13 ALs), AV1a1 (n=10 ALs), PV5c1(n=18 ALs) and AV6a1 (n=13 ALs). Grey dotted line indicates average scoring of 3 (weak). (**E**) Heatmap of the Manhattan distance between the labelling scorings for each LHN type. (**F**) Principal Component Analysis for the vectors describing scoring results. (**G**) Summary of published data for the 4 LHN cell types. R/S: average expression level for <u>R</u>eceptor and <u>S</u>ensor (see Fig. S5). BT: <u>B</u>Ac<u>T</u>race results as quantified in (C). EM: <u>E</u>lectron <u>M</u>icroscopy connectivity data. EPhys: Electrophysiological recordings from LHNs during opto-stimulation of PNs [24]. LM: <u>L</u>ight <u>M</u>icroscopy overlap of single cell PNs and LHNs [8] and methods. (**H**) 3D renderings of ALs including BT: glomeruli with signal > 3, EM: glomeurli with more than 10 synapses and BT∩EM: meeting both conditions. Glomeruli colouring in BT and EM same as in (D); in BT∩EM green satisfy BT and EM criteria, blue EM only and green BT only. Epitopes detected by antibodies are indicated in bold in A and C. Genotypes for each panel in Table S15. Scale bars (A) 50µm and (C) 10µm.

To quantify these results, we first identified the 15 glomeruli with the highest expression level of Receiver components (Fig. S5C), reasoning that weaker glomeruli would be less efficiently labelled. We then scored glomerular labelling using a simple rating system (strong=10, weak=3, no labelling=0). We mixed samples within a single batch and the annotator was blind to genotype until the end of the process when we used toxin staining to identify LHN Donor cell types for each sample. Stacks are available online for readers to examine. The results are summarised per glomerulus in Fig. 5D. When comparing PD2a1/b1 and AV1a1, 12 out of 15 glomeruli were labelled by at least one of the two Donor cell types; 11 of those 12 were labelled with different strengths, suggesting differences in connectivity. We quantified these differences by computing the average distance with and across LHN cell types between the 15 dimensional annotation vectors for each sample (1 dimension per glomerulus, see methods). As expected, the distance was always lowest for comparisons within an LHN cell-type, indicating similar patterns of labelling. For the across cell-type comparisons, PD2a1/b1 and AV6a1 shared the most similar connectivity, differing from AV1a1 and PV5c1 (which clustered together, Fig. 5E).

We next asked: is there enough information within single labelled ALs to identify the Donor cell type of each specimen? We addressed this using principal component analysis (PCA) on the labelling annotations for all ALs. We found the first principal component efficiently separated individual PD2a1/b1 and Av1a1 labelled ALs while the second axis improved separation of PV5c1 and AV6a1(Fig. 5F). The labelling therefore contains enough information at the single specimen level to identify connectivity differences between different LHN types, especially those with more dissimilar connections.

Having established that BAcTrace show consistent differences in PN labelling when triggered by different Donor LHNs, we compared these results with previously published data. In Fig. 5G we summarise BAcTrace (BT), EM [12, 23], electrophysiology (EPhys)[24] and Light Microscopy (LM) overlap (see methods) data for PNs and LHNs. The comparison with PD2a1/b1 EM data is particularly informative as there is unequivocal correspondence between the neurons in the split GAL4 line used in our experiments and cells reconstructed by EM. For AV1a1 the correspondence is incomplete as the Gal4 line expresses strongly in 2-3 and more weakly in 6-10 neurons, all of which are likely to share similar morphologies. In the EM volume only 6 AV1a1 candidates have been identified and traced to varying levels of completion. Besides neuron correspondence, the comparison to EM data rely on the assumption of invariance in connectivity between animals; so far these neurons have only been traced in one brain and to what extent stereotypy will hold true for LHN-PN connections across animals remains to be tested.

We found good correspondence between BAcTrace, EM and LM data. For PD2a1/b1 neurons, 5 out of the 7 PN types that provide more than 10 synapses in the EM volume showed significant (>3) labelling in BAcTrace experiments (see also Fig. 5H). In the case of AV1a1, 7 out of 9 PN types with more than 10 EM synapses induced labelling in BAcTrace experiments. Unexpectedly, toxin expression in AV1a1 neurons additionally induced strong labelling in DL3 PNs (whereas EM tracing identified just one AV1a-DL3 synapse). While this might represent a false positive for BAcTrace, it is intriguing that there is considerable overlap between AV1a1and DL3 PNs as measured by light microscopy. Furthermore electrophysiological recordings [24] show a response in AV1a1induced by DL3 PN activation; this raises the possibility that DL3 PNs may connect to some of the AV1a1neurons weakly labelled by the L1983 line that have not yet been characterised by EM.

Lastly, assuming animal-to-animal stereotypy, the variability of labelling strength in our data indicates that BAcTrace efficiency is not only a function of synapse number. For instance, in PD2a1/b1 experiments DM3 PNs were efficiently labelled by just 11 synapses while 43 synapses induced very poor labelling in VA7l, despite VA7l PN expressing sensor and receptor more strongly than VM2 PNs. Future experiments and community feedback will help identify the different factors affecting labelling efficiency.

## 3. Discussion

In this study we present the design and implementation of BAcTrace and experiments showing its *in vivo* performance. BAcTrace is, to our knowledge, the first modular, fully genetically encoded retrograde labelling system as well as the first application of *C. botulinum* neurotoxin as a circuit tracer. In contrast to anterograde systems which identify ‘the next neuron’ in a circuit, BAcTrace reveals neuronal inputs. This is essential for mapping circuits ‘backwards’, starting in the motor rather than sensory periphery, and enables studies of neuronal integration throughout the brain. Our implementation of BAcTrace in *Drosophila* is therefore highly complementary to the recent *trans*-Tango [54] and TRACT [22] systems for anterograde labelling.

Exploiting the detailed connectivity information that exists for the fly’s olfactory system [55, 5, 33, 23, 18, 12, 63] we explored three important and related questions about BAcTrace: First, does labelling occur only in the retrograde direction? Second, is labelling specific? Third, how many synapses are required? Clear support for the retrograde direction of labelling comes from the PN->LHN experiments; comprehensive tracing of PD2a1/b1 and AV1a1 neurons in a whole brain EM volume failed to identify a single reciprocal LHN->PN synapse (Fig. S14D). This implies that the BAcTrace labelling observed must be retrograde. BAcTrace is specific since we only labelled the correct PNs when toxin was expressed in ORNs. This was confirmed by the match between our PN->LHN results and EM connectivity data. Regarding BAcTrace’s sensitivity, in our experiments we could detect labelling in connections ranging from 10 synapses (PN->LHN) to >200 synapses (PN->KC). Does this range encompass functional connections? While there is no single answer for how many synapses constitute a ‘functional connection’, recent work has found that seemingly low connection strengths, in the region of rv2% of a neuron’s total post-synaptic budget, can be functional [12, 16, 38, 48]. In the case of PD2a1/b1 and AV1a1neurons used in this study, the average number of post-synapses per neuron is 673, implying a 2% threshold value of around 12 synapses (Table S16). While these numbers are tentative, they indicate that BAcTrace should be sensitive enough to detect weak, functional connections. Consistent with this, experiments between neurons of high and medium strength connectivity showed consistent labelling patterns on the left and right sides of the brain and between animals (Fig. 3 and Fig. 4). In contrast, experiments on PN->LHN connections, which rely on fewer synapses, showed higher variability (Fig. 5). While this dichotomy could still result from technical issues, it likely reflects higher biological variability in weak connections. Addressing the origin and significance of variable connectivity throughout the nervous system will require analysis of circuit elements in large numbers of individuals; BAcTrace offers a scalable and affordable approach to characterising inter-individual connectivity.

In the last decade viral tracers such as rabies have been the primary source of connectivity data [31] but recently rapid advances in technology have made EM connectomics an increasingly viable approach [27]. Given that EM connectomics may reveal dense connectivity for all the neurons within a brain, one might wonder if it will supplant transsynaptic tracing going forwards. However, transsynaptic labelling has numerous advantages, which mean that these approaches should remain complementary rather than competitive for the foreseeable future. First, EM connectomics is limited to post-mortem specimens, while transsynaptic labelling methods can rapidly reveal connectivity in living animals, enabling neurons to be targeted for recording or manipulation or followed over time; furthermore connection based labelling can enable more precise labelling of neurons than can be achieved with genetic drivers alone. Second, EM connectomics will continue to remain prohibitively expensive and resource intensive for many laboratories and for studies requiring the comparison of multiple specimens. Third, although densely reconstructed connectomics datasets have the great advantage that they can reveal all the connections within a brain region, they are still missing important information e.g. identification of excitatory vs inhibitory synapses, electrical synapses, extra-synaptic communication; in contrast genetic tools can provide a readout for these properties. There are also cases in which the two approaches are synergistic: transsynaptic tools are particularly well suited to unequivocally link neurons identified in EM volumes to those labelled by genetic drivers since they simultaneously reveal neuronal morphology and connectivity; this process will be essential to the functional exploitation of the small number of reference connectomes that will become available over the next few years [50, 3].

Finally, the BAcTrace strategy should be applicable to other organisms such as mice and fish, which are still out of reach for whole brain EM imaging. Here the main hurdle for implementing BAcTrace might be the toxicity of the wild type toxin light chain. This can be tackled by using re-targeted light chains or even catalytically inactive light chains fused to non-toxic proteases which would ‘piggyback’ into the neuron [34]. We would encourage other scientists to improve (see supplemental text 7.3.1) and adapt BAcTrace to their specific needs. Many modifications could be implemented, tested and refined using the tissue culture tools and flies developed in this study.

## 4. Acknowledgments

This work was supported by a Dorothy Hodgkin Fellowship from the Royal Society to S.C. (DH120072), ERC Starting (211089) and Consolidator (649111) grants and core support from the MRC (MC-U105188491) to G.S.X.E.J. We are grateful to Bazbek Davletov, Enrico Ferrari and Jason Arsenault for sharing reagents and advice at early stages of the project. We acknowledge the Bloomington Stock Center and the Developmental Studies Hybridoma Bank for fly stocks and antibodies. Finally, we would like to thank Richard Benton and all members of the Jefferis group for many insightful comments on the manuscript.

## Supplemental data

### 6. Material and Methods

#### 6.1. Molecular cloning and transgenic flies

Backbones of plasmids used in S2 cell experiments were based on the *Drosophila* Gateway vector collection (kind gift from the Murphy lab [53]). Backbones of plasmids used for making transgenic flies were derived from pJFRC19 [41], pJFRC81 [42] and pJFRC161 [37]. Synthesised DNA sequences were codon optimised for *Drosophila* expression and made by GeneArt (Thermo Fisher Scientific, Inc) or IDT (Integrated DNA Technologies, Inc).

Plasmids were made using Gibson assembly [19]. All fragments were PCR amplified with overlapping primers yielding scarless products. All fusions were sequenced to control for mutations introduced during the cloning. GenBank accession numbers for all constructs can be found in Tables S10, S11, S12 and S13. Transgenic flies were made by BestGene Inc.

Briefly, components for each plasmid were generated as follows:

##### pAWG-hTfR::Syb

The intracellular portion and transmembrane segment of the hTfR followed by the extracellular segment of *Drosophila* Synaptobrevin were codon optimised and synthesised. The backbone of pAWG was PCR amplified. See Fig. 2A.

##### pAFW-hSNAP25

Full length hSNAP25 was PCR amplified from a plasmid source (kind gift from Bazbek Davletov). The backbone of pAFW was PCR amplified. The resulting fusion creates an N-terminal FLAG-tagged hSNAP25. See Fig. 2A.

##### pAFW-QF2::V5::hSNAP25::Syx

Amino acids 1-183 and 660-817 from QF were amplified by PCR and fused to create QF2 similarly to in [46]. Amino acids 141-206 from hSNAP25 were amplified adding a V5 tag to the 5’ end primer. Finally, full length *Drosophila* Syntaxin was amplified. All fragments were fused together with a PCR amplified pAFW backbone taking care that the N-terminal flag was in frame with the QF2. See Fig. 2A.

##### pGEX-kg-BoNT/A::GFPnb

BoNT/A protease and translocation domains were PCR amplified from a plasmid source (kind gift from Bazbek Davletov). A *Drosophila* codon optimised anti-GFP nanobody [28] was synthesised and PCR amplified. Both fragments were assembled together with the PCR amplified pGEX-kg backbone for expression in *E. coli*. In this construct there are two thrombin cleavage sites. One separates the GST from the toxin (used to release the toxin from the affinity column during purification) and the second thrombin site separates the toxin protease and translocation domains; this site is cleaved during purification and allows for light chain release after translocation.

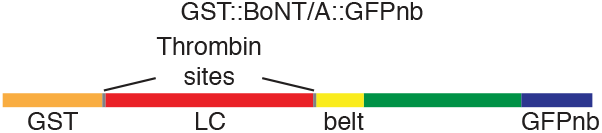

##### pJFRC81-BoNT/A::GFPnb::CD2

BoNT/A sequence was codon optimsed and synthesised. At the N-terminus a PAT3 signal peptide sequence from *C. elegans* was added for endoplasmic reticulum targeting. C-terminal to the toxin an anti-GFP nanobody was placed (codon optimsed, [28]) followed by a full length rat CD2 (excluding the first 20 amino acids encoding its signal peptide) ending with the ER export signal from Kir2.1. In between the toxin light and heavy chains and between the GFPnb and CD2 we introduced TEV and Thrombin cleavage sites, the latter for experiments not discussed in this work.

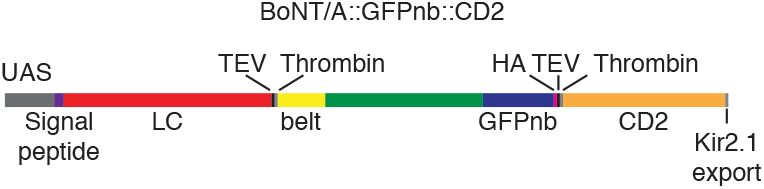

##### pAWH-BoNT/A-LC

BoNT/A-LC was codon optimised, synthesised and PCR amplified. Note that the codon optimisation was different from that used in pJFRC81-BoNT/A::GFPnb::CD2. The PCR fragment was then assembled with a pAWH backbone PCR amplified. The final product puts the LC in frame with a C-terminal HA tag from the vector.

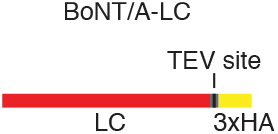

##### pAWH-TEV^T173V^::CD2

TEV^T173V^ was PCR amplified in two fragments from a plasmid source (non-condon optimised) with the T173V being introduced in the overlapping segment between both fragments. Full length rat CD2 was amplified by PCR. See Fig. S4C.

##### QUAS-3xHalo

3xHalo::CAAX-P10 was PCR amplified from a plasmid (Addgene #87646) and assembled with the backbone of QUAS-mtdTomato PCR amplified (excluding the mtdTomato). See [52].

##### UAS-B3RT-B2-B3RT-BoNT/A

B2 was PCR amplified from pJFRC153-20XUAS-IVS-B2::PEST (Addgene #32134). A tandem of Hsp70b and SV40 3’UTRs was PCR amplified followed by BoNT/A::GFPnb::CD2 amplified from pJFRC81-BoNT/A::GFPnb::CD2. B3 recombination sites were located in front of B2 and in front of the toxin signal peptide. In this way B3 recombination activity removes B2 and the Hsp70-SV40 UTRs placing the toxin under control of the UAS driven promoter.

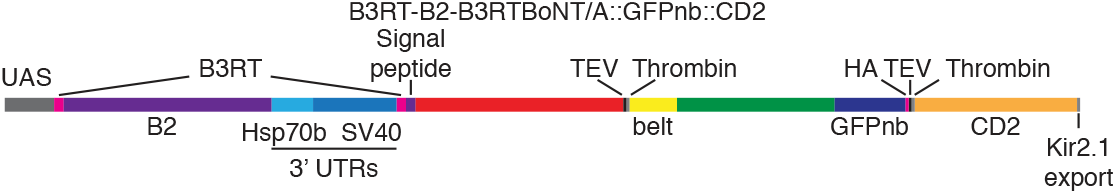

##### UAS-B3RT-BoNT/A

The B2-Hsp70-SV40 coding sequences were removed by crossing a fly containing UAS-B3RT-B2-B3RT-BoNT/A to a fly containing Nos-Gal4 (providing Gal4 in the germ line) and UAS-B3. Progeny from this cross was then PCR screened for lost of the B2 cassette. Isolated flies keep a B3RT site product of the recombination. We found this scar in the DNA to have no noticeable impact on Toxin expression (not shown).

##### UAS-B3RT-B2-B3RT-HBMBoNT/A

This construct was made by replacing the PAT3 signal peptide that drives the toxin into the ER by the HoneyBee Melittin signal peptide. HBM signal peptide was shown to be the most efficient peptide of several tested in the baculovirus protein expression system [51].

##### LexAop2-Syb::GFP-P10

Synaptobrevin was PCR amplified from genomic fly DNA. GFP-P10 was amplified from pJFRC81. The backbone of pJFRC19 was PCR amplified. See Fig. S7B.

##### LexAop2-QF2::V5::hSNAP25::Syx

QF2::V5::hSNAP25::Syx was PCR amplified from pAFW-QF2::V5::hSNAP25::Syx and assembled with the PCR amplified backbone of pJFRC19.

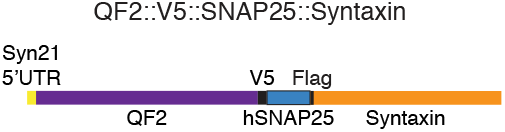

##### LexAop2-QF2::V5::hSNAP25::HIVNES::Syx

The flag tag present between hSNAP25 and Syntaxin in LexAop2-QF2::V5::hSNAP25::Syx was replaced by site directed mutagenesis with the nuclear export signal from HIV. See Fig. S6A.

##### LexAop2-QF2::hSNAP25::HIVNES::Syx

The V5 tag present between QF2 and hSNAP25 in LexAop2-QF2::V5::hSNAP25::HIVnes::Syx was replaced by site directed mutagenesis with a GlySer linker.

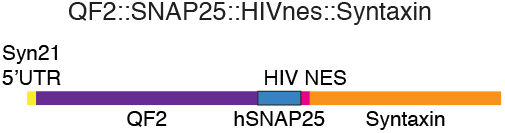

##### LexAop2-HIVNES::QF2::V5::hSNAP25::Syx

Two rounds of site directed mutagenesis were used on LexAop2-QF2::V5::hSNAP25::Syx to replace the flag tag between hSNAP25 and Syntaxin for a GlySer linker and to then introduce an HIV nuclear export signal in front of QF2. See Fig. S6A.

##### LexAop2-QF2::V5::hSNAP25::HIVNES::Syx-Hsp70UTR

Hsp70b 3’ UTR was PCR amplified from fly genomic DNA and assembled with the plasmid LexAop2-HIVNES::QF2::V5::hSNAP25::Syx amplified with a reverse primer at the end of Syntaxin and a forward primer after the SV40 3’UTR. This strategy replaces the SV40 3’UTR by the Hsp70 3’UTR. See Fig. S6A.

##### LexAop2-lowUTR-QF2::V5::hSNAP25::HIVNES::Syx-Hsp70UTR

The Syn21 5’ UTR from LexAop2-QF2::V5::hSNAP25::HIVNES::Syx-Hsp70UTR was replaced via site directed mutagenesis with the canonical *Drosophila* Kozak sequence CAAA. See Fig. S6A.

##### LexAop2-QF2-Syntaxin

Site directed mutagenesis was used to remove hSNAP25 from LexAop2-QF2::V5::hSNAP25::HIVNES::Syx. See Fig. S6A.

##### LexAop2-RSRT-Flp1-RSRT-TEV::CD2-P10

The Flp1 DNA recombinase was PCR amplified and a recombination site for R DNA recombinase (RSRT) added on the 5’ end. Hsp70-SV40 was amplified with a second RSRT site on the 3’ end. TEV was codon optimised and synthesised. Full length rat CD2 was PCR amplified. P10 was amplified from pJFRC81. The backbone of pJFRC19 was PCR amplified. Gibson assembly was used to generate LexAop2-RSRT-Flp1-RSRT-TEV::CD2-P10.

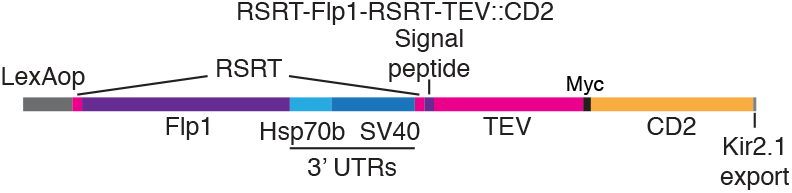

##### LexAop2-RSRT-TEV::CD2-P10

Flies with this insert were made by crossing LexAop2-RSRT-Flp1-RSRT-TEV::CD2-P10 flies to flies containing UAS-Flp and Nos-Gal4 (providing Gal4 in the Germ line). Progeny from this cross was then PCR screened for lost of the Flp1 cassette. Isolated flies keep an RSRT site product of the recombination. We found this scar in the DNA to have little or no impact on Toxin expression (not shown).

##### LexAop2-RSRT-Flp1-RS^T173V^::CD2-P10

The T173V mutation was introduced by site directed mutagenesis on LexAop2-RSRT-Flp1-RSRT-TEV::CD2-P10.

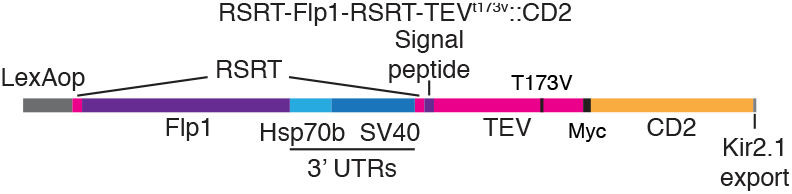

##### LexAop2-RSRT-T^T173V^::CD2-P10

Flies were made by removing the Flp1 cassette from LexAop2-RSRT-Flp1-RSRT-TEV^T173V^::CD2-P10 as in LexAop2-RSRT-TEV::CD2-P10.

##### LexAop2-RSRT-T^3xmut^::CD2-P10

A triple TEV mutant with all 3 potential glycosilation sites mutated was synthesised. Note that this sequence was codon optimised and synthesised using a different supplier than wild type TEV and therefore codon usage is slightly different. Following PCR amplification TEV was assembled together with the fragment resulting from PCR amplification of LexAop2-RSRT-TEV::CD2-P10 using a forward primer downstream of TEV and a reverse primer upstream of TEV. This strategy replaces TEV with TEV^3xmut^. Transgenic flies were made and the Flp1 cassette was removed as in LexAop2-RSRT-TEV::CD2-P10.

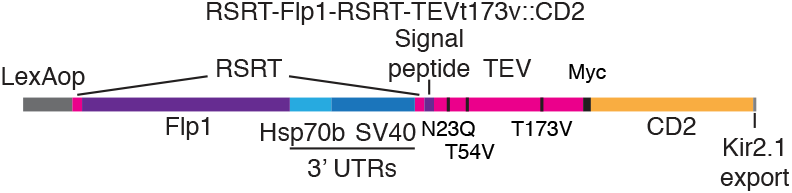

##### LexAop2-Syb::GFP^N146I^

The N146I mutation was introduced in LexAop2-Syb::GFP using site directed mutagenesis. See Fig. S7B.

##### LexAop2-Syb::GFP^N146I^::TEV

TEV was PCR amplified from LexAop2-RSRT-Flp1-RSRT-TEV::CD2-P10 and assembled with the PCR amplification of LexAop2-Syb::GFP^N146I^ using primers that anneal downstream of GFP^N146I^. See Fig. S7B.

##### LexAop2-Syb::GFP^N146I T173V^

TEV^T173V^ was amplified from LexAop2-RSRT-Flp1-RSRT-TEV^T173V^::CD2-P10 and assembled with the PCR amplification of LexAop2-Syb::GFP^N146I^ using primers that anneal downstream of GFP^N146I^. See Fig. S7B.

##### pAWG-mCherry::TEVs::V5::CD2

mCherry was PCR amplified from a plasmid source with a TEV cleavage site and a V5 tag on the reverse primer. Full length rat CD2 (excluding the first 20 amino acids encoding its signal peptide) was PCR amplified from a plasmid source incorporating the TEV cleavage site and V5 tag on the forward primer to allow for overlap with mCherry. The pAWG backbone was PCR amplified with overlapping primers in such a way that the mCherry::TEVs::V5::CD2 would be cloned in frame with the GFP from the backbone. See Fig. S7C.

##### pAWG-mCherry::V5::CD2

The TEV site in pAWG-mCherry::TEVs::V5::CD2 deleted using site directed mutagenesis. See Fig. S7C.

#### 6.2. S2 cell transfections

S2 cells were acquired from ThermoFisher Scientific (cat no R69007) and cultured according to the manufacturer recommendations. Once the culture was established the cells were transferred from Serum containing Schneider’s medium into increasing proportions of serum free Express 5 medium (ThermoFisher Scientific, cat no 10486025).

Plasmid DNA for S2 cell transfections was prepared using a MidiPrep DNA purification kit according to the manufacturer instructions (QIAGEN, cat no 12143).

S2 cell transfections were done in 6 well plates, each well containing 2ml of 1×10^6^ cells/ml. A total of 2µg of plasmid DNA was diluted with culture medium to a volume of 100µl and the mixture was vortexed. 3µl of FuGENE-HD transfection reagent (Promega, cat no E2311) were added to the diluted DNA, mixed gently and incubated for 10 minutes at room temperature. This mixture was then added drop-wise to the well with cells and the plate was put back into the incubator for 24h. Cells were then rinsed to remove transfection mix. Depending on the experiment either toxin was added as required (as shown in Fig. 2A) or cells were mixed 1:1 (as shown in Fig. 2B) followed by further 48h incubation before staining or western blot analysis. Plasmids for experiments shown in Fig. 2B are listed in Table S1, Fig. 2D in Table S2, Fig. S4A in Table S3, Fig. S4B in Table S4 and those from Fig. S4G in Table S5.

**Table S1:**
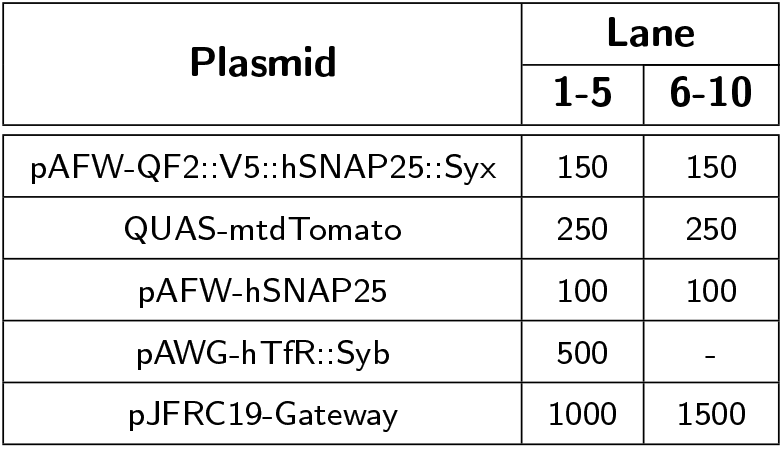
Amounts of plasmid DNA used for experiments presented in Fig. 2B. DNA amounts are in ng.

| Plasmid | Lane |  |
| --- | --- | --- |
|  | 1-5 | 6-10 |
| pAFW-QF2::V5::hSNAP25::Syx | 150 | 150 |
| QUAS-mtdTomato | 250 | 250 |
| pAFW-hSNAP25 | 100 | 100 |
| pAWG-hTfR::Syb | 500 | - |
| pJFRC19-Gateway | 1000 | 1500 |

**Table S2:** Amounts of plasmid DNA used for experiments presented in Fig. 2D. DNA amounts are in ng. R = Receiver and D = Donor.

| Plasmid | Lane |  |  |  |  |  |
| --- | --- | --- | --- | --- | --- | --- |
|  | 1,2,5,6R | 2D | 3,4,5D | 3,7R | 4,8R | 6,7,8D |
| pJFRC81-BoNT/A::GFPnb::CD2 | - | 1000 | 1000 | - | - | - |
| pAFW-hSNAP25 | 500 | - | - | 500 | 500 | - |
| pAWG-hTfR::Syb | 500 | - | - | 500 | - | - |
| pAWH-TEV<sup>T173V</sup>::CD2 | - | 500 | - | 500 | - | - |
| pBlueScript | 600 | - | 500 | 100 | 1100 | - |
| pMT-Gal4 | - | 500 | 500 | - | - | - |
| pAFW-QF2::V5::hSNAP25::Syx | 150 | - | - | 150 | 150 | - |
| QUAS-mtdTomato | 250 | - | - | 250 | 250 | - |
| pAWH-BoNT/A-LC | - | - | - | - | - | 2000 |

**Table S3:**
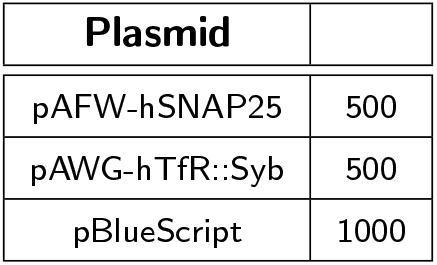
Amounts of plasmid DNA used for experiments presented in Fig. S4A. DNA amounts are in ng.

**Table S4:** Amounts of plasmid DNA used for experiments presented in Fig. S4B. DNA amounts are in ng.

| Plasmid | Lane |  |  |
| --- | --- | --- | --- |
|  | 1,4 | 2,5 | 3,6 |
| pAFW-QF2::V5::hSNAP25::Syx | 150 | 150 | 150 |
| QUAS-mtdTomato | 250 | 250 | 250 |
| pAFW-hSNAP25 | 100 | 100 | 100 |
| pAWG-hTfR::Syb | 250 | 500 | 1000 |
| pJFRC19-Gateway | 1250 | 1000 | 500 |

**Table S5:**
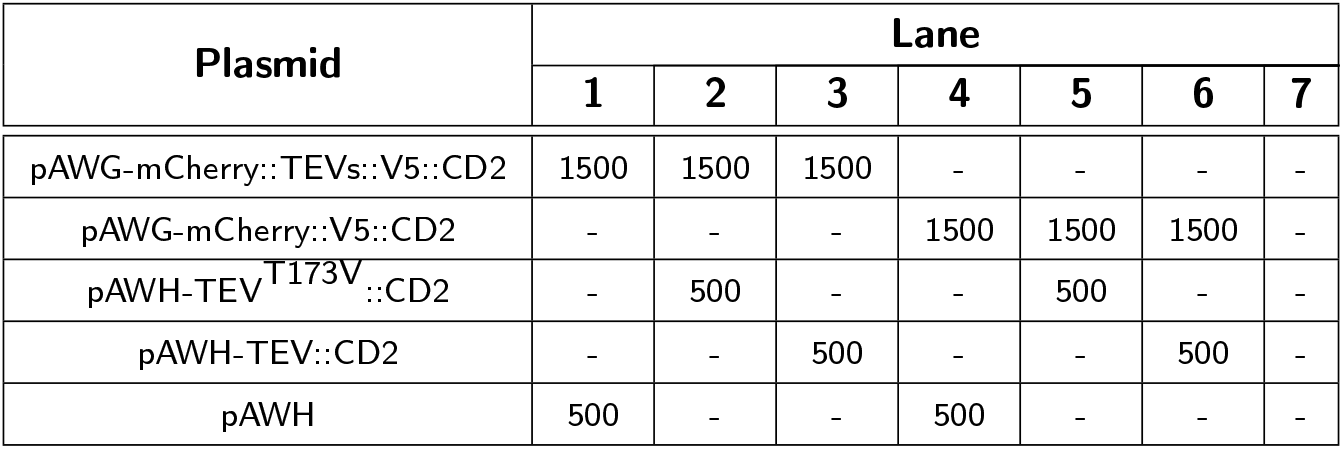
Amounts of plasmid DNA used for experiments presented in Fig. S4G. DNA amounts are in ng.

#### 6.3. BoNT/A::GFPnb purification from E. coli

Toxin was purified from *E. coli* using a low temperature expression protocol to improve toxin folding.

Day 1

1. Transform pLysS cells with pGEX-kg-BoNT/A::GFPnb or start a plate (TYE) from a glycerol stock.
2. Incubate at 37°C overnight.

Low temperature expression and purification of GST fusion proteins

Day 2

1. Pick a colony from a plate into 5ml 2xTY medium + 5*µ*l of both ampicillin 50mg/ml and chloramphenicol 50 mg/ml and incubate overnight at 37°C. Day 3
2. Inoculate 25ml of 2xTY medium + 25*µ*l of both ampicillin and chloramphenicol with 3ml of the overnight culture in a 50ml Falcon and incubate for 4h at 37°C.
3. Dilute into 1l of 2xTY medium + 1ml of both ampicillin and chloramphenicol in a 2l flask and grow for 1.5-2h at 37°C. Stop the incubation when the OD at 600nm reaches 0.6.
4. Add 1ml of IPTG 100mM to induce BoNT/A::GFPnb expression and **incubate overnight at 20°C.** Day 4
5. Pellet bacteria using a centrifuge at maximum speed for 20’ and discard the supernatant.
6. From here on work in ice. Suspend the pellet using 10ml of Lysis buffer. Transfer to a 50ml Falcon tube and rinse the bottle with an extra volume of 10ml. Prepare a 50X stock of Complete protease inhibitor cocktail, Roche (cat no. 4693132001) and add 600*µ*l.
7. Freeze the Falcon tube in liquid nitrogen for 10’ (can pause here by storing frozen pellet in −80°C freezer) and then thaw it in a water bath at room temperature.
8. Add 12*µ*l of 1M MgCl_2_ and a tip of deoxyribonuclease I from bovine pancreas. Rotate at room temperature for 10’.
9. Add Triton X-100 for a final concentration of 2% and incubate for 20’ at 4°C.
10. Equilibrate a GST column (Pierce, cat no 16107) to 4°C. Remove bottom tab by twisting and place in a 15ml Falcon tube.
11. Centrifuge column at 700g for 2min to remove storage buffer.
12. Equilibrate with two resin bed volumes (2-3ml) of equilibration/wash buffer from Pierce kit. Allow buffer to enter resin bed by gently inverting several times.
13. Centrifuge column at 700g for 2min to remove buffer.
14. Pellet the cell lysate using a centrifuge at 4°C for 30min at 7000g (we use reusable round bottomed PPCO 50ml tubes from Nalgene which fit a JA25.50 rotor). Save 20*µ*l of the supernatant for later SDS-PAGE analysis.
15. Add as much lysate to the column as fits (roughly 5-6ml) and allow it to enter the resin bed by gently inverting several times. Incubate at 4°C with rotation for 30min-1h.
16. Centrifuge the column at 700g for 2min, collect flow through and save an aliquot for SDS-PAGE analysis.
17. Repeat previous two steps until all the sample has been loaded on the column. About 3 times.
18. Wash resin with 2ml of equilibration/wash buffer from Pierce kit. Centrifuge at 700g for 2min and collect flow through for analysis. Repeat at least 2 times. Monitor the absorbance at 280nm and perform additional washes until the absorbance approaches baseline.
19. Keep some beads for SDS-PAGE analysis.
20. Wash resin with 2ml of buffer A twice to remove DTT and triton X-100.
21. Cap the bottom of the column with the white cap provided with the kit and then add 1.5ml of buffer A and 25 units of thrombin and incubate for 30min at 37°C in an orbital shaker at 100rpm.
22. Remove cap, put column in a 15ml falcon tube and spin at 700xg for 2min. Save the flow-through containing the toxin.
23. Repeat the thrombin treatment twice and save the flow-throughs.
24. Remove the GST-tags by rinsing the column with 2ml of elution buffer.
25. Centrifuge at 700xg for 2min.
26. Repeat the GST-tag elution twice more.
27. Regenerate the columns as recommended by the manufacturer.
28. Toxin can then be concentrated and buffer can be exchanged using Amicon spin columns. For BoNT/A::GFPnb toxin, buffer was exchanged for PBS, toxin was filtered and aliquoted and stored at −80°C.

**Table S6:** Solutions used during toxin purification.

| Lysis Buffer 500ml |  |
| --- | --- |
| 20mM HEPES | 10ml of 1M |
| 500mM NaCl | 50ml of 5M |
| 1mM EDTA | 1ml of 0.5M |
| 1mM DTT | 0.5ml of 1M |
| Buffer A 500ml |  |
| 20mM HEPES | 10ml of 1M |
| 100mM NaCl | 10ml of 5M |
| 2xTY 1000ml |  |
| Tryptone | 16g |
| Yeast extract | 10g |
| NaCl | 5g |
| Take volume to 1l, adjust pH to 7.4 if needed and autoclave |  |
| TYE plates |  |
| Agar | 15g |
| NaCl | 8g |
| Bacto Tryptone | 10g |
| Yeast extract | 5g |
| Take volume to 1l and autoclave. Pour on plates. |  |

#### 6.4. Western blot analysis

Cells for western blot analysis were resuspended in culture medium, pelleted, rinsed in PBS and pelleted again. Pellets were resuspended in 1x sample buffer (ThermoFisher Scientific, cat no NP0007) with a reducing agent (ThermoFisher Scientific, cat no NP0004) and heat denatured. Samples were then loaded in 12% bis-tris gels (ThermoFisher Scientific, cat no NP0342BOX) and run using MOPS buffer (ThermoFisher Scientific, cat no NP0001). Gels were transferred to PVDF membranes (Millipore Inc, cat no IPVH00010) and developed using the antibodies shown in Table S7 and ECL reagents (GE Healthcare Ltd, cat no RPN2232) following the manufacturer recommendations. HRP conjugated secondary antibodies were purchased from Cell Signalling Technology, Inc.

**Table S7:** Antibodies used in western blot experiments.

| Species | Target | Concentration | Supplier | Cat no |
| --- | --- | --- | --- | --- |
| Rabbit | FLAG | 1:1000 | Cell Signalling | 2368 |
| Mouse | rat CD2 | 1:2000 | GeneTex | GTX75123 |
| Chicken | GFP | 1:5000 | Abcam | ab13970 |
| Rat | HA | 1:4000 | Roche | 11 867 423 001 |

#### 6.5. Brain staining

For a full description of brain stainings see [39].

Briefly:

1. Dissect brains in 1xPB and keep them on ice.
2. Fix for 30’ in 4% paraformaldehyde in 1xPB.
3. Rinse 3-4 times with PBT, 10min each, in a rotating wheel.
4. Block between 1h and over night in block solution (5% normal goat serum in PBT).
5. Incubate in primary antibodies diluted in block solution for 2-3days at 4°C on a rotating wheel.
6. Wash 3-4 times with PBT, 2-3h each, in a rotating wheel at room temperature.
7. Incubate in secondary antibodies diluted in block solution for 2-3days at 4°C on a rotating wheel.
8. Wash 3-4 times with PBT, 2-3h each, in a rotating wheel at room temperature.
9. Equilibrate over night in Vectashield (Vector Laboratories, cat no H-1000).
10. Mount on slides and image.

In cases where chemical labelling and immunostaining were required, the former was done first as described in [52] and at the end of the protocol, instead of adding Vectashield, brains were put through the immunostaining procedure as described above. Solutions are listed in Table S8 and antibodies in Table S9.

**Table S8:**
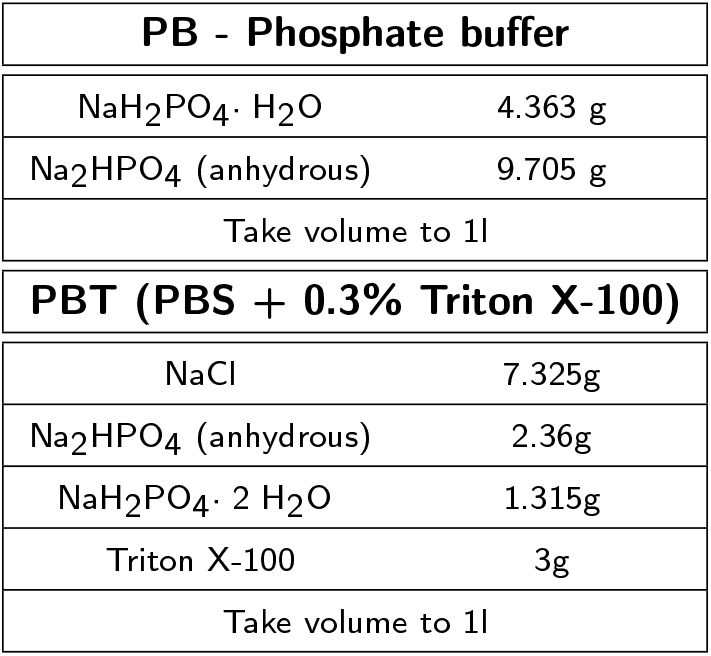
Solutions used for brain immunostainings.

| PB - Phosphate buffer |  |
| --- | --- |
| NaH<sub>2</sub>PO<sub>4</sub> · H<sub>2</sub>O | 4.363 g |
| Na<sub>2</sub>HPO<sub>4</sub> (anhydrous) | 9.705 g |
| Take volume to 1l |  |
| PBT (PBS + 0.3% Triton X-100) |  |
| NaCl | 7.325g |
| Na<sub>2</sub>HPO<sub>4</sub> (anhydrous) | 2.36g |
| NaH<sub>2</sub>PO<sub>4</sub> · 2 H<sub>2</sub>O | 1.315g |
| Triton X-100 | 3g |
| Take volume to 1l |  |

**Table S9:**
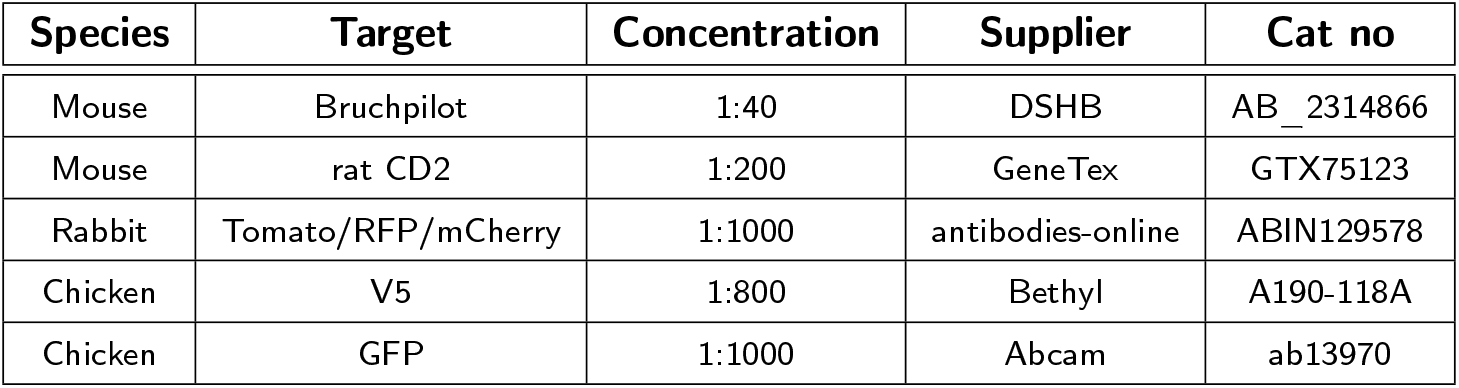
Antibodies used for brain immunostainings.

#### 6.6. Image acquisition

Confocal stacks of fly brains were imaged at 768 *×* 768 pixels every 1 µm (voxel size of 0.46 *×* 0.46 *×* 1 µm; 0.6 zoom factor) using an EC Plan-Neofluar 40*×*/1.30 Oil DIC M27 objective and 16-bit color depth. For LHN glomeruli scoring higher magnification images were taken at 1024 x 1024 pixels every 0.5 µm (voxel size of 0.19 x 0.19 x 0.5 µm; 1-1.1 zoom factor). All images were acquired using Zeiss LSM710 and Zeiss LSM880 confocal microscopes.

#### 6.7. Fluorescence quantification

To quantify the expression levels of Syb::GFP and QF2::V5::hSNAP25::Syx driven by the VT033006-LexA::P65 driver (S5) we co-stained brains for the neuropile marker Bruchpilot and GFP and Bruchpilot and V5 respectively. We then acquired high magnification confocal stacks of the stained ALs. Next we split the Zeiss LSM files into NRRD files using FIJI [49] followed by segmentation of each glomerulus by drawing a region of interest in the centre of the glomerulus (the plane which captured most of its area). Both glomerulus identification and segmentation were done using the nc82 channel. Segmentations were saved as ROI files. GFP and V5 mean intensities were obtained using FIJI and were normalised to the mean intensity of the nc82 channel for each glomerulus. This normalisation is intended to counter the drop in intensity due to light scatter when imaging deeper into the tissue.

#### 6.8. Drosophila stocks

Fly stocks were maintained at 25°C on Iberian food. The driver lines used in this study are summarised in Table S14, LexA responsive transgenes in Table S13, Gal4 ones in Table S12 and QUAS ones in Table S11. All brain images are from female flies.

#### 6.9. PN->LHN BAcTrace labelling quantification

We took high magnification confocal stacks of antennal lobes and used the nc82 channel to identify and annotate glomeruli using regions of interest in FIJI. We assigned a value of 0 (absent), 3 (present but weak) and 10 (strong) to annotated glomeruli by examining the intensity of the QUAS-Halo reporter channel. Once all antennal lobes were annotated we used the toxin labelling (CD2) channel to assign each specimen to the correct LHN cell type.

#### 6.10. Light microscopy PN-LHN overlap score

In order to quantify the overlap between neuronal skeletons for PNs and LHNs, derived from both light-level and EM data, we employed the ‘overlap score’ from [18]:

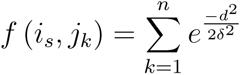

Skeletons were resampled so that we considered ‘points’ in the neuron at 1 µm intervals and an ‘overlap score’ calculated as the sum of *f* (*i_s_, j_k_*) over all points *s* of *i*. Here, *i* is the axonal portion of a neuron, *j* is the dendritic portion of a putative target, *δ* is the distance between two points at which a synapse might occur (e.g. 1 µm), and *d* is the euclidean distance between points *s* and *k*. The sum was taken of the scores between each point in *i* and each point in *j*.

Overlap scores were calculated between light-level reconstructions from stochastic labelling experiments [13, 8], that have been previously been registered from hundreds of brains to a common template, categorised and identified [9, 18]. We also made use of a complete set of uniglomerular PNs, reconstructed from a single EM dataset comprising a whole fly brain (Bates & Schlegel, in preparation).

#### 6.11. Electrophysiology

Electrophysiological recordings were carried out as described in [18] with minor modifications. Briefly, one day after eclosion flies were CO2 anesthetized, females of the correct genotype were selected, transferred to all trans-retinal fly food and kept in the dark. Five days later, flies were cold anaesthetised, placed in the recording chamber and dissected under dim light for recording as described in [26]. Data acquisition was performed as previously described with the only difference that a pco.edge 4.2 CMOS camera was used. For CsChrimson excitation of ORNs, a short (0.5 sec) pulse of light (550nm) was applied via a Cairn OptoLED controller.

### 7. Supplemental text

#### 7.1. BAcTrace is active in Drosophila cells

##### 7.1.1. Designing a BoNT/A::GFPnb receptor for S2 cells

While Syb::GFP was designed to target the toxin to neuronal neurotransmitter vesicles in flies, we reasoned it might not work in non-neuronal S2 cells. To overcome this limitation, in tissue culture experiments we replaced Syb::GFP with a chimera between GFP and the human transferrin receptor (hTfR::GFP). A similar construct has previously been shown to traffic to a low pH compartment in S2 cells [20]. We also tested chimeras between GFP and the *Drosophila* protein eater (FBgn0243514) and the scavenger receptor CI (FBgn0014033) and found that, similarly to GFP::hTfR, both were able to support Flag::hSNAP25 cleavage (not shown).

##### 7.1.2. Sensitivity of S2 cells to BoNT/A::GFPnb

All concentrations of toxin tested in Fig. 2B. showed nearly complete hSNAP25 cleavage. This motivated us to repeated the experiment using lower toxin concentrations. We found that as little as 0.1 to 0.3pM of toxin caused 50% hSNAP25 cleavage after 2 days of incubation (Fig. S4A). We also tried different amounts of plasmid encoding the toxin receptor and found that increases beyond 250 ug did not improve Flag::hSNAP25 cleavage, even though it resulted in higher amounts of GFP::hTfR as detected by WB (Fig. S4B).

##### 7.1.3. Optimising TEV for extracellular activity

To release the membrane tethered BoNT/A::GFPnb from the post-synapse we chose to use the Tobacco Etch Protease (TEV) fused to the transmembrane protein CD2. In plants, viral TEV is cytosolic while in our system TEV is targeted to the cell surface. We reasoned that post-translational modifications in the secretory pathway (e.g. glycosylation) might impact TEV’s activity. We tested this by co-expressing TEV::CD2 and mCherry::TEVsite::CD2::GFP (“TEV sensor” in Fig. S4C) in S2 cells. Fluorescence microscopy and western blot analysis indicated limited cleavage of the extracellular mCherry protein by wild type TEV (red solid arrowheads in Fig. S4D and G). The lack of extracellular TEV activity lead us to analyse TEV’s secondary structure where we found three predicted N-Glycosylation sites: N23, N52 and N171 (Fig. S4E and blue residues in F). We mutagenised the three sites individually (N23Q, T54V and T173V) and re-tested for Sensor cleavage activity. After mutating threonine 173 to valine (T173V) there was little mCherry signal left in the plasma membrane and the Sensor band was absent from the western blot (red open arrowheads in Fig. S4D and G). Consistently with this result, N171 has been shown to interact with the substrate during the catalysis while the other two sites are further away from the active site [43].

#### 7.2. BAcTrace works as a transsynaptic system in flies

##### 7.2.1. Using LexA to drive BAcTrace components in Receiver neurons

We reasoned that driving the detection system with LexA instead of tying it to pan-neuronal expression would allow more flexibility into the system; for example, Receiver neurons might have too many inputs and by using restricted LexA drivers these inputs could be immediately broken down by anatomical (e.g. using a LexA line specific for olfactory projection neurons), physiological (e.g LexA neurotransmitter specific) or other criteria for which LexA drivers are available.

##### 7.2.2. V5 tag is non-specifically cleaved inducing sensor background

To identify the source of and minimise the non-stochastic, toxin-independent background we made a series of flies carrying modified sensors (Fig. S6A). We hypothesised that one possible reason for the background could be the sensor not inserting properly in the membrane and therefore being free to migrate to the nucleus and activate the QUAS-effector in the absence of cleavage. To counter this we inserted the Nuclear Export Signal of the AIDS virus (HIV-NES) between hSNAP25 and Syntaxin. In this position this peptide will keep the protein out of the nucleus until toxin cleavage separates QF2 from it. Nevertheless we found this strategy failed to remove the background (Fig. S6A1 and B1). The HIV-NES is functional as inserting it N-terminal to QF2 removes any detectable background (Fig. S6A2 and B2).

We also designed two more sensors with weaker UTRs and found that expression level is not the main problem as both showed non-stochastic background. Nevertheless, we did confirm that the UTRs have an impact in expression levels as detected by the V5 tag (Fig. S6A3,4 and B3,4).

Lastly we made a construct lacking hSNAP25 to address the possibility of non-specific cleavage, maybe triggered by unusual interactions between hSNAP25 and *Drosophila* Syntaxin. To our surprise we found a much weaker V5 signal (compare Fig. S6B5 and B1) and a strong QUAS-Tomato signal, much stronger than the non-stochastic background.

Based on these set of results we reasoned that the source of the background is non-specific cleavage N-terminal to QF2 but not in the hSNAP25. The considerably weaker V5 signal in Fig. S6B5 suggests the V5 tag is being either cleaved and thus becomes non-immunogenic or cleavage in Syntaxin creates a more unstable cytosolic QF2::V5. Nevertheless, the fact that the HIV-NES in Fig. S6B1 is not active seems to support the cleavage of V5 as the source of background. We confirmed this hypothesis by creating a sensor like Fig. S6A1 but with the V5 replaced by a gly-ser linker. We found this sensor to be completely silent in the absence of toxin.

##### 7.2.3. TEV mutants modestly increase labelling efficiency

To further explore the impact of TEV in our system we repeated the experiments shown in Fig. 3F using the T173V mutant and a triple mutant in all three glycosylation sites. To allow the detection of increases in labelling efficiency for these experiments we used the split Gal4 line MB005C instead of MB247. MB005C drives toxin expression in a smaller subset of Kenyon cells (350 α’β’ neurons) than MB247-Gal4 (c. 1650 neurons of most types [1]). Using one or two copies of TEV::CD2 we could not detect an increase in labelling when compared to the no TEV control, implying that wild type TEV does not make toxin transfer more efficient (Fig. S7A1-3). Next we tested TEV^T173V^ (Fig. S4D-G) and a triple TEV mutant that abolishes all three sites (Fig. S4E). We found these mutants induce a modest increase in labelling efficiency of PNs. From this experiment it is not clear if the effect is due to cleavage between the toxin and CD2 (i.e. improving toxin release) or in the belt region (i.e. allowing LC detachment after translocation). The contribution of each site could be addressed by using toxins with only one TEV site.

Because extracellular TEV::CD2 is made inefficiently, as judged by antibody staining of CD2 (not shown), we reasoned that targeting it with Synaptobrevin instead of CD2 would increase its synaptic concentration and activity. Additionally, we wanted to mitigate the TEV independent toxin transfer mediated by Syb::GFP receptor. To this end we made a new receptor using a mutant GFP, GFP^N146I^, that has 10 time less affinity for the GFPnb than wild type GFP [28] (Fig. S7B1 and 2). We found this new receptor (Syb::GFP^N146I^) to be equally efficient as the one with wild type GFP (compare Fig. S7C1 and C2). We used this lower affinity receptor to target TEV. GFP staining indicated Syb::GFP^N146I^::TEV accumulates less than Syb::GFP^N146I^ possibly due to a de-stabilising effect of TEV (compare Fig. S7C2 and C3). Interestingly, this was partially alleviated by the T173V mutation in Syb::GFP^N146I^::TEV^T173V^(compare Fig. S7C3 and C4). Furthermore, while Syb::GFP^N146I^::TEV barely induced labelling of PNs the converse was true for Syb::GFP^N146I^::TEV^T173V^. This result mimics what we found in tissue culture, i.e. that the T173V mutant has increased activity, either due to increased stability or proteolytic activity.

##### 7.2.4. Subsets of Kenyon cells induce differential labelling of PNs

MB247-Gal4 is expressed in most KCs [1] and induces labelling on most Receiver PNs. In order to confirm this results and test BAcTrace at this synapse further, we used a panel of split Gal4 lines that label different subsets of KCs [2]. PN to KC connections in the adult fly are random [36] therefore we expected that the smaller the subset of KCs expressing BAcTrace the weaker the label would be, eventually resulting in fewer labelled PNs. In order to expedite the genetics in this experiment we made a new QUAS-3xHalo7 [52] reporter on the X chromosome. While this reporter recapitulates the previously used QUAS-mtdTomato it is considerably less sensitive (compare Fig. S9A and B).

A negative control missing a hemidriver failed to induce PN labelling (Fig. S8A). When both hemidrivers were present, expression in all subsets of KCs produced labelling in most Receiver PNs (Fig. S8B). Toxin expression in discrete KC subsets also induced labelling in Receiver PNs (Fig. S8C-E). Furthermore, qualitatively we didn’t see differences in the identity of the labelled PNs between the driver lines. However, the strength of labelling in PNs did not simply correlate with the number of KCs expressing toxin. For instance MB418B which only labels 140 α’/β’m neurons induces labelling as strongly as MB010B which labels 1940 neurons of most subtypes. Labelling strength was α’/β’ > α/β *≈* γ. This observation might be explained by a higher connectivity of α’/β’m KC than the other subtypes, more efficient toxin transfer through these synapses or stronger BAcTrace expression. The influence of expression strength can be seen in lines MB370B and MB463B which drive expression in the same cells at different levels and show different labelling efficiencies (Fig. S8C).

Another unexpected result was the relatively weak labelling induced by γd KCs. These neurons have dendrites in the ventral accessory calyx where they receive inputs from visual projection neurons and have been reported to not receive input from olfactory PNs [61, 59]. The labelling observed is unlikely to be of larval origin as most labelled PNs are not present in the larva. Several possible scenarios might contribute to the observed labelling: during metamorphosis there might be exploratory contacts from olfactory PNs which are later pruned, the use of non-localising toxin might have led to extra-synaptic contacts or alternatively there might exist real contacts between PNs and γd KCs which went undetected in the two aforementioned studies. In support of this later possibility γd KCs have process in the main calyx which are stained by the dendritic marker DenMark (see Fig. 2B in [59]). Higher magnification images of brains mounted dorsal side up also showed toxin puncta in the the calyx of the MB, in close apposition to labeled PNs (Fig. S8F). Using temporal control of toxin expression and localising the toxin to dendrites might shed more light on these results.

Lastly, we did experiments to assess the impact of components’ dosage in labelling efficiency. We used MB005B to drive toxin in 350 KCs. We found that having a second copy of the toxin or the receptor has little to no impact (Fig. S10A and B). On the other hand, having a second copy of the LexA driver induced a noticeable increase in labelling (Fig. S10C). While background staining increased as well, this experiments were done with the V5 containing sensor; we anticipate background would be mostly absent when using the newer sensor. A note of caution is due here, in this and other experiments using two copies of VT033006-LexA::P65 we noted unusual PN morphologies, e.g. axons terminating early and with simpler arborisations. This anomalies could be due to high expression levels of BAcTrace components, a toxic effect of LexA::P65 or the homozygous state of the VT033006-LexA::P65 insertion locus (JK22).

#### 7.3. Discussion

##### 7.3.1. System Optimisation

Based on our results there are several areas where optimisation might increase the sensitivity and flexibility of the system:

**Receptor-Ligand:** Fine-tuning this interaction could increase the efficiency of the transfer. For instance multimerising the GFP on the Syb::GFP would make more ligand available while multimerising the GFPnb in the toxin would allow each toxin to bind more than one receptor; if the limiting step is the force the receptor exerts to pull the toxin across the cleft then binding 2 or 3 receptors instead of 1 might make this process more efficient. In addition GFP could be mutated to make it non-fluorescent or the receptor ligand pair could be replaced altogether.

**Sensor**: Our experiments used a full length *Drosophila* Syntaxin as carrier for the QF2::hSNAP25. Syntaxin and SNAP25 are part of the SNARE complex in neurons. In the sensor, these helices are in close proximity potentially pushing them into forming a complex, with or without Synaptobrevin, the third component of the SNARE complex, which could affect the availability of hSNAP25 for toxin cleavage. Once the cleavage takes place, the interaction between the helices could also affect the release of the QF2 which retains 48 amino acids from hSNAP25 on its C terminus (see Fig. 1.A). To explore and potentially mitigate the negative impact of full length Syntaxin, deletions of the sensor should be tested. In addition, we found the Syntaxin sensor to be lethal when expressed very broadly and at high level within the nervous system (e.g. when driven with Syb-LexA::P65). We attribute this toxicity to the overexpression of the Syntaxin portion of the sensor as a similar sensor based in Synaptobrevin did not show lethality, although it was considerably less sensitive (not shown).

**Genetics:** Transsynaptic systems are by design extremely sensitive and will produce false positives when used with leaky Gal4 lines or when Gal4 protein persist in the adult after developmental expression. For this reason all genetically encoded tracing tools will benefit from better genetic means by which to express tracers. In this study we took a recombinase based approach to activate the toxin transgene and decrease background. While this is sufficient for the very clean Gal4s in combination with the very restricted LexA line used here, it will not be enough for many other applications. Approaches that conditionally refine expression in time by means of temperature sensitive components (e.g. the temperature sensitive Gal4 repressor Gal80) or fine-tuning the stability of the components will help tackle this issue.

One important limitation of BAcTrace is that the LexA and Gal4 lines cannot overlap. If they do, very strong activation takes place in the overlapping cells whereby the toxin gets internalised becoming unavailable to induce transsynaptic labelling. In future systems the same recombinase that activates the toxin transgene could be used to remove the DNA encoding the receptor allowing for overlapping Gal4-LexA expression. This might need to be complemented by a mechanism to trigger the degradation of receptor made before the DNA recombination.

In addition, new implementations of the system might shy away from using Gal4 and LexA as direct drivers but they could activate recombinases that in turn switch on component transgenes. In this way these transgenes could be driven by pan-neuronal promoters making expression levels less variable and resulting in more comparable results between cell types.

#### 7.4. Supplemental figures

**Figure S1:**
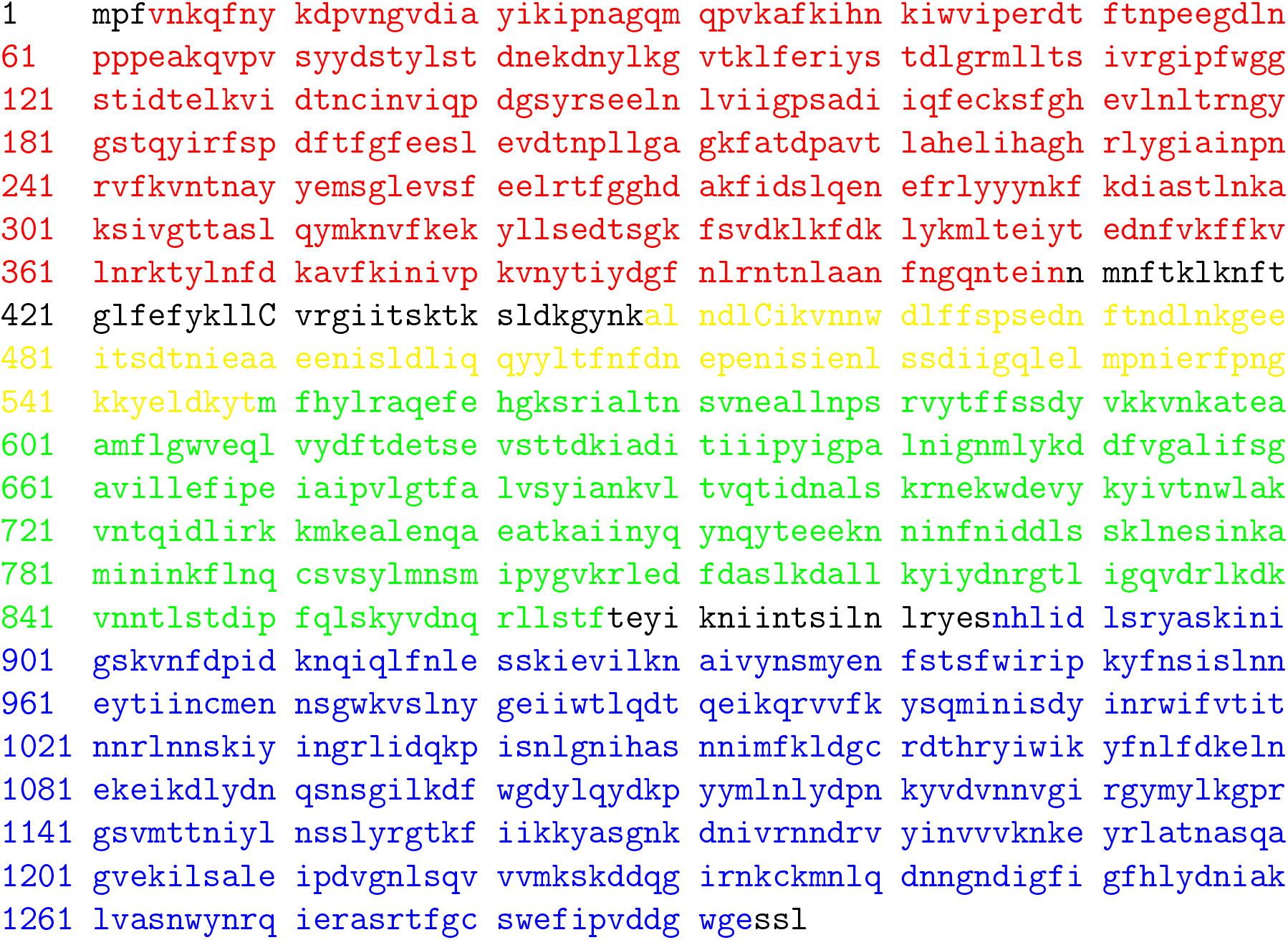
Amino acid sequence of BoNT/A1 from Clostridium botulinum (AFN57627.1). The protease domain is indicated in red, belt in yellow, translocation domain in green and receptor binding domain in blue, same as in Fig. 1A. Cys 430 and 454 which form a disulphide bond to hold the light and heavy chains together are indicated in capital.

**Figure S2:**
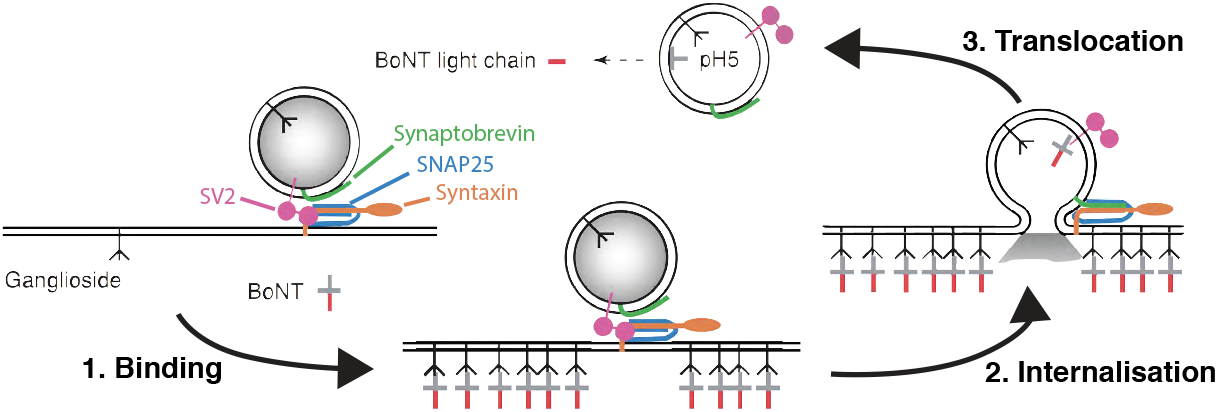
BoNT/A1 mechanism of action in vertebrates. Adapted from [11].

**Figure S3:**
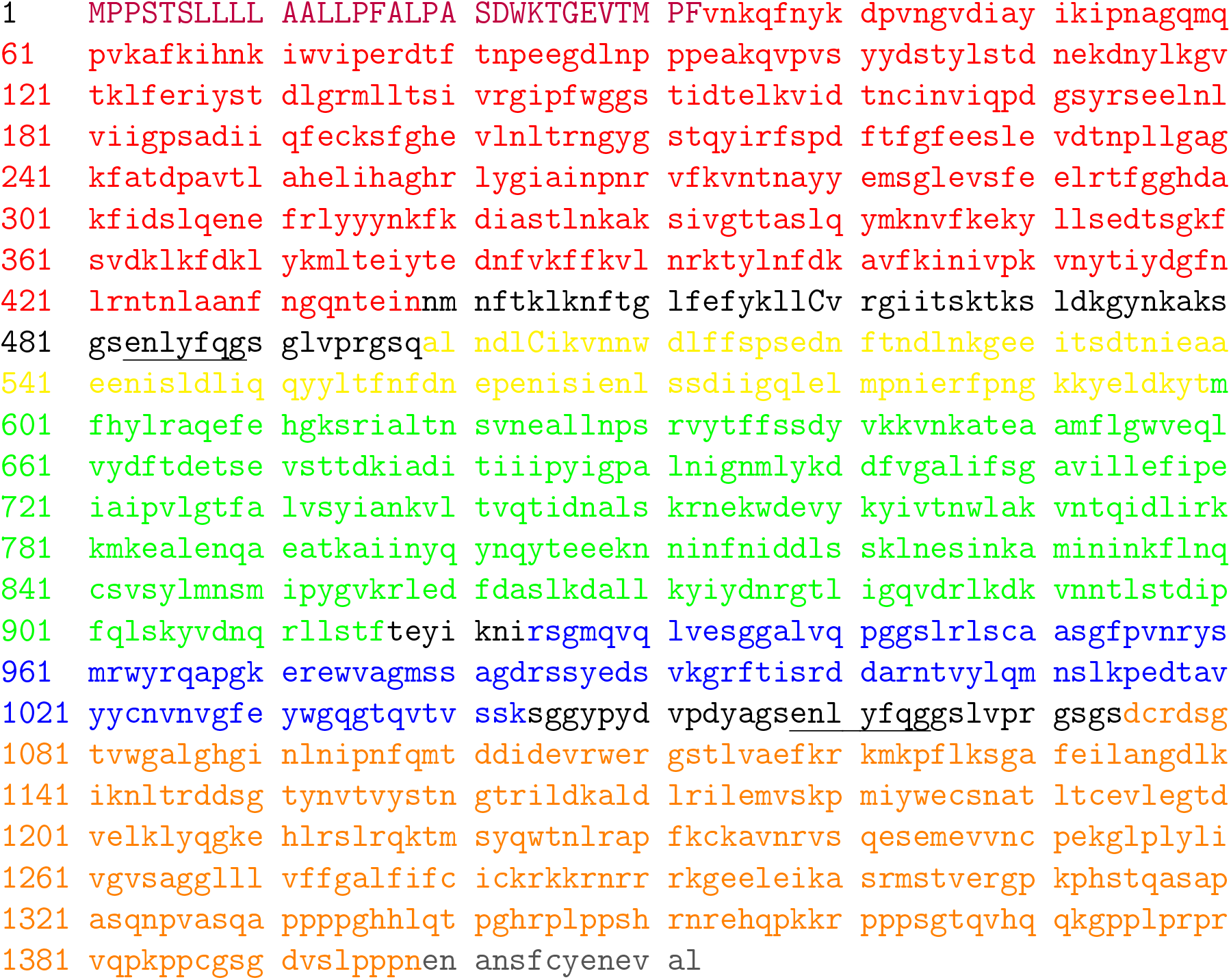
Amino acid sequence of BAcTrace BoNT/A::GFPnb::CD2. Colours as in Fig. S1 except the signal peptide for ER targeting which is indicated in purple and capital, the anti-GFP nanobody in blue, rat CD2 in orange and the C-terminal Kir2.1 ER export signal to improve trafficking to the plasma membrane in grey [32]. TEV sites for toxin release are underlined.

**Figure S4:**
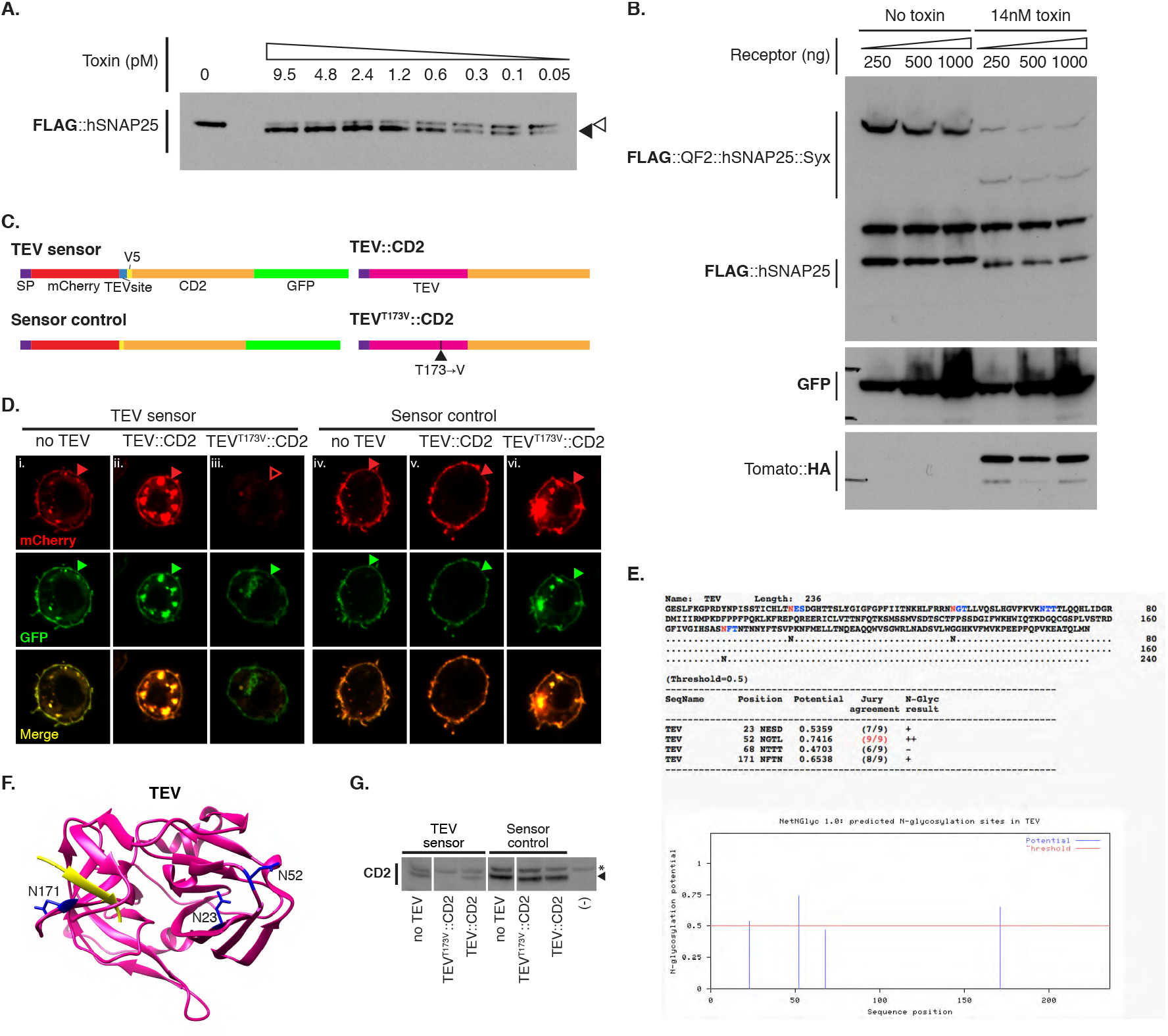
Testing BAcTrace in tissue culture. (**A**) Western blot analysis of S2 cell extracts after transfection and incubation with decreasing amounts of toxin. At around 0.3pM half of the hSNAP25 appears to be cleaved. The empty arrowhead indicates un-cleaved products while the solid arrowhead indicates cleaved products. (**B**) Flag::hSNAP25 cleavage efficiency does not change at the tested concentrations of hTfR::GFP receptor. (**C**) Schematics of TEV constructs used in (D) and (G). (**D**) Immunochemistry of S2 cells transfected with the indicated sensors and TEV variants. Solid arrowheads indicate non-cleaved sensor on the plasma membrane and empty arrowheads indicate cleaved sensor on the plasma membrane as determined by the presence of red and absence of green fluorescence. (**E**) Prediction of N-glycosilation sites on TEV [21]. (**F**) TEV structure (1lvb, [43]) with potential glycosilation sites from F highlighted in blue. Bound pseudosubstrate is shown in yellow. (**G**) Western blot against CD2 showing that only the TEV^T173V^ is capable of cleaving the TEV sensor. The band marked with an asterix is non-specific. Band marked with a solid black arrowhead corresponds to the TEV sensor. Epitopes detected by antibodies are indicated in bold in (A), (B), (D) and (G).

**Figure S5:**
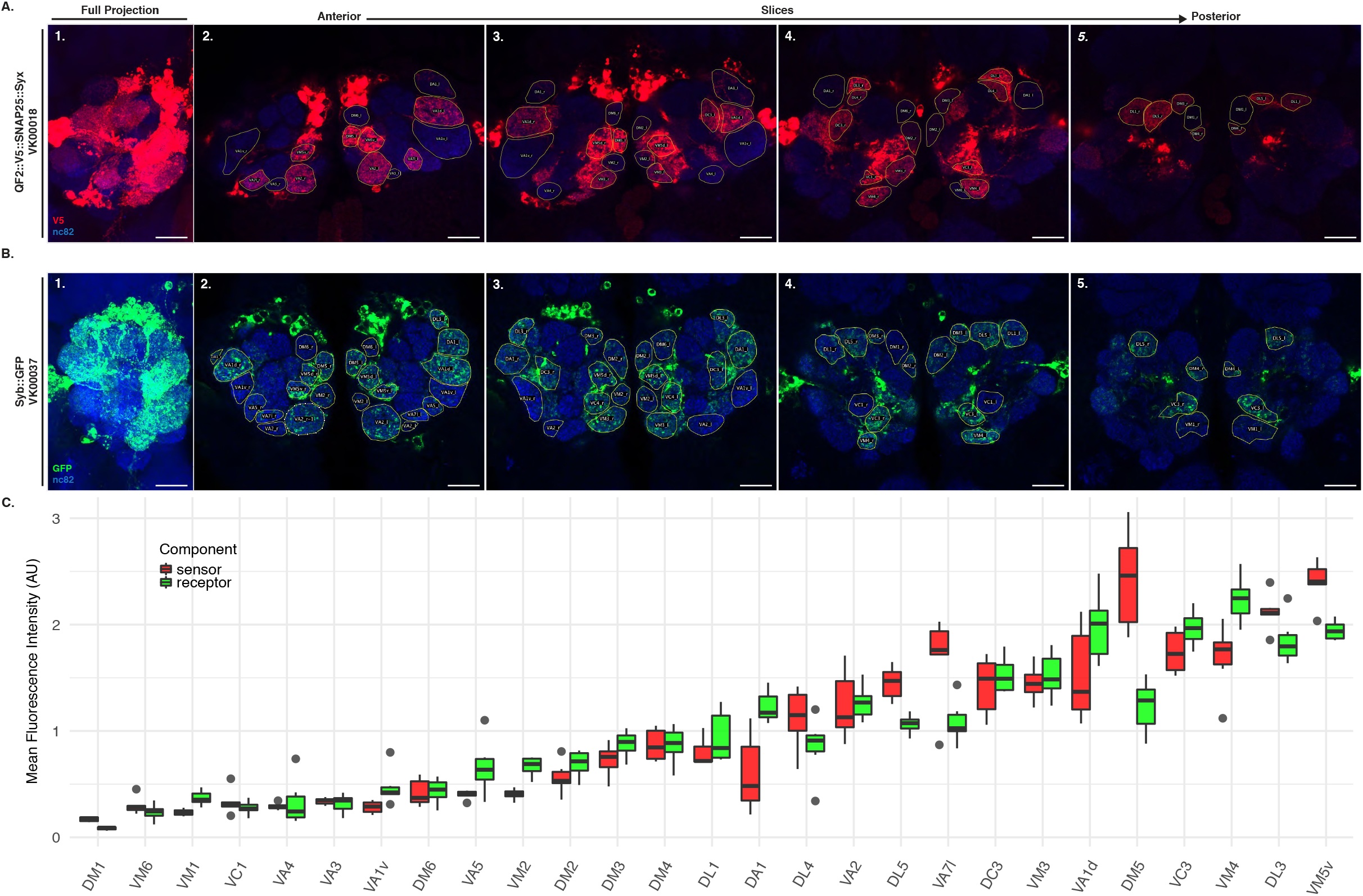
Expression strength of VT033006-LexA::P65. (**A**) V5 tag immunostaining of QF2::V5::hSNAP25::Syx in VK000018. (**B**) GFP immunostaining of Syb::GFP in VK000037. (**C**) Quantification of fluorescence per glomerulus from A and B. Each glomerulus measured in 6 ALs from 3 brains. Leftmost panel in (A) and (B) are maximum intensity projections while all other panels are single slices from confocal microscopy stacks. Epitopes detected by antibodies are indicated in bold in A and B. Scale bars 30µm.

**Figure S6:**
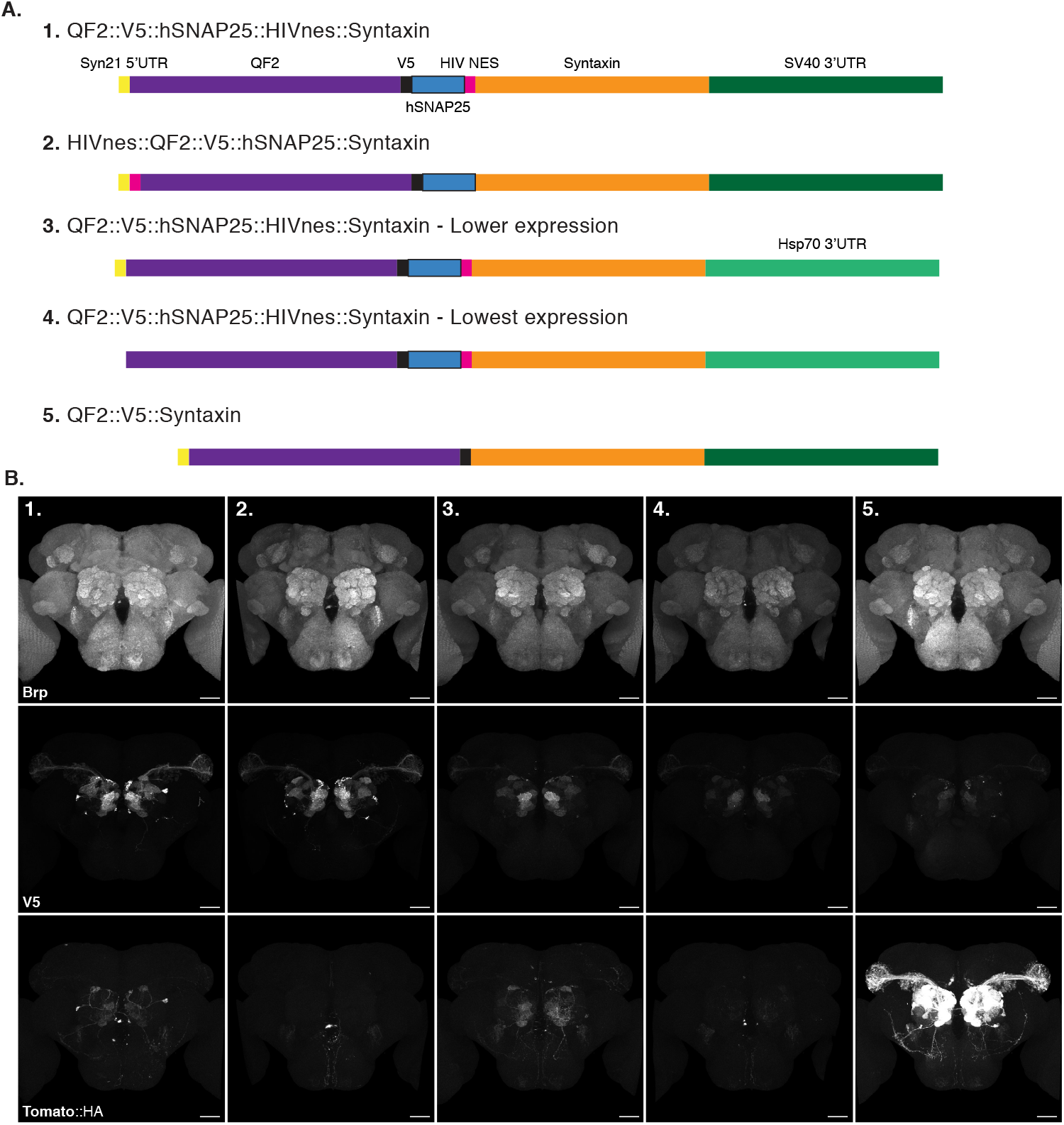
V5 tag induces non-specific activation of the toxin sensor. (**A**) Schematic of the sensors used in (B). (**B**) Background expression of the sensors from (A) in the absence of toxin. Maximum intensity projections of registered confocal stacks from age matched animals; images taken using the same microscope settings. Epitopes detected by antibodies are indicated in bold in B. Scale bars = 30µm.

**Figure S7:**
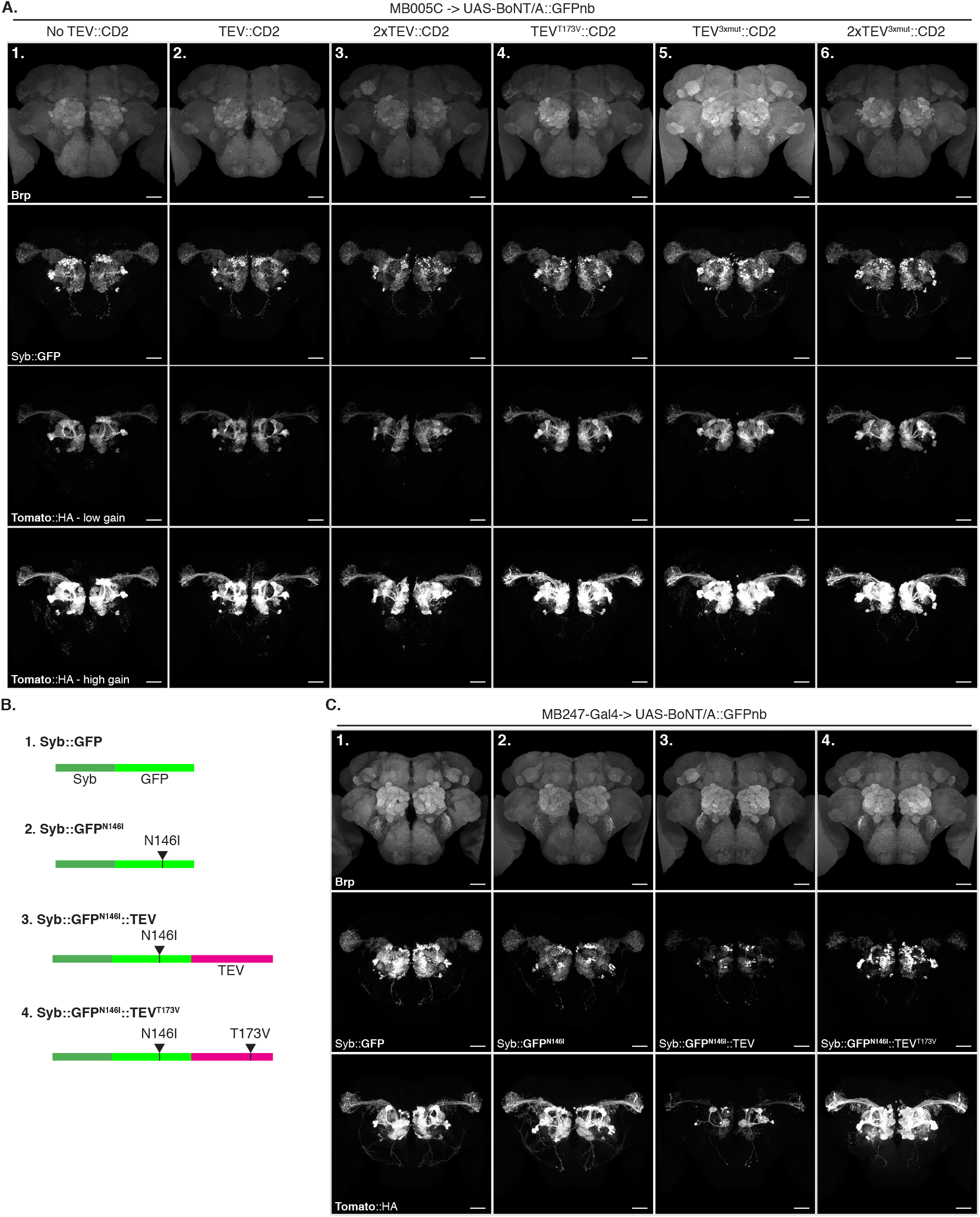
TEV^T173V^ modestly increases BAcTrace efficiency. (**A**) BAcTrace labelling of PNs from sparse KC Donors driven by MB005C. (**A1**) TEV negative control, (**A2**) one copy and (**A3**) two copies of wild type TEV show no difference in labelling efficiency. All (**A4**), one copy of TEV^T173V^, (**A5**), one copy of TEV^3xmut^ and (**A6**), two copies of TEV^3xmut^, show a modest increase in labelling when compared to the no TEV control (A1). (**B**) Schematic of constructs used in (C) for minimising TEV independent labelling. (**C**) (**C1**) Control with the regular Syb::GFP receptor. (**C2**) Lower affinity Syb::GFP^N146I^ receptor still supports efficient transsynaptic labelling. (**C3**) Addition of wild type TEV to the Syb::GFP^N146I^ in (C2) decreases expression and transssynaptic labelling. (**C4**) Introducing T173V mutation in TEV from (C3) increases expression strength and labelling efficiency. Maximum intensity projections of registered confocal stacks from age matched animals; images taken using the same microscope settings. Epitopes detected by antibodies are indicated in bold in (A) and (C). Scale bars 30µm.

**Figure S8:**
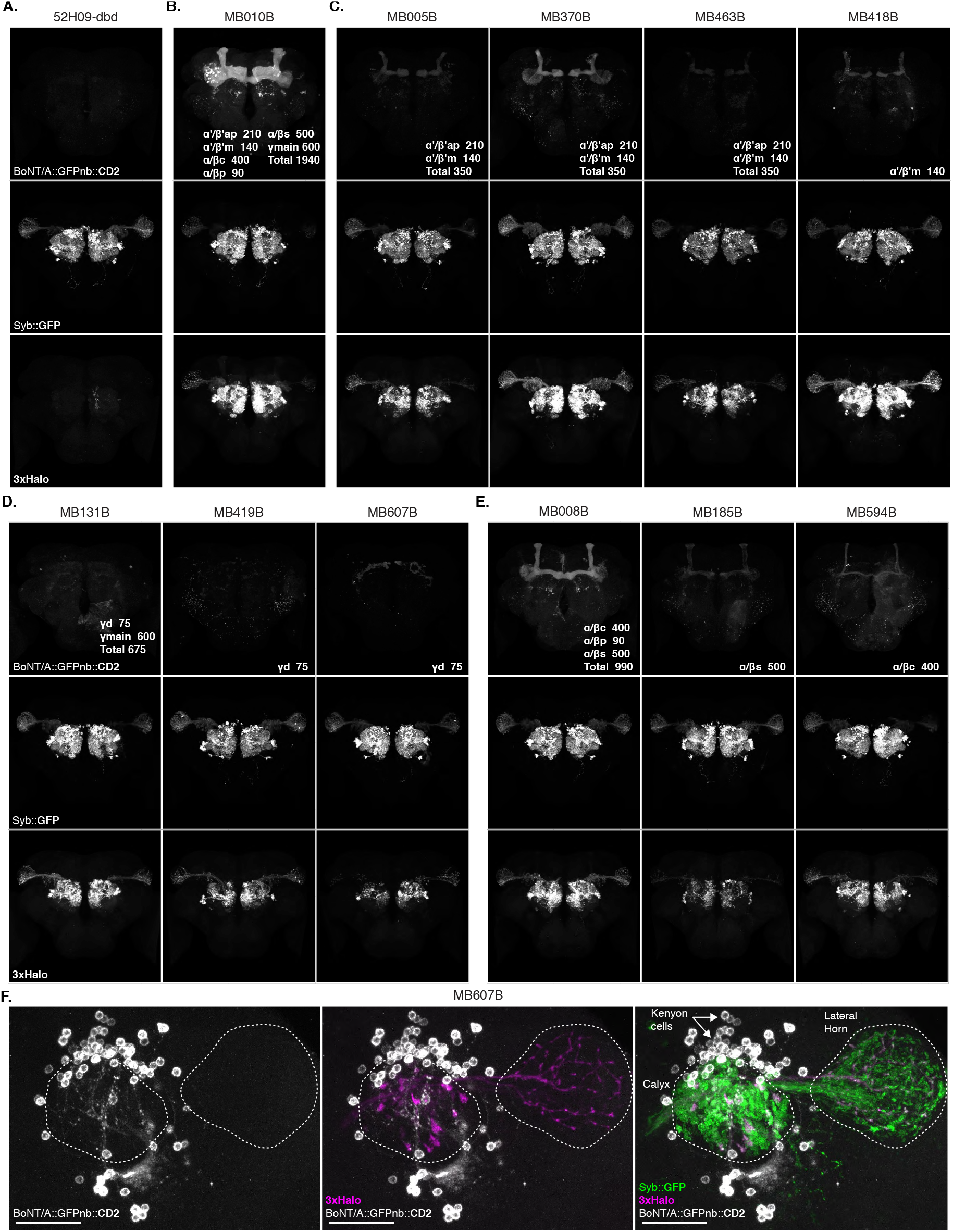
BAcTrace expression in subsets of KCs induces labelling in PNs. (**A**) Negative control missing a split Gal4 hemidriver. Donor KCs: (**B**) all, (**C**) α’/β’, (**D**) γ and (**E**) α/β. (**F**) Higher magnification view of the MB calyx and LH of a brain with Donor MB607B neurons (same as in D). The brain was mounted dorsal side up to provide better image quality of the MB calyx and LH. Within each lobe subtypes are: γ lobe: main (m) and dorsal (d), α’/β’ lobe: anterior-posterior (ap) and middle (m) and α/β: posterior (p), core (c) and surface (s). Maximum intensity projections of registered confocal stacks from age matched animals; (A), (B), (C) and (D) taken using the same microscope settings. Epitopes detected by antibodies are indicated in bold in each panel. Number of KCs per subtype from [2]. Scale bars 30µm.

**Figure S9:**
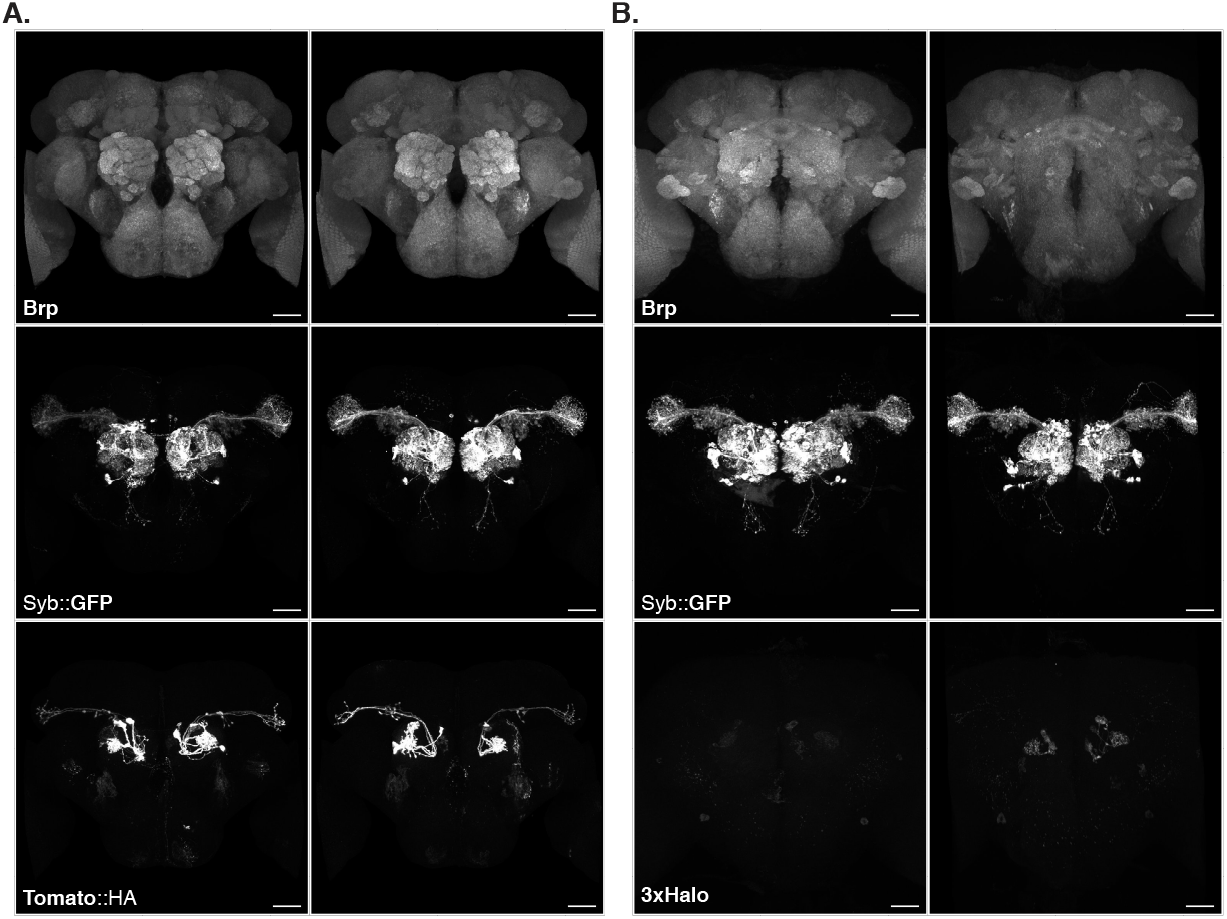
Sensitivity of BAcTrace QUAS-reporters. BAcTrace labelling of VA1d PNs using Or88a ORNs as Donors. (**A**) Labelling using QUAS-mtdTomato #26(94E7) from [45]. (**B**) Labelling using QUAS-3xHalo (this work). Maximum intensity projections of registered confocal stacks from age matched animals; images taken using the same microscope settings. Epitopes detected by antibodies are indicated in bold in each panel. Scale bars 30µm.

**Figure S10:**
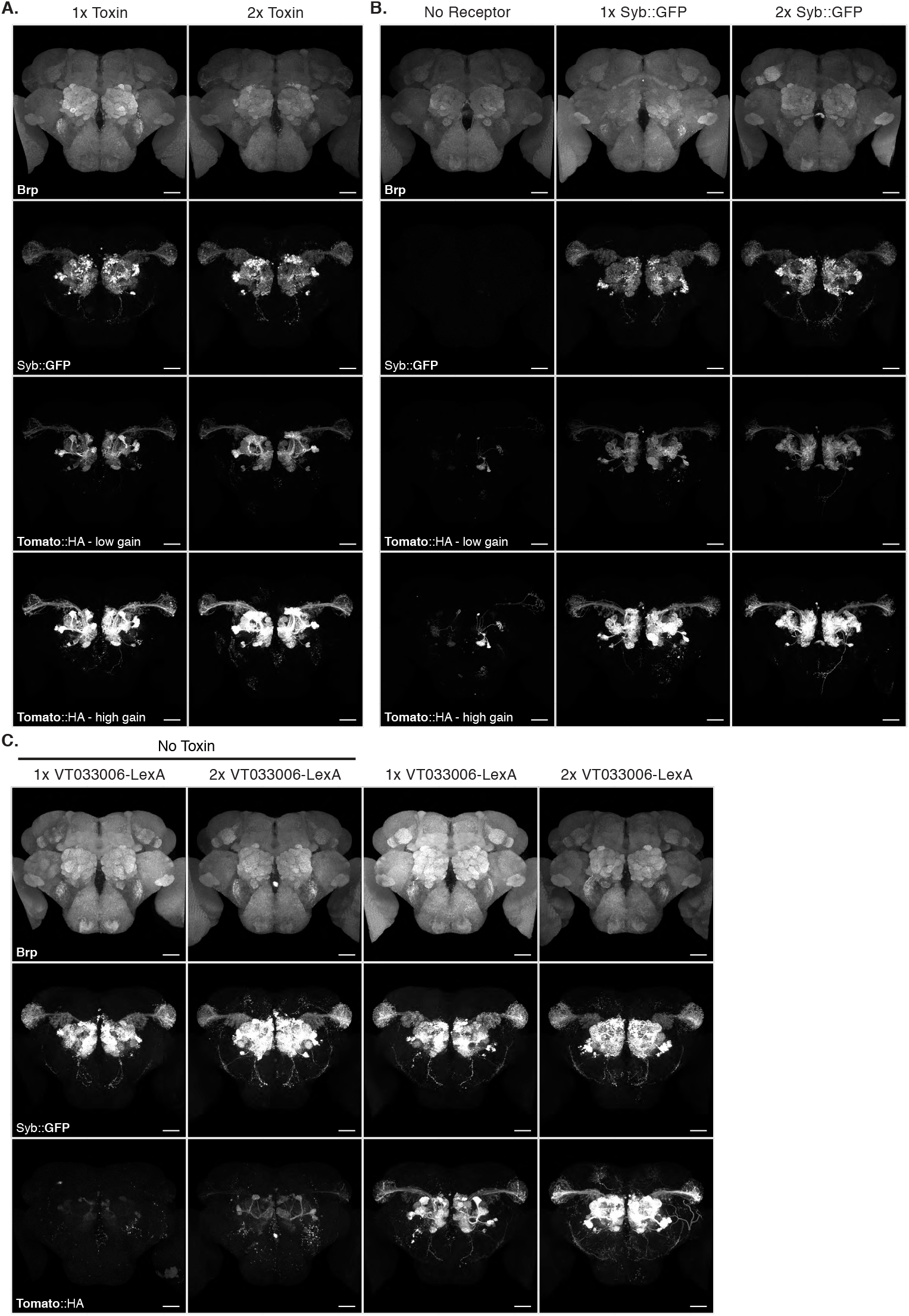
Effect of manipulating BAcTrace components dosage. In all experiments we used MB005B (Fig. S8) to drive toxin in rv350 Donor α’/β’ KCs. (**A**) Compared to a single copy, two copies of Toxin induce a modest increase in labelling. (**B**) A second copy of receptor induces no increase in labelling. (**C**) A second copy of the LexA driver induces an increase in labelling strength. Note that the sensor used in these experiments has a V5 tag which is the likely source of most background in the absence of toxin. Maximum intensity projections of registered confocal stacks from age matched animals; images taken using the same microscope settings. Epitopes detected by antibodies are indicated in bold in each panel. Scale bars 30µm.

**Figure S11:**
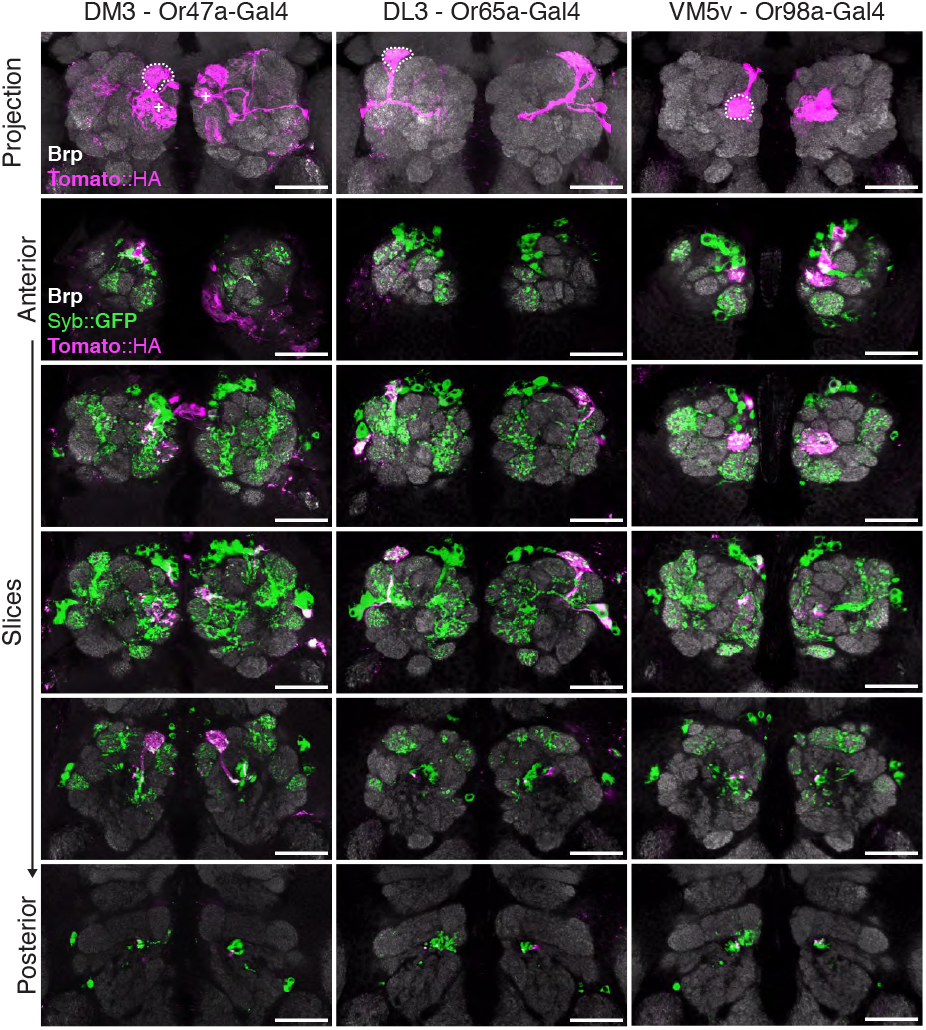
BAcTrace expression in ORNs labels connected PNs. BAcTrace expression in specific ORNs induces labelling in connected PNs (dotted lines). Occasionally labelling is also induced in PNs from neighbouring glomeruli and less frequently non-neighbouring ones. The age of shown animals were: DM3 10-13 days old, DL3 and VM5v 2-4 days old. Top panel is a maximum intensity projections of registered confocal stacks. 5 next panels are single slices from the same specimen. Epitopes detected by antibodies are indicated in bold in each panel. Scale bars 30µm.

**Figure S12:**
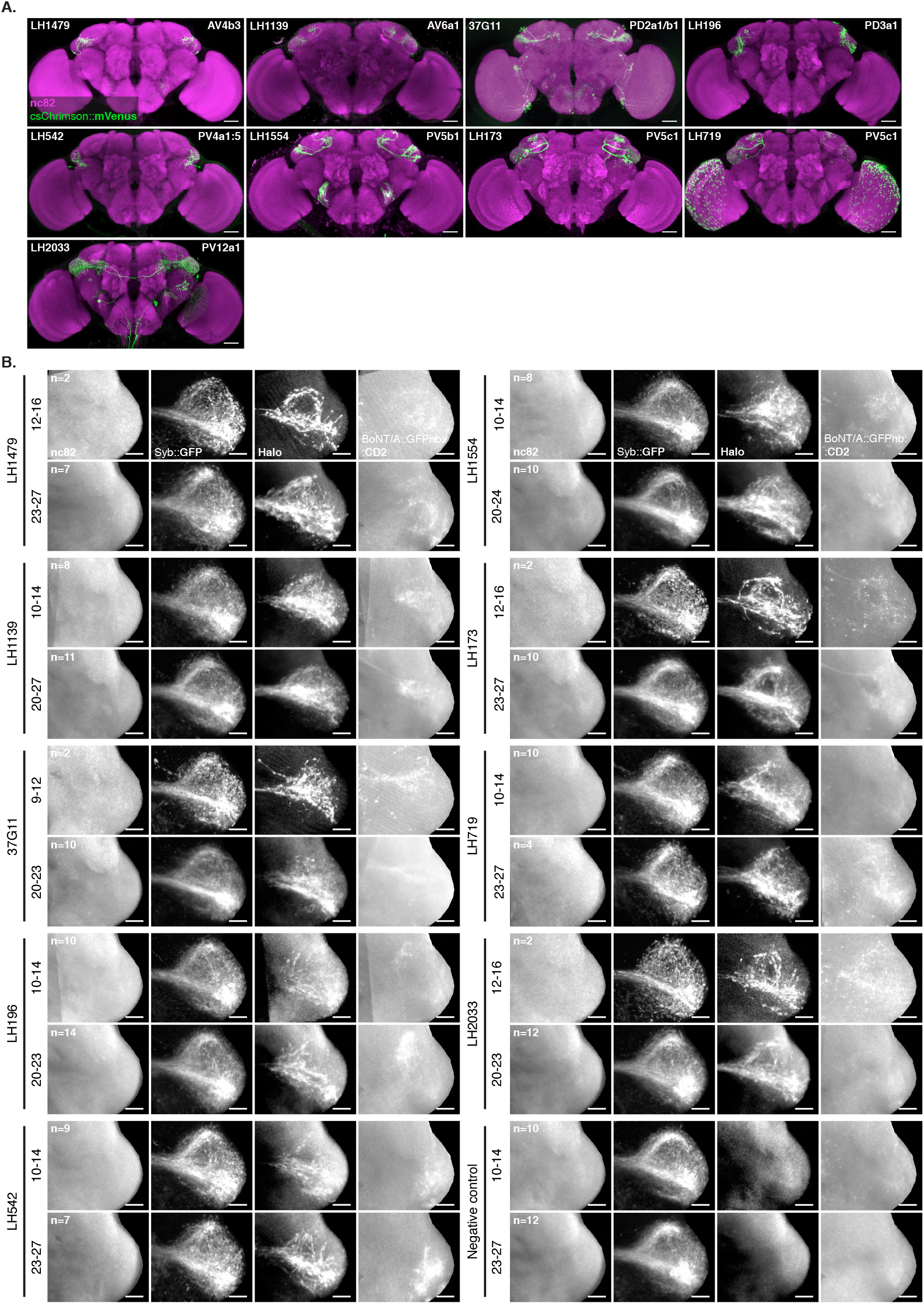
BAcTrace expression in LHNs induce labelling in PNs. (**A**) Expression of split Gal4 LHN lines tested. Anti-GFP immunostaining against UAS-csChrimson::mVenus in attP18. Cell types in each line are indicated on the top left and the name of the line on the bottom right. Adapted from [13] (**B**) For each line animals of two ages were dissected. Several LHs were imaged, registered to a template and averaged to produce the different panels. Bottom two panels on the right are a negative control made from line LH196 with one hemidriver missing. Epitopes detected by antibodies are indicated in bold in each panel. Scale bars (A) = 30µm and (B) = 10µm.

**Figure S13:**
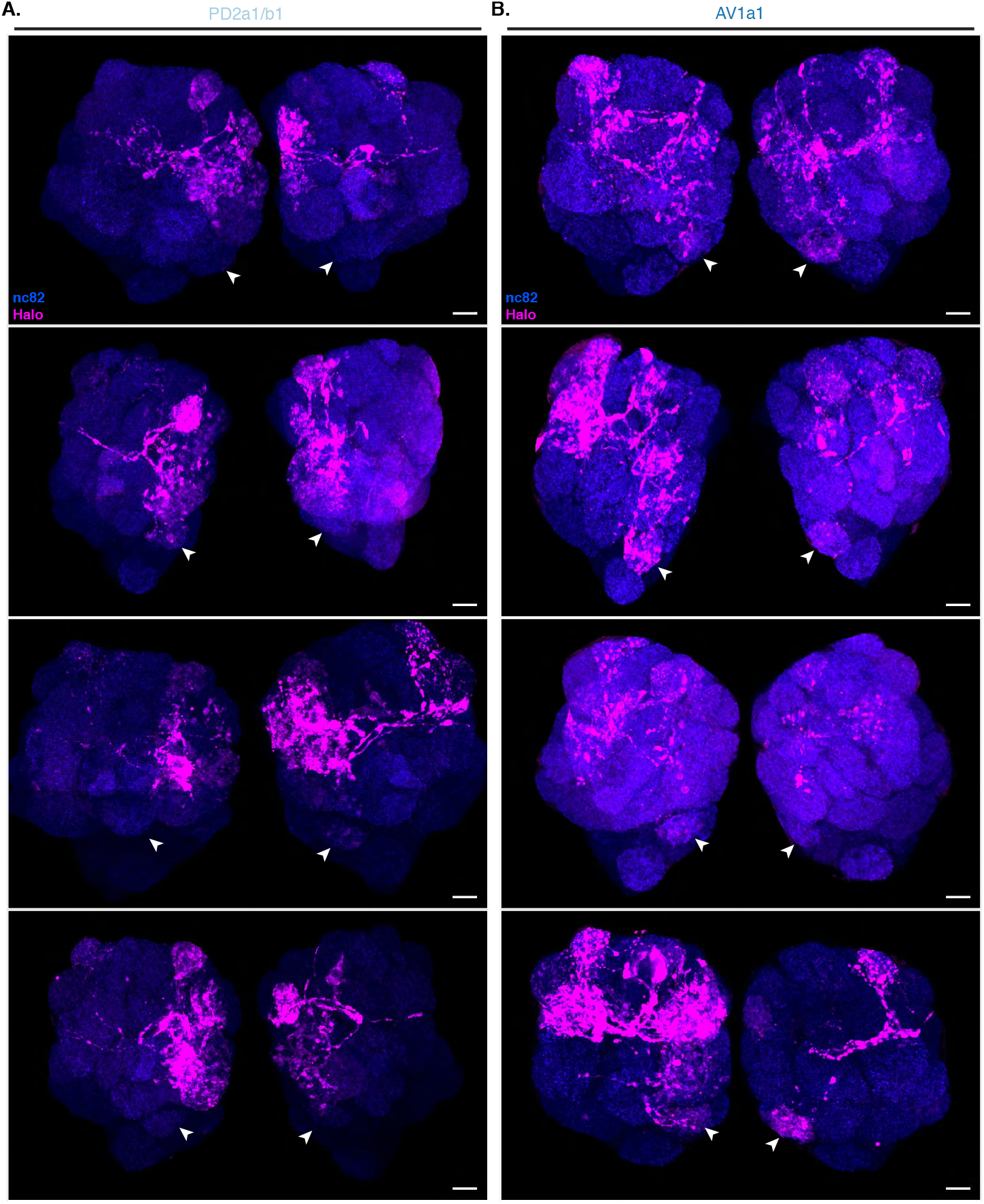
LHN-PN connectivity for 2 LHN types in 4 different specimens. BAcTrace labelled PNs from Donor cells: (**A**) PD2a1/b1 neurons (LH989) and (**B**) AV1a1 neurons (LH1983). Images are maximum intensity projections of confocal stacks, affine registered to a template for the AL region. The surface outside the neuropile region was masked out to avoid obscuring the glomeruli. Epitopes detected by antibodies are indicated in bold in each panel. Scale bars = 10µm.

**Figure S14:**
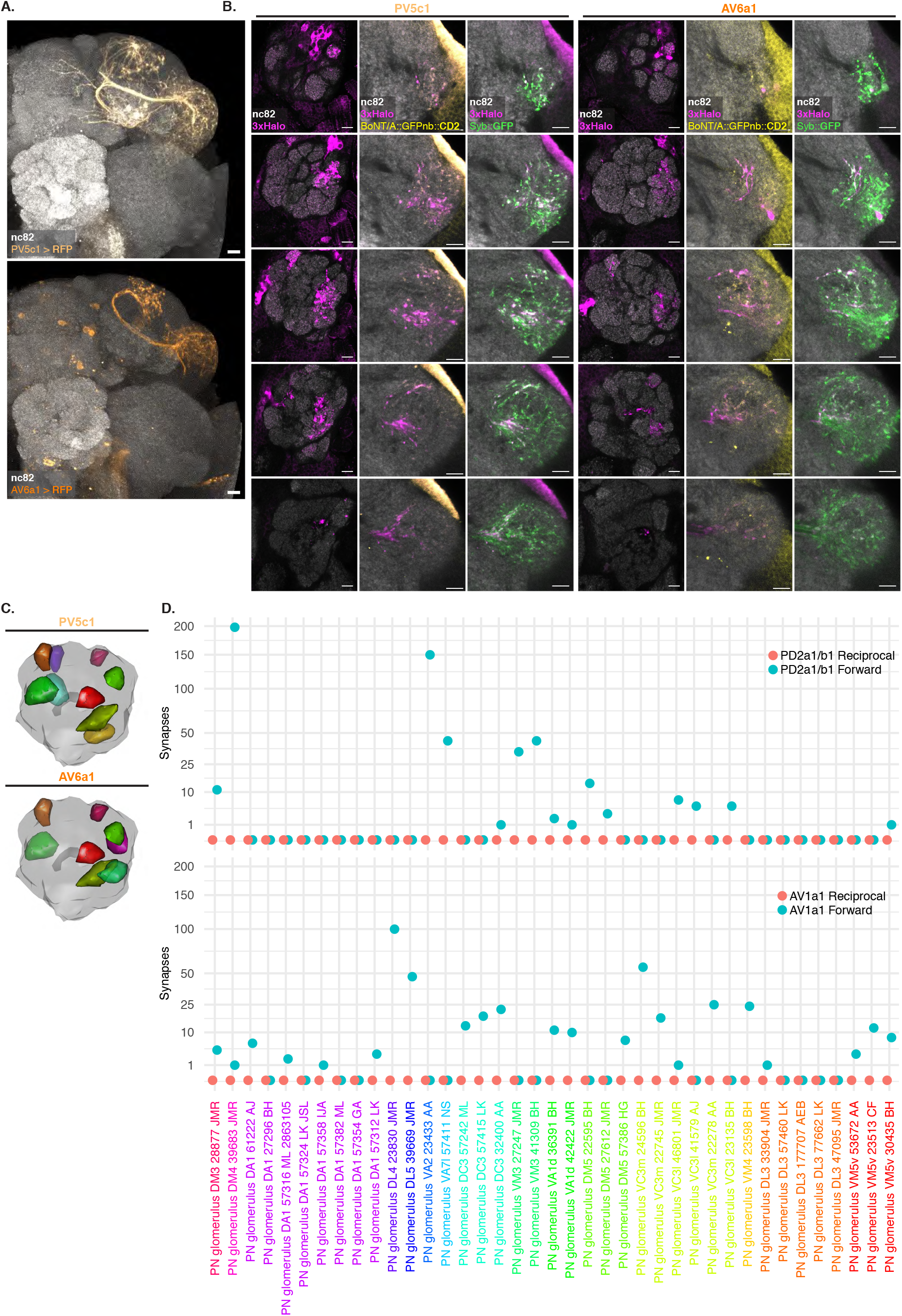
BAcTrace can reveal connections between PNs and LHNs. (**A**) Split Gal4 lines LH173 and LH1139 drive GFP expression in cell types PV5c1 and AV6a1, respectively. (**B**) Single slices from a representative antennal lobe and corresponding LH showing PNs labelled by expression of toxin in PV5c1 and AV6a1. The BAcTrace reporter Halo is shown in the AL; Halo, Syb::GFP receptor and Toxin are shown in the LH. CD2 stainings (labelling the toxin) have high background outside the neuropile. (**C**) 3D renderings summarising BAcTrace results for PV5c1 and AV6a1. (**D**) Plot of the number of forward and reciprocal synapses between PNs for the 15 glomeruli analysed in 5 and PD2a1/b1 and AV1a1 as assessed by EM. The number next to the glomerular identity of the PN is its EM identification number. Epitopes detected by antibodies are indicated in bold in (A) and (B). Scale bars (A) 50µm and (B) 10µm.

**Table S10:** Plasmids used in S2 cell experiments.

| Name | Source | Plasmid acc. |
| --- | --- | --- |
| pAWG-hTfR::Syb | This study | MN658769 |
| pAFW-hSNAP25 | This study | MN658770 |
| pAFW-QF2::V5::hSNAP25::Syx | This study | MN658771 |
| pGEX-kg-BoNT/A::GFPnb | This study | MN658772 |
| pJFRC81-BoNT/A::GFPnb::CD2 | This study | MN658773 |
| pAWH-BoNT/A-LC | This study | MN658774 |
| pAWH-TEV<sup>T173V</sup>::CD2 | This study | MN658775 |
| pAWG-mCherry::TEVs::V5::CD2 | This study | MN658793 |
| pAWG-mCherry::V5::CD2 | This study | MN658794 |
| pAWH-TEV::CD2 | This study | MN658795 |
| pAWH | [53] | - |

**Table S11:** QUAS transgenes used in this study.

| Insertion site | Transgenes | Source | Plasmid stock number |
| --- | --- | --- | --- |
| Cytological band 94E7 | QUAS-mtdTomato | Bloomington stock 30005 | - |
| attp3 | QUAS-3xHalo | This study | MN658776 |

**Table S12:**
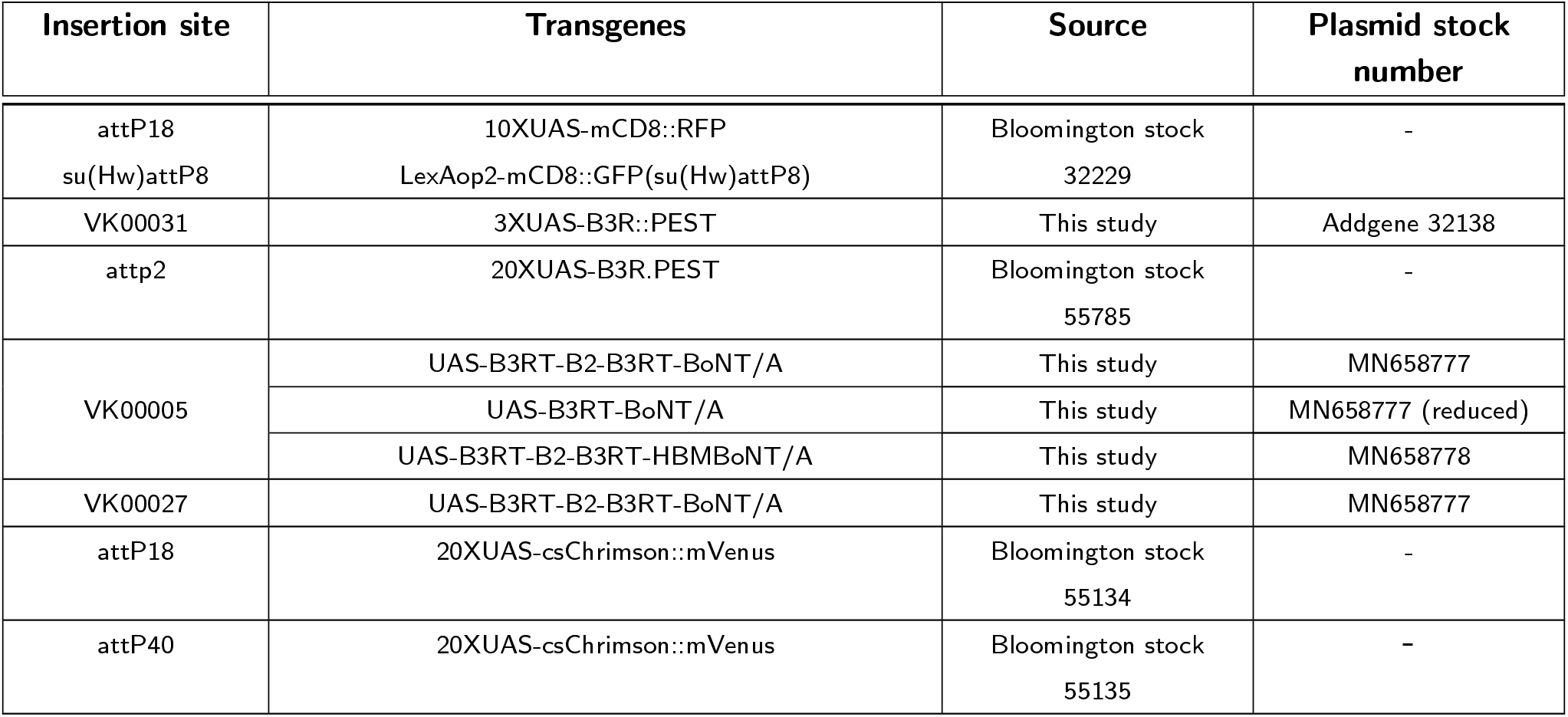
UAS transgenes used in this study.

**Table S13:** LexAop transgenes and LexA drivers used in this study. lowUTR indicates the construct does not have Syn21 or L21 5’UTRs for enhancing protein expression.

| Insertion site | Transgenes | Source | Plasmid stock number |
| --- | --- | --- | --- |
| VK00037 | LexAop2-Syb::GFP-P10 | This study | MN658779 |
| Su(Hw)attP5 | LexAop2-Syb::GFP-P10 | This study | MN658779 |
| attP40 | LexAop2-QF2::V5::hSNAP25::Syx | This study | MN658780 |
| VK00018 | LexAop2-QF2::hSNAP25::HIVNES::Syx | This study | MN658781 |
|  | LexAop2-QF2::V5::hSNAP25::Syx | This study | MN658780 |
|  | LexAop2-QF2::V5::hSNAP25::HIVNES::Syx | This study | MN658782 |
|  | LexAop2-HIVNES::QF2::V5::hSNAP25::Syx | This study | MN658783 |
|  | LexAop2-QF2::V5::hSNAP25::HIVNES::Syx-Hsp70UTR | This study | MN658784 |
|  | LexAop2-lowUTR-QF2::V5::hSNAP25::HIVNES::Syx-Hsp70UTR | This study | MN658785 |
|  | LexAop2-QF2::V5::Syntaxin | This study | MN658786 |
| Su(Hw)attP2 | LexAop2-RSRT-Flp1-RSRT-TEV::CD2-P10 | This study | MN658787 |
|  | LexAop2-RSRT-TEV::CD2-P10 | This study | MN658787 (reduced) |
|  | LexAop2-RSRT-Flp1-RSRT-TEV<sup>T173V</sup>::CD2-P10 | This study | MN658788 |
|  | LexAop2-RSRT-TEV<sup>T173V</sup>::CD2-P10 | This study | MN658788 (reduced) |
|  | LexAop2-RSRT-Flp1-RSRT-TEV<sup>3xmut</sup>::CD2-P10 | This study | MN658789 |
|  | LexAop2-RSRT-TEV<sup>3xmut</sup>::CD2-P10 | This study | MN658789 (reduced) |
| VK00033 | LexAop2-RSRT-TEV::CD2-P10 | This study | MN658787 (reduced) |
| VK00002 | LexAop2-Syb::GFP<sup>N146I</sup> | This study | MN658790 |
|  | LexAop2-Syb::GFP<sup>N146I</sup>::TEV | This study | MN658791 |
|  | LexAop2-Syb::GFP<sup>N146I</sup>::TEV<sup>T173V</sup> | This study | MN658792 |

**Table S14:** Driver lines used in this study.

| Insertion site | Line Name | Genotype | Cell types | Source |
| --- | --- | --- | --- | --- |
| JK22C |  | VT033006-LexA::p65 | Multiple PN types | Yoshinori Aso |
| NA |  | MB247-Gal4 | All KCs | Bloomington stock 50742 |
| NA |  | Orco-Gal4 | Multiple ORN types | Bloomington stock 26818 |
| NA |  | Or83c-Gal4 | Or83 ORNs | Bloomington stock 23132 |
| NA |  | Or88a-Gal4 | Or88 ORNs | Bloomington stock 23294 |
| NA |  | Or92a-Gal4 | Or92 ORNs | Bloomington stock 23139 |
| NA |  | Or47a-Gal4 | Or47 ORNs | Bloomington stock 9982 |
| NA |  | Or65a-Gal4 | Or65 ORNs | Bloomington stock 9994 |
| NA |  | Or98a-Gal4 | Or98 ORNs | Bloomington stock 23141 |
| attp40 attp2 | LH1479 | w ; 84F07-p65ADZp ; 65C07-ZpGAL4DBD | AV4b3 LHNs | http://splitgal4.janelia.org/cgi-bin/splitgal4.cgi |
| attp40 attp2 | LH1139 | w ; 93A02-p65ADZp ; 44G08-ZpGAL4DBD | AV6a1 LHNs | http://splitgal4.janelia.org/cgi-bin/splitgal4.cgi |
| attp2 | 37G11 | w ; 37G11-GAL4 | PD2a1/b1 LHNs | Bloomington stock 49539 |
| attp40 attp2 | LH196 | w ; 79C09-p65ADZp ; 31B01-ZpGAL4DBD | PD3a1 LHNs | Jefferis lab |
| attp40 attp2 | LH542 | w ; 42D01-p65ADZp ; 86A05-ZpGAL4DBD | PV4a1:5 LHNs | http://splitgal4.janelia.org/cgi-bin/splitgal4.cgi |
| attp40 attp2 | LH1554 | w ; 91F03-p65ADZp ; 71D08-ZpGAL4DBD | PV5b1 LHNs | http://splitgal4.janelia.org/cgi-bin/splitgal4.cgi |
| attp40 attp2 | LH173 | w ; 16C09-p65ADZp ; 28A10-ZpGAL4DBD | PV5c1 LHNs | http://splitgal4.janelia.org/cgi-bin/splitgal4.cgi |
| attp40 attp2 | LH719 | w ; 34C08-p65ADZp ; 17B08-ZpGAL4DBD | PV5c1 LHNs | Jefferis lab |
| attp40 attp2 | LH2033 | w ; VT043152-p65ADZp ; VT027015-ZpGAL4DBD | PV12a1 LHNs | http://splitgal4.janelia.org/cgi-bin/splitgal4.cgi |
| attp40 attp2 | LH989 | w ; 29G05-p65ADZp ; 37G11-ZpGAL4DBD | PD2a1/b1 LHNs | http://splitgal4.janelia.org/cgi-bin/splitgal4.cgi |
| attp40 attp2 | LH1983 | w; 76E07-p65ADZp ; VT008671-ZpGAL4DBD | AV1a1 LHNs | http://splitgal4.janelia.org/cgi-bin/splitgal4.cgi |
| attp40 attp2 | MB010B | w ; R13F02-p65ADZp ; R52H09-ZpGAL4DBD | KC: γmain, $\alpha'/\beta'$ap, $\alpha'/\beta'$m, $\alpha/\beta$c, $\alpha/\beta$p and $\alpha/\beta$s | Bloomington stock 68293 |
| attp40 attp2 | MB005B | w ; R13F02-p65ADZp ; R34A03-ZpGAL4DBD | KC: $\alpha'/\beta'$ap and $\alpha'/\beta'$m | Bloomington stock 68306 |
| attp40 attp2 | MB370B | w ; R13F02-p65ADZp ; R41C07-ZpGAL4DBD | KC: $\alpha'/\beta'$ap and $\alpha'/\beta'$m | Bloomington stock 68319 |
| attp40 attp2 | MB463B | w ; R35B12-p65ADZp ; R34A03-ZpGAL4DBD | KC: $\alpha'/\beta'$ap and $\alpha'/\beta'$m | Bloomington stock 68370 |
| attp40 attp2 | MB418B | w ; R26E07-p65ADZp ; R30F02-ZpGAL4DBD | KC: $\alpha'/\beta'$m | Bloomington stock 68322 |
| attp40 attp2 | MB131B | w ; R13F02-p65ADZp ; R89B01-ZpGAL4DBD | KC: γmain and γd | Bloomington stock 68165 |
| attp40 attp2 | MB419B | w ; R26E07-p65ADZp ; R39A11-ZpGAL4DBD | KC: γd | Bloomington stock 68323 |
| attp40 attp2 | MB607B | w ; R19B03-p65ADZp ; R39A11-ZpGAL4DBD | KC: γd | Bloomington stock 68256 |
| attp40 attp2 | MB008B | w; R13F02-p65ADZp ; R44E04-ZpGAL4DBD | KC: $\alpha/\beta$c, $\alpha/\beta$p and $\alpha/\beta$s | Bloomington stock 6825C91 |
| attp40 attp2 | MB185B | w ; R52H09-p65ADZp ; R18F09-ZpGAL4DBD | KC: $\alpha/\beta$s | Bloomington stock 68267 |
| attp40 attp2 | MB594B | w ; R13F02-p65ADZp ;<br>R58F02-ZpGAL4DBD | KC: $\alpha/\beta$c | Bloomington stock 68255 |

**Table S15:** Genotypes of flies used in this study. Genes in italics were present in the experiment but are not relevant to the result. All LexAop2 transgenes have 13 copies of the unitary DNA operator. QUAS constructs have 5 QUAS repeats.

| Figure | Genotype |
| --- | --- |
| 3C | w 10XUAS-mCD8::RFP(attP18) LexAop2-mCD8::GFP(su(Hw)attP8) ; VT033006-LexA::p65 / + ; MB247-Gal4 *QUAS-mtdTomato::HA* / + |
| 3D | w ; ; 20XUAS-B3R.PEST(attP2) UAS-B3RT-B2-B3RT-BoNT/A(VK00005) UAS-B3RT-B2-B3RT-BoNT/A(VK00027) / MB247-Gal4 *QUAS-mtdTomato* |
| 3E1 | w ; LexAop2-Syb::GFP-P10(VK00037) LexAop2-QF2::V5::hSNAP25::Syx(attp40) / VT033006-LexA::p65(JK22C) ; UAS-B3RT-BoNT/A(VK00005) / *QUAS-mtdTomato* |
| 3E2 | w ; LexAop2-Syb::GFP-P10(VK00037) LexAop2-QF2::V5::hSNAP25::Syx(attp40) / VT033006-LexA::p65(JK22C) ; 20XUAS-B3R.PEST(attP2) UAS-B3RT-B2-B3RT-BoNT/A(VK00005) UAS-B3RT-B2-B3RT-BoNT/A(VK00027) LexAop2-RSRT-TEV::CD2-P10(Su(Hw)attp2) / *QUAS-mtdTomato* |
| 3E3 | w ; LexAop2-Syb::GFP-P10(VK00037) LexAop2-QF2::V5::hSNAP25::Syx(attp40) / VT033006-LexA::p65(JK22C) ; UAS-B3RT-B2-B3RT-BoNT/A(VK00005) UAS-B3RT-B2-B3RT-BoNT/A(VK00027) / MB247-Gal4 *QUAS-mtdTomato* |
| 3F1 | w ; LexAop2-Syb::GFP-P10(VK00037) LexAop2-QF2::V5::hSNAP25::Syx(attp40) / VT033006-LexA::p65(JK22C) ; 20XUAS-B3R.PEST(attP2) UAS-B3RT-B2-B3RT-BoNT/A(VK00005) UAS-B3RT-B2-B3RT-BoNT/A(VK00027) LexAop2-RSRT-TEV::CD2-P10(Su(Hw)attp2) / MB247-Gal4 *QUAS-mtdTomato* |
| 3F2 | w ; LexAop2-Syb::GFP-P10(VK00037) LexAop2-QF2::V5::hSNAP25::Syx(attp40) / VT033006-LexA::p65(JK22C) ; 20XUAS-B3R.PEST(attP2) UAS-B3RT-B2-B3RT-BoNT/A(VK00005) UAS-B3RT-B2-B3RT-BoNT/A(VK00027) / MB247-Gal4 *QUAS-mtdTomato* |
| 3F3 | w ; LexAop2-QF2::V5::hSNAP25::Syx(attp40) / VT033006-LexA::p65(JK22C) ; 20XUAS-B3R.PEST(attP2) UAS-B3RT-B2-B3RT-BoNT/A(VK00005) UAS-B3RT-B2-B3RT-BoNT/A(VK00027) LexAop2-RSRT-TEV::CD2-P10(Su(Hw)attp2) / MB247-Gal4 *QUAS-mtdTomato* |
| 4B | w ; Orco-GAL4 VT033006-LexA::p65(JK22C) / LexAop2-Syb::GFP-P10(VK00037) LexAop2-QF2::hSNAP25::HIVNES::Syx(VK00018) ; *QUAS-mtdTomato::HA* / 20XUAS-B3R.PEST(attP2) UAS-B3RT-B2-B3RT-BoNT/A(VK00005) UAS-B3RT-B2-B3RT-BoNT/A(VK00027) |
| 4C DC3 | w ; LexAop2-Syb::GFP-P10(VK00037) VT033006-LexA::p65(JK22C) LexAop2-QF2::hSNAP25::HIVNES::Syx(VK00018) / CyO ; *QUAS-mtdTomato::HA* Or83c-GAL4 / 20XUAS-B3R.PEST(attP2) UAS-B3RT-B2-B3RT-BoNT/A(VK00005) UAS-B3RT-B2-B3RT-BoNT/A(VK00027) |
| 4C VA1d | w ; VT033006-LexA::p65(JK22C) Or88a-Gal4 / LexAop2-Syb::GFP-P10(VK00037) LexAop2-QF2::hSNAP25::HIVNES::Syx(VK00018) ; *QUAS-mtdTomato::HA* / 20XUAS-B3R.PEST(attP2) UAS-B3RT-B2-B3RT-BoNT/A(VK00005) UAS-B3RT-B2-B3RT-BoNT/A(VK00027) |
| 4C VA2 | w ; LexAop2-Syb::GFP-P10(VK00037) VT033006-LexA::p65(JK22C) LexAop2-QF2::hSNAP25::HIVNES::Syx(VK00018) / CyO ; *QUAS-mtdTomato::HA* Or92a-GAL4 / 20XUAS-B3R.PEST(attP2) UAS-B3RT-B2-B3RT-BoNT/A(VK00005) UAS-B3RT-B2-B3RT-BoNT/A(VK00027) |
| 4D | *QUAS-3xHalo(attp3)* w / w ; LexAop2-Syb::GFP-P10(VK00037) VT033006-LexA::p65(JK22C) LexAop2-QF2::hSNAP25::HIVNES::Syx(VK00018) / UAS-CsChrimson::mVenus(attp40) ; *QUAS-mtdTomato::HA* Or83c-GAL4 / 20XUAS-B3R.PEST(attP2) UAS-B3RT-B2-B3RT-BoNT/A(VK00005) UAS-B3RT-B2-B3RT-BoNT/A(VK00027) |
| 5A PD2a1 | w 10XUAS-mCD8::RFP(attP18) LexAop2-mCD8::GFP(su(Hw)attP8) / + ; 29G05-p65ADZp(attp40) / + ; 37G11-ZpGdbd(attp2) / + |
| 5A AV1a1 | w UAS-csChrimson::mVenus(attP18) / w ; 76E07-p65ADZp(attp40) / + ;<br>VT008671-ZpGAL4DBD(attp2) / + |
| 5C PD2a1/b1<br>S13A | QUAS-3xHalo(attp3) w / w ; LexAop2-Syb::GFP-P10(VK00037) VT033006-LexA::p65(JK22C)<br>LexAop2-QF2::hSNAP25::HIVNES::Syx(VK00018) / 29G05-p65ADZp(attp40) ;<br>20XUAS-B3R.PEST(attP2) UAS-B3RT-B2-B3RT-BoNT/A(VK00005)<br>UAS-B3RT-B2-B3RT-BoNT/A(VK00027) / 37G11-ZpGdbd(attp2) |
| 5C AV1a1<br>S13B | QUAS-3xHalo(attp3) w / w ; LexAop2-Syb::GFP-P10(VK00037) VT033006-LexA::p65(JK22C)<br>LexAop2-QF2--SNAP25-HIVNES-Syx(VK00018) / 76E07-AD(atpp40) ;<br>20XUAS-B3R.PEST(attP2) UAS-B3RT-B2-B3RT-BoNT/A(VK00005)<br>UAS-B3RT-B2-B3RT-BoNT/A(VK00027) / VT008671-DBD(attp2) |
| S5A | w ; VT033006-LexA::p65(JK22C) LexAop2-Syb::GFP-P10(Su(Hw)attp5)<br>LexAop2-QF2::V5::hSNAP25::Syx(VK00018) / CyO ; MKRS / TM6 |
| S5B | w ; LexAop2-Syb::GFP-P10(VK00037) LexAop2-QF2::V5::hSNAP25::Syx(attp40) /<br>VT033006-LexA::p65(JK22C) ; UAS-B3RT-B2-B3RT-BoNT/A(VK00005)<br>UAS-B3RT-B2-B3RT-BoNT/A(VK00027) / MB247-Gal4 QUAS-mtdTomato |
| S7A1 | w ; VT033006-LexA::p65(JK22C) LexAop2-Syb::GFP-P10(Su(Hw)attp5)<br>LexAop2-QF2::V5::hSNAP25::Syx(VK00018) / CyO ; 20XUAS-B3R.PEST(attP2)<br>UAS-B3RT-B2-B3RT-BoNT/A(VK00005) UAS-B3RT-B2-B3RT-BoNT/A(VK00027) /<br>R13F02-p65ADZp(VK00027) R34A03-ZpGAL4DBD(attP2) QUAS-mtdTomato::HA |
| S7A2 | w ; VT033006-LexA::p65(JK22C) LexAop2-Syb::GFP-P10(Su(Hw)attp5)<br>LexAop2-QF2::V5::hSNAP25::Syx(VK00018) / CyO ; 20XUAS-B3R.PEST(attP2)<br>UAS-B3RT-B2-B3RT-BoNT/A(VK00005) UAS-B3RT-B2-B3RT-BoNT/A(VK00027)<br>LexAop2-RSRT-TEV::CD2-P10(Su(Hw)attp2) / R13F02-p65ADZp(VK00027)<br>R34A03-ZpGAL4DBD(attP2) QUAS-mtdTomato::HA |
| S7A3 | w ; VT033006-LexA::p65(JK22C) LexAop2-Syb::GFP-P10(Su(Hw)attp5)<br>LexAop2-QF2::V5::hSNAP25::Syx(VK00018) / CyO ;<br>LexAop2-RSRT-TEV::CD2-P10(VK00033) 20XUAS-B3R.PEST(attP2)<br>UAS-B3RT-B2-B3RT-BoNT/A(VK00005) UAS-B3RT-B2-B3RT-BoNT/A(VK00027)<br>LexAop2-RSRT-TEV::CD2-P10(Su(Hw)attp2) / R13F02-p65ADZp(VK00027)<br>R34A03-ZpGAL4DBD(attP2) QUAS-mtdTomato::HA |
| S7A4 | w ; VT033006-LexA::p65(JK22C) LexAop2-Syb::GFP-P10(Su(Hw)attp5)<br>LexAop2-QF2::V5::hSNAP25::Syx(VK00018) / CyO ; 20XUAS-B3R.PEST(attP2)<br>UAS-B3RT-B2-B3RT-BoNT/A(VK00005) UAS-B3RT-B2-B3RT-BoNT/A(VK00027)<br>LexAop2-RSRT-TEV<sup>T173V</sup>::CD2-P10(Su(Hw)attp2) / R13F02-p65ADZp(VK00027)<br>R34A03-ZpGAL4DBD(attP2) QUAS-mtdTomato::HA |
| S7A5 | w ; VT033006-LexA::p65(JK22C) LexAop2-Syb::GFP-P10(Su(Hw)attp5)<br>LexAop2-QF2::V5::hSNAP25::Syx(VK00018) / CyO ; 20XUAS-B3R.PEST(attP2)<br>UAS-B3RT-B2-B3RT-BoNT/A(VK00005) UAS-B3RT-B2-B3RT-BoNT/A(VK00027)<br>LexAop2-RSRT-TEV<sup>3xmut</sup>::CD2-P10(Su(Hw)attp2) / R13F02-p65ADZp(VK00027)<br>R34A03-ZpGAL4DBD(attP2) QUAS-mtdTomato::HA |
| S7A6 | w ; VT033006-LexA::p65(JK22C) LexAop2-Syb::GFP-P10(Su(Hw)attp5)<br>LexAop2-QF2::V5::hSNAP25::Syx(VK00018) / CyO ; 20XUAS-B3R.PEST(attP2)<br>UAS-B3RT-B2-B3RT-BoNT/A(VK00005) UAS-B3RT-B2-B3RT-BoNT/A(VK00027)<br>LexAop2-RSRT-TEV::CD2-P10(Su(Hw)attp2) / R13F02-p65ADZp(VK00027)<br>R34A03-ZpGAL4DBD(attP2) QUAS-mtdTomato::HA |
| S7C1 | w ; LexAop2-Syb::GFP-P10(Su(Hw)attp5) LexAop2-QF2::V5::hSNAP25::Syx(VK00018) /<br>VT033006-LexA::p65(JK22C) ; UAS-B3RT-B2-B3RT-HBMBoNT/A(VK00005)<br>3XUAS-B3R::PEST(VK00031) / MB247-Gal4 QUAS-mtdTomato |
| S7C2 | w ; LexAop2-Syb::GFP<sup>N146I</sup>(VK00002) LexAop2-QF2::hSNAP25::HIVNES::Syx(VK00018) /<br>VT033006-LexA::p65(JK22C) ; UAS-B3RT-B2-B3RT-HBMBoNT/A(VK00005)<br>3XUAS-B3R::PEST(VK00031) / MB247-Gal4 QUAS-mtdTomato |
| S7C3 | w ; LexAop2-Syb::GFP<sup>N146I</sup>::TEV(VK00002)<br>LexAop2-QF2::hSNAP25::HIVNES::Syx(VK00018) / VT033006-LexA::p65(JK22C) ;<br>UAS-B3RT-B2-B3RT-HBMBoNT/A(VK00005) 3XUAS-B3R::PEST(VK00031) / MB247-Gal4<br>QUAS-mtdTomato |
| S7C4 | w ; LexAop2-Syb::GFP<sup>N146I</sup>::TEV<sup>T173V</sup>-P10(VK00002)<br>LexAop2-QF2::hSNAP25::HIVNES::Syx(VK00018) / VT033006-LexA::p65(JK22C) ;<br>UAS-B3RT-B2-B3RT-HBMB<sup>BoNT</sup>/A(VK00005) 3XUAS-B3R::PEST(VK00031) / MB247-Gal4<br>QUAS-mtdTomato |
| S8A | QUAS-3xHalo(attp3) w / w ; LexAop2-Syb::GFP-P10(VK00037) VT033006-LexA::p65(JK22C)<br>LexAop2-QF2::minSNAP25::HIVNES::Syx(VK00018) / CyO ; 20XUAS-B3R.PEST(attP2)<br>UAS-B3RT-B2-B3RT-<sup>BoNT</sup>/A(VK00005) UAS-B3RT-B2-B3RT-<sup>BoNT</sup>/A(VK00027) /<br>52H09-ZpGdbd(attP2) |
| S8B | QUAS-3xHalo(attp3) w / w ; LexAop2-Syb::GFP-P10(VK00037) VT033006-LexA::p65(JK22C)<br>LexAop2-QF2::minSNAP25::HIVNES::Syx(VK00018) / 13F02-p65ADZp(attP40) ;<br>20XUAS-B3R.PEST(attP2) UAS-B3RT-B2-B3RT-<sup>BoNT</sup>/A(VK00005)<br>UAS-B3RT-B2-B3RT-<sup>BoNT</sup>/A(VK00027) / 52H09-ZpGdbd(attP2) |
| S8C MB005B | QUAS-3xHalo(attp3) w / w ; LexAop2-Syb::GFP-P10(VK00037) VT033006-LexA::p65(JK22C)<br>LexAop2-QF2::minSNAP25::HIVNES::Syx(VK00018) / 13F02-p65ADZp(attP40) ;<br>20XUAS-B3R.PEST(attP2) UAS-B3RT-B2-B3RT-<sup>BoNT</sup>/A(VK00005)<br>UAS-B3RT-B2-B3RT-<sup>BoNT</sup>/A(VK00027) / 34A03-ZpGdbd(attP2) |
| S8C MB370B | QUAS-3xHalo(attp3) w / w ; LexAop2-Syb::GFP-P10(VK00037) VT033006-LexA::p65(JK22C)<br>LexAop2-QF2::minSNAP25::HIVNES::Syx(VK00018) / 13F02-p65ADZp(attP40) ;<br>20XUAS-B3R.PEST(attP2) UAS-B3RT-B2-B3RT-<sup>BoNT</sup>/A(VK00005)<br>UAS-B3RT-B2-B3RT-<sup>BoNT</sup>/A(VK00027) / 41C07-ZpGdbd(attP2) |
| S8C MB463B | QUAS-3xHalo(attp3) w / w ; LexAop2-Syb::GFP-P10(VK00037) VT033006-LexA::p65(JK22C)<br>LexAop2-QF2::minSNAP25::HIVNES::Syx(VK00018) / 35B12-p65ADZp(attP40) ;<br>20XUAS-B3R.PEST(attP2) UAS-B3RT-B2-B3RT-<sup>BoNT</sup>/A(VK00005)<br>UAS-B3RT-B2-B3RT-<sup>BoNT</sup>/A(VK00027) / 34A03-ZpGdbd(attP2) |
| S8C MB418B | QUAS-3xHalo(attp3) w / w ; LexAop2-Syb::GFP-P10(VK00037) VT033006-LexA::p65(JK22C)<br>LexAop2-QF2::minSNAP25::HIVNES::Syx(VK00018) / 26E07-p65ADZp(attP40) ;<br>20XUAS-B3R.PEST(attP2) UAS-B3RT-B2-B3RT-<sup>BoNT</sup>/A(VK00005)<br>UAS-B3RT-B2-B3RT-<sup>BoNT</sup>/A(VK00027) / 30F02-ZpGdbd(attP2) |
| S8D MB131B | QUAS-3xHalo(attp3) w / w ; LexAop2-Syb::GFP-P10(VK00037) VT033006-LexA::p65(JK22C)<br>LexAop2-QF2::minSNAP25::HIVNES::Syx(VK00018) / 13F02-p65ADZp(attP40) ;<br>20XUAS-B3R.PEST(attP2) UAS-B3RT-B2-B3RT-<sup>BoNT</sup>/A(VK00005)<br>UAS-B3RT-B2-B3RT-<sup>BoNT</sup>/A(VK00027) / 89B01-ZpGdbd(attP2) |
| S8D MB419B | QUAS-3xHalo(attp3) w / w ; LexAop2-Syb::GFP-P10(VK00037) VT033006-LexA::p65(JK22C)<br>LexAop2-QF2::minSNAP25::HIVNES::Syx(VK00018) / 26E07-p65ADZp(attP40) ;<br>20XUAS-B3R.PEST(attP2) UAS-B3RT-B2-B3RT-<sup>BoNT</sup>/A(VK00005)<br>UAS-B3RT-B2-B3RT-<sup>BoNT</sup>/A(VK00027) / 39A11-ZpGdbd(attP2) |
| S8D MB607B | QUAS-3xHalo(attp3) w / w ; LexAop2-Syb::GFP-P10(VK00037) VT033006-LexA::p65(JK22C)<br>LexAop2-QF2::minSNAP25::HIVNES::Syx(VK00018) / 19B03-p65ADZp(attP40) ;<br>20XUAS-B3R.PEST(attP2) UAS-B3RT-B2-B3RT-<sup>BoNT</sup>/A(VK00005)<br>UAS-B3RT-B2-B3RT-<sup>BoNT</sup>/A(VK00027) / 39A11-ZpGdbd(attP2) |
| S8E MB008B | QUAS-3xHalo(attp3) w / w ; LexAop2-Syb::GFP-P10(VK00037) VT033006-LexA::p65(JK22C)<br>LexAop2-QF2::minSNAP25::HIVNES::Syx(VK00018) / 13F02-p65ADZp(attP40) ;<br>20XUAS-B3R.PEST(attP2) UAS-B3RT-B2-B3RT-<sup>BoNT</sup>/A(VK00005)<br>UAS-B3RT-B2-B3RT-<sup>BoNT</sup>/A(VK00027) / 44E04-ZpGdbd(attP2) |
| S8E MB185B | QUAS-3xHalo(attp3) w / w ; LexAop2-Syb::GFP-P10(VK00037) VT033006-LexA::p65(JK22C)<br>LexAop2-QF2::minSNAP25::HIVNES::Syx(VK00018) / 52H09-p65ADZp(attP40) ;<br>20XUAS-B3R.PEST(attP2) UAS-B3RT-B2-B3RT-<sup>BoNT</sup>/A(VK00005)<br>UAS-B3RT-B2-B3RT-<sup>BoNT</sup>/A(VK00027) / 18F09-ZpGdbd(attP2) |
| S8E MB594B | QUAS-3xHalo(attp3) w / w ; LexAop2-Syb::GFP-P10(VK00037) VT033006-LexA::p65(JK22C)<br>LexAop2-QF2::minSNAP25::HIVNES::Syx(VK00018) / 13F02-p65ADZp(attP40) ;<br>20XUAS-B3R.PEST(attP2) UAS-B3RT-B2-B3RT-<sup>BoNT</sup>/A(VK00005)<br>UAS-B3RT-B2-B3RT-<sup>BoNT</sup>/A(VK00027) / 58F02-ZpGdbd(attP2) |
| S9A | w ; VT033006-LexA::p65(JK22C) Or88a-Gal4 / LexAop2-Syb::GFP-P10(VK00037)<br>LexAop2-QF2::minSNAP25::HIVNES::Syx(VK00018) ; 20XUAS-B3R.PEST(attP2)<br>UAS-B3RT-B2-B3RT-<sup>BoNT</sup>/A(VK00005) UAS-B3RT-B2-B3RT-<sup>BoNT</sup>/A(VK00027) /<br>QUAS-mtdTomato::HA |
| S9B | w QUAS-3xHalo(attP3) / y ; VT033006-LexA::p65(JK22C) Or88a-Gal4 / LexAop2-Syb::GFP-P10(VK00037) LexAop2-QF2::minSNAP25::HIVNES::Syx(VK00018) ; 20XUAS-B3R.PEST(attP2) UAS-B3RT-B2-B3RT-BoNT/A(VK00005) UAS-B3RT-B2-B3RT-BoNT/A(VK00027) / TM6 |
| S10A 1x Toxin | w ; VT033006-LexA::p65(JK22C) LexAop2-Syb::GFP-P10(Su(Hw)attP5) LexAop2-QF2::V5::hSNAP25::Syx(VK00018) / CyO ; 20XUAS-B3R.PEST(attP2) UAS-B3RT-B2-B3RT-BoNT/A(VK00027) / R13F02-p65ADZp(VK00027) R34A03-ZpGAL4DBD(attP2) QUAS-mtdTomato::HA |
| S10A 2x Toxin | w ; VT033006-LexA::p65(JK22C) LexAop2-Syb::GFP-P10(Su(Hw)attP5) LexAop2-QF2::V5::hSNAP25::Syx(VK00018) / CyO ; 20XUAS-B3R.PEST(attP2) UAS-B3RT-B2-B3RT-BoNT/A(VK00005) UAS-B3RT-B2-B3RT-BoNT/A(VK00027) / R13F02-p65ADZp(VK00027) R34A03-ZpGAL4DBD(attP2) QUAS-mtdTomato::HA |
| S10B No Receptor | w ; LexAop2-QF2::V5::hSNAP25::Syx / VT033006-LexA::p65(JK22C) ; LexAop2-RSRT-TEV::CD2-P10(VK00033) 20XUAS-B3R.PEST(attP2) UAS-B3RT-B2-B3RT-BoNT/A(VK00005) UAS-B3RT-B2-B3RT-BoNT/A(VK00027) LexAop2-RSRT-TEV::CD2-P10(Su(Hw)attP2) / R13F02-p65ADZp(VK00027) R34A03-ZpGAL4DBD(attP2) QUAS-mtdTomato::HA |
| S10B 1x Syb::GFP | w ; LexAop2-Syb::GFP-P10(VK00037) LexAop2-QF2::V5::hSNAP25::Syx(attP40) / VT033006-LexA::p65(JK22C) ; LexAop2-RSRT-TEV::CD2-P10(VK00033) 20XUAS-B3R.PEST(attP2) UAS-B3RT-B2-B3RT-BoNT/A(VK00005) UAS-B3RT-B2-B3RT-BoNT/A(VK00027) LexAop2-RSRT-TEV::CD2-P10(Su(Hw)attP2) / R13F02-p65ADZp(VK00027) R34A03-ZpGAL4DBD(attP2) QUAS-mtdTomato::HA |
| S10B 2x Syb::GFP | w ; LexAop2-Syb::GFP-P10(Su(Hw)attP5) LexAop2-Syb::GFP-P10(VK00037) LexAop2-QF2::V5::hSNAP25::Syx(attP40) / VT033006-LexA::p65(JK22C) ; LexAop2-RSRT-TEV::CD2-P10(VK00033) 20XUAS-B3R.PEST(attP2) UAS-B3RT-B2-B3RT-BoNT/A(VK00005) UAS-B3RT-B2-B3RT-BoNT/A(VK00027) LexAop2-RSRT-TEV::CD2-P10(Su(Hw)attP2) / R13F02-p65ADZp(VK00027) R34A03-ZpGAL4DBD(attP2) QUAS-mtdTomato::HA |
| S10C No Toxin<br>1xVT033006-LexA | w ; VT033006-LexA::p65(JK22C) LexAop2-Syb::GFP-P10(Su(Hw)attP5) LexAop2-QF2::V5::hSNAP25::Syx(VK00018) / CyO ; MB247-Gal4 QUAS-mtdTomato::HA/ TM2 |
| S10C No Toxin<br>2xVT033006-LexA | w ; VT033006-LexA::p65(JK22C) LexAop2-Syb::GFP-P10(Su(Hw)attP5) LexAop2-QF2::V5::hSNAP25::Syx(VK00018) / VT033006-LexA::p65(JK22C) ; MB247-Gal4 QUAS-mtdTomato::HA/ TM2 |
| S10C<br>1xVT033006-LexA | w ; VT033006-LexA::p65(JK22C) LexAop2-Syb::GFP-P10(Su(Hw)attP5) LexAop2-QF2::V5::hSNAP25::Syx(VK00018) / CyO ; pJFRC158-3XUAS-B3R::PEST(VK00031) UAS-B3RT-B2-B3RT-BoNT/A(VK00005) UAS-B3RT-B2-B3RT-BoNT/A(VK00027) LexAop2-RSRT-TEV<sup>T173V</sup>::CD2-P10(Su(Hw)attP2) / MB247-Gal4 QUAS-mtdTomato |
| S10C<br>2xVT033006-LexA | w ; VT033006-LexA::p65(JK22C) LexAop2-Syb::GFP-P10(Su(Hw)attP5) LexAop2-QF2::V5::hSNAP25::Syx(VK00018) / VT033006-LexA::p65(JK22C) ; pJFRC158-3XUAS-B3R::PEST(VK00031) UAS-B3RT-B2-B3RT-BoNT/A(VK00005) UAS-B3RT-B2-B3RT-BoNT/A(VK00027) LexAop2-RSRT-TEV<sup>T173V</sup>::CD2-P10(Su(Hw)attP2) / MB247-Gal4 QUAS-mtdTomato |
| S11 DM3 | QUAS-3xHalo(attP3) w / w ; LexAop2-Syb::GFP-P10(VK00037) VT033006-LexA::p65(JK22C) LexAop2-QF2::minSNAP25::HIVNES::Syx(VK00018) / CyO ; QUAS-mtdTomato::HA Or47a-GAL4 / 20XUAS-B3R.PEST(attP2) UAS-B3RT-B2-B3RT-BoNT/A(VK00005) UAS-B3RT-B2-B3RT-BoNT/A(VK00027) |
| S11 DL3 | QUAS-3xHalo(attP3) w / w ; LexAop2-Syb::GFP-P10(VK00037) VT033006-LexA::p65(JK22C) LexAop2-QF2::minSNAP25::HIVNES::Syx(VK00018) / CyO ; Or65a-Gal4 QUAS-mtdTomato::HA / 20XUAS-B3R.PEST(attP2) UAS-B3RT-B2-B3RT-BoNT/A(VK00005) UAS-B3RT-B2-B3RT-BoNT/A(VK00027) |
| S11 VM5v | QUAS-3xHalo(attP3) w / w ; LexAop2-Syb::GFP-P10(VK00037) VT033006-LexA::p65(JK22C) LexAop2-QF2::minSNAP25::HIVNES::Syx(VK00018) / CyO ; QUAS-mtdTomato::HA Or98a-Gal4 / 20XUAS-B3R.PEST(attP2) UAS-B3RT-B2-B3RT-BoNT/A(VK00005) UAS-B3RT-B2-B3RT-BoNT/A(VK00027) |
| S12 B LH1479 | QUAS-3xHalo(attp3) w / w; LexAop2-Syb::GFP-P10(VK00037) VT033006-LexA::p65(JK22C)<br>LexAop2-QF2::minSNAP25::HIVNES::Syx(VK00018) / 84F07-p65ADZp(attp40) ;<br>20XUAS-B3R.PEST(attP2) UAS-B3RT-B2-B3RT-BoNT/A(VK00005)<br>UAS-B3RT-B2-B3RT-BoNT/A(VK00027) / 65C07-ZpGdbd(attp2) |
| S12 B LH1139 | QUAS-3xHalo(attp3) w / w; LexAop2-Syb::GFP-P10(VK00037) VT033006-LexA::p65(JK22C)<br>LexAop2-QF2::minSNAP25::HIVNES::Syx(VK00018) / 93A02-Gal4AD(attp40) ;<br>20XUAS-B3R.PEST(attP2) UAS-B3RT-B2-B3RT-BoNT/A(VK00005)<br>UAS-B3RT-B2-B3RT-BoNT/A(VK00027) / 44G08-Gal4DBD(attp2) |
| S12 B 37G11 | QUAS-3xHalo(attp3) w / w; LexAop2-Syb::GFP-P10(VK00037) VT033006-LexA::p65(JK22C)<br>LexAop2-QF2::minSNAP25::HIVNES::Syx(VK00018) / + ; 20XUAS-B3R.PEST(attP2)<br>UAS-B3RT-B2-B3RT-BoNT/A(VK00005) UAS-B3RT-B2-B3RT-BoNT/A(VK00027) /<br>37G11-Gal4(attp2) |
| S12 B LH196 | QUAS-3xHalo(attp3) w / w; LexAop2-Syb::GFP-P10(VK00037) VT033006-LexA::p65(JK22C)<br>LexAop2-QF2::minSNAP25::HIVNES::Syx(VK00018) / 79C09-p65ADZp(attp40) ;<br>20XUAS-B3R.PEST(attP2) UAS-B3RT-B2-B3RT-BoNT/A(VK00005)<br>UAS-B3RT-B2-B3RT-BoNT/A(VK00027) / 31B01-ZpGdbd(attp2) |
| S12 B LH542 | QUAS-3xHalo(attp3) w / w; LexAop2-Syb::GFP-P10(VK00037) VT033006-LexA::p65(JK22C)<br>LexAop2-QF2::minSNAP25::HIVNES::Syx(VK00018) / 42D01-p65ADZp(attp40) ;<br>20XUAS-B3R.PEST(attP2) UAS-B3RT-B2-B3RT-BoNT/A(VK00005)<br>UAS-B3RT-B2-B3RT-BoNT/A(VK00027) / 86A05-ZpGdbd(attp2) |
| S12 B LH1554 | QUAS-3xHalo(attp3) w / w; LexAop2-Syb::GFP-P10(VK00037) VT033006-LexA::p65(JK22C)<br>LexAop2-QF2::minSNAP25::HIVNES::Syx(VK00018) / 91F03-p65ADZp(attp40) ;<br>20XUAS-B3R.PEST(attP2) UAS-B3RT-B2-B3RT-BoNT/A(VK00005)<br>UAS-B3RT-B2-B3RT-BoNT/A(VK00027) / 71D08-ZpGdbd(attp2) |
| S12 B LH173 | QUAS-3xHalo(attp3) w / w; LexAop2-Syb::GFP-P10(VK00037) VT033006-LexA::p65(JK22C)<br>LexAop2-QF2::minSNAP25::HIVNES::Syx(VK00018) / 16C09-p65ADZp(attp40) ;<br>20XUAS-B3R.PEST(attP2) UAS-B3RT-B2-B3RT-BoNT/A(VK00005)<br>UAS-B3RT-B2-B3RT-BoNT/A(VK00027) / 28A10-ZpGdbd(attp2) |
| S12 B LH719 | QUAS-3xHalo(attp3) w / w; LexAop2-Syb::GFP-P10(VK00037) VT033006-LexA::p65(JK22C)<br>LexAop2-QF2::minSNAP25::HIVNES::Syx(VK00018) / 34C08-p65ADZp(attp40) ;<br>20XUAS-B3R.PEST(attP2) UAS-B3RT-B2-B3RT-BoNT/A(VK00005)<br>UAS-B3RT-B2-B3RT-BoNT/A(VK00027) / 17B08-ZpGdbd(attp2) |
| S12 B LH2033 | QUAS-3xHalo(attp3) w / w; LexAop2-Syb::GFP-P10(VK00037) VT033006-LexA::p65(JK22C)<br>LexAop2-QF2::minSNAP25::HIVNES::Syx(VK00018) / VT043152-p65ADZp(attp40) ;<br>20XUAS-B3R.PEST(attP2) UAS-B3RT-B2-B3RT-BoNT/A(VK00005)<br>UAS-B3RT-B2-B3RT-BoNT/A(VK00027) / VT027015-ZpGdbd(attp2) |
| S12 B Negative control | QUAS-3xHalo(attp3) w / w; LexAop2-Syb::GFP-P10(VK00037) VT033006-LexA::p65(JK22C)<br>LexAop2-QF2::minSNAP25::HIVNES::Syx(VK00018) / 79C09-p65ADZp(attp40) ;<br>20XUAS-B3R.PEST(attP2) UAS-B3RT-B2-B3RT-BoNT/A(VK00005)<br>UAS-B3RT-B2-B3RT-BoNT/A(VK00027) / + |
| S6B1 | w ; VT033006-LexA::p65(JK22C) / LexAop2-QF2-minSNAP25-HIVNES-Syntaxin(VK00018) ;<br>MB247-Gal4 QUAS-mtdTomato::HA / + |
| S6B2 | w ; VT033006-LexA::p65(JK22C) / LexAop2-HIVNES-QF2-minSNAP25-Syntaxin(VK00018) ;<br>MB247-Gal4 QUAS-mtdTomato::HA / + |
| S6B3 | w ; VT033006-LexA::p65(JK22C) / LexAop2-QF2mSN25SyxHsp70UTR(VK00018) ;<br>MB247-Gal4 QUAS-mtdTomato / + |
| S6B4 | w ; VT033006-LexA::p65(JK22C) / LexAop2-lowUTRQF2mSN25SyxHsp70UTR(VK00018) ;<br>MB247-Gal4 QUAS-mtdTomato::HA / + |
| S6B5 | w ; VT033006-LexA::p65(JK22C) / LexAop2-QF2-Syntaxin(VK00018) ; MB247-Gal4<br>QUAS-mtdTomato::HA/+ |
| S14A PV5c1 | w 10XUAS-mCD8::RFP(attP18) LexAop2-mCD8::GFP(su(Hw)attP8) / + ;<br>16C09-p65ADZp(attp40) / + ; 28A10-ZpGdbd(attp2) / + |
| S14A AV6a1 | w 10XUAS-mCD8::RFP(attP18) LexAop2-mCD8::GFP(su(Hw)attP8) / +<br>;93A02-p65ADZp(attp40) / + ; 44G08-ZpGdbd(attp2) / + |
| S14B PV5c1 | QUAS-3xHalo(attp3) w / w ; LexAop2-Syb::GFP-P10(VK00037) VT033006-LexA::p65(JK22C)<br>LexAop2-QF2::minSNAP25::HIVNES::Syx(VK00018) / 16C09-p65ADZp(attp40) ;<br>20XUAS-B3R.PEST(attP2) UAS-B3RT-B2-B3RT-BoNT/A(VK00005)<br>UAS-B3RT-B2-B3RT-BoNT/A(VK00027) / 28A10-ZpGdbd(attp2) |
| S14B AV6a1 | QUAS-3xHalo(attp3) w / w ; LexAop2-Syb::GFP-P10(VK00037) VT033006-LexA::p65(JK22C)<br>LexAop2-QF2::minSNAP25::HIVNES::Syx(VK00018) / 93A02-p65ADZp(attp40) ;<br>20XUAS-B3R.PEST(attP2) UAS-B3RT-B2-B3RT-BoNT/A(VK00005)<br>UAS-B3RT-B2-B3RT-BoNT/A(VK00027) / 44G08-ZpGdbd(attp2) |

**Table S16:**
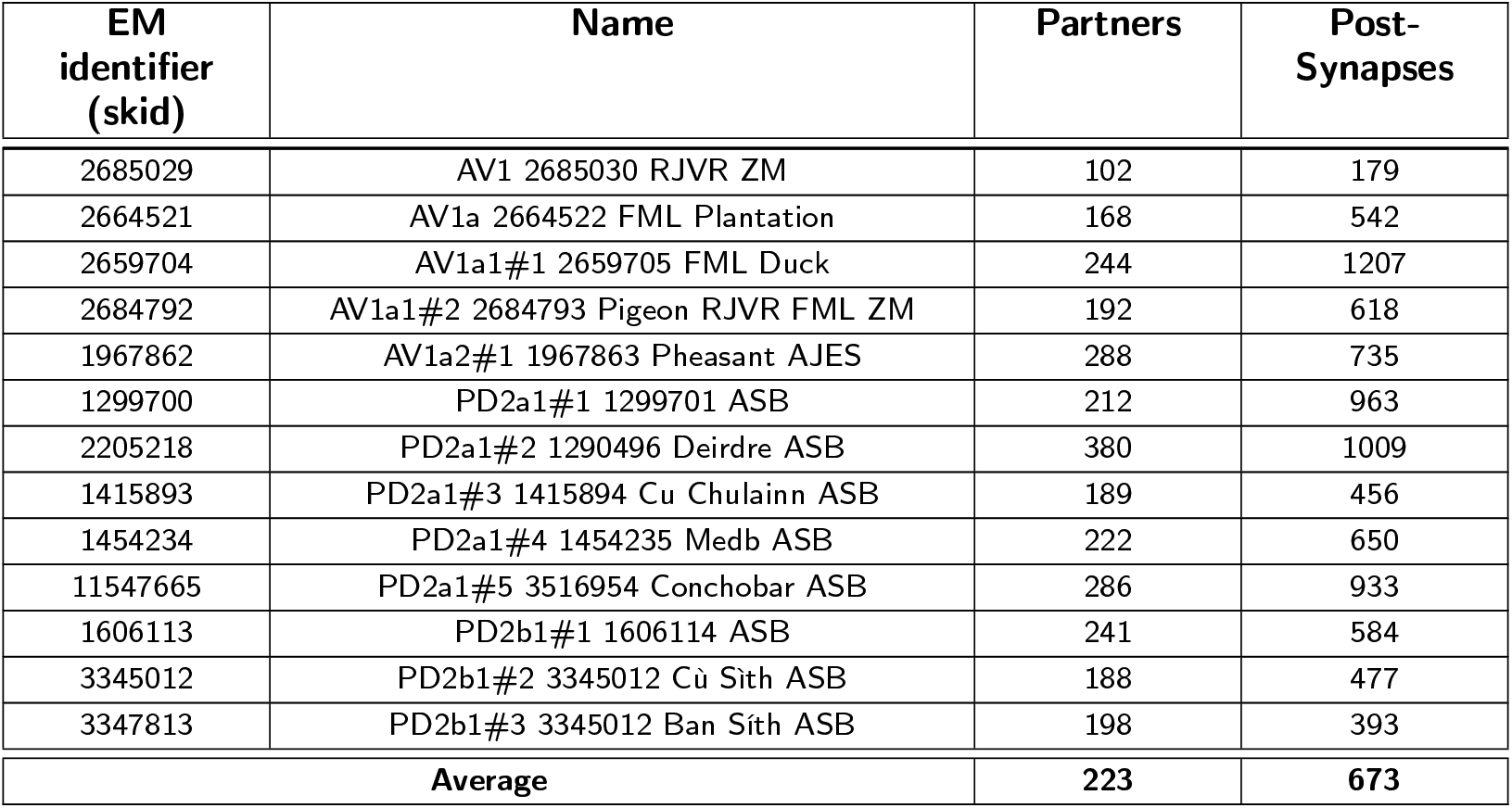
Summary of connected neurons and post-synapses of AV1a and PD2a1/b1 neurons traced on the EM volume.

| EM identifier (skid) | Name | Partners | Post-Synapses |
| --- | --- | --- | --- |
| 2685029 | AV1 2685030 RJVR ZM | 102 | 179 |
| 2664521 | AV1a 2664522 FML Plantation | 168 | 542 |
| 2659704 | AV1a1#1 2659705 FML Duck | 244 | 1207 |
| 2684792 | AV1a1#2 2684793 Pigeon RJVR FML ZM | 192 | 618 |
| 1967862 | AV1a2#1 1967863 Pheasant AJES | 288 | 735 |
| 1299700 | PD2a1#1 1299701 ASB | 212 | 963 |
| 2205218 | PD2a1#2 1290496 Deirdre ASB | 380 | 1009 |
| 1415893 | PD2a1#3 1415894 Cu Chulainn ASB | 189 | 456 |
| 1454234 | PD2a1#4 1454235 Medb ASB | 222 | 650 |
| 11547665 | PD2a1#5 3516954 Conchobar ASB | 286 | 933 |
| 1606113 | PD2b1#1 1606114 ASB | 241 | 584 |
| 3345012 | PD2b1#2 3345012 Cù Sith ASB | 188 | 477 |
| 3347813 | PD2b1#3 3345012 Ban Sith ASB | 198 | 393 |
| Average |  | 223 | 673 |

